# Temporal coding carries more stable cortical visual representations than firing rate over time

**DOI:** 10.1101/2025.05.13.652528

**Authors:** Hanlin Zhu, Fei He, Pavlo Zolotavin, Saumil Patel, Andreas S. Tolias, Lan Luan, Chong Xie

**Affiliations:** Department of Electrical and Computer Engineering, Rice University, Houston, TX, USA; NeuroEngineering Initiative, Rice University, Houston, TX, USA; Department of Ophthalmology, Stanford University, Stanford, CA, USA; Stanford BioX, Stanford University, Stanford, CA, USA; Wu Tsai Neurosciences Institute, Stanford University, Stanford, CA, USA; Department of Electrical Engineering, Stanford University, Stanford, CA, USA; Department of Bioengineering, Rice University, Houston, TX, USA

## Abstract

The brain’s ability to stably represent recurring visual scenes is crucial for behavior. Previous studies have used slow dynamic (1-5 seconds) rate code measurements to study visual tuning, revealing varying degrees of gradual activity changes over time or “representational drifts.” However, it remains unclear if there is an underlying neural code that maintains the encoding of information stable over time. In this study, we extracted structures in fast (tens of milliseconds) temporal responses and explored the role of such temporal codes in supporting the stability of visual representations. We tracked the spiking activity of the same visual cortical populations in male mice for 15 consecutive days using custom-developed, large-scale, ultraflexible electrode arrays. Across various types of stimuli, we found that neurons exhibited varying degrees of day-to-day stability in their firing rate-based tuning. The across day stability correlated with tuning reliability. Notably, accounting for spiking temporal dynamics increased single neuron tuning stability, especially for less reliable neurons. Temporal coding further improved population representation discriminability and decoding accuracy. The stability of temporal codes was more correlated with network functional connectivity than rate coding. These results show that temporal coding is crucial for stably encoding sensory stimuli, suggesting its significant role in ensuring consistent sensory experiences.

## Introduction

The plasticity in neuronal responses^1, 2^ allows us to adapt behaviorally to the ever-changing environment. Yet, even without overt adaptation pressure, spontaneous remapping of neural activity, known as drifts, has been widely reported in multiple brain regions^3–6^. In the visual cortex, prior studies have revealed various degrees of drifting in response to diverse stimulation patterns. While single neuron tuning to orientation^7^, spatial frequency^8^, size^9^ of artificial grating stimuli is moderately consistent over the course of weeks, naturalistic stimuli evoke responses that drift considerably over the course of minutes to weeks^10^, occurring in a large number of visual sub-regions and neuron types^10, 11^. In recent literature^12^. it is observed that while responses to 30 seconds-long natural movie drifted across weeks at the level of individual neurons, a populational neural code denoised by unsupervised machine learning provided a more stable representation of time within the movie clip. However, it remains unclear what neural mechanisms generally anchor consistent visual perception at sub-second timescale and are relevant to both individual neurons and neural populations. Importantly, prior studies predominantly rely on two-photon imaging of Ca^2+^ transients to reveal visually evoked population responses, which only provide a proxy of spiking with limited temporal resolution^13–17^. Consequently, these studies mostly use seconds-long rate codes (firing rates, spike counts normalized by elapsed time) to compute visual representations, largely eliminating structures and information in the fast spiking dynamics. Information and structures in the timing of stimulus-evoked spikes, referred to as the temporal code^18^, may describe unique aspects of external stimuli. We thus hypothesize that temporal code and rate code have different rates of representational drift over extended periods.

Temporal code is supported by both anatomical and functional evidence. Visual cortical neurons receive diverse types of synaptic inputs, ranging from feedforward retinal input and between-layer recurrent connections to feedback connections from higher visual areas^19, 20^, which naturally give rise to varying temporal dynamics. These asynchronous inputs and their complicated interactions could form the basis of a temporal code^21^. Accordingly, spiking dynamics responding to sustained visual scenes have been hypothesized to code for contrast, spatial frequencies, and figure-ground differentiation^22–26^. These snapshot studies, however, have not determined whether temporal dynamics remain consistent over days or how they contribute to the stability of visual representations. In addition, only a handful of neurons are typically recorded, thus limiting the study of population representation and visual decoding. Recently, high-density silicon probes have enabled chronic^27^, large-scale^28^ recording in mouse visual cortices, yet their ability to study longitudinal representation stability has not been clearly established.

Here we report a time-resolved evaluation of long-term visual representations in the mouse visual cortex using stable, large-scale electrophysiological recordings by ultraflexible nanoelectronic threads^29, 30^ (NETs). NETs, being ultraflexible and only 1 μm thick, form seamless, glial scar-free integration with the surrounding tissue, largely eliminating instability at the tissue-electrode interface in both the short and long terms that could otherwise confound the study of the representation stability. We tracked the same neuronal populations over the course of 15 consecutive days while mice experienced diverse, repeated stimuli including drifting gratings, static gratings, receptive field Gabors and natural scenes. Neurons displayed varying levels of day-to-day stability in their rate code tuning. This stability correlated with their tuning reliability. We found that visually evoked spiking time courses could largely represent stimulus features over time consistently when the readout time bins were fixed. Critically, taking into account temporal codes with fast dynamics (tens of milliseconds), compared with rate codes (hundreds of milliseconds), improved the stability of single neuron tuning, especially for less reliable neurons. At the populational level, temporal coding led to more stable and distinct representations in all tested stimuli. Accordingly, adding temporal information consistently improved the stimulus decoding accuracy in future days. Lastly, we found that the stability of temporal code was associated with network functional connectivity and its changes. Collectively, our time-resolved, day-to-day, multi-scale analysis of visual representation stability, facilitated by large-scale recordings from ultraflexible electrode arrays, has shown that temporal coding plays a crucial role in ensuring consistent visual perception.

## Results

### Longitudinal tracking of hundreds of single units under diverse visual stimuli reveals tuning stability and drifts

Understanding visual representations and their longitudinal stability requires recording at multiple scales, encompassing individual cells to large neuronal populations. This approach is essential because instabilities may exist at the level of individual units while stability may emerge at the population level^7^. Additionally, various factors such as inconsistent familiarity with the stimuli and instability at the tissue-electrode interface could potentially mask or contaminate the intrinsic stability of the neuronal responses and thus need to be carefully controlled. To fulfill these requirements, we implanted a total of 25 32-channel NETs in the visual area (V1 and LM) of n = 5 mice (3-7 NETs in each animal) and recorded neuronal populations during visual stimuli over 15 consecutive daily sessions. We performed these recordings 50 days post-surgery to allow ascular and neuronal remodeling to cease sufficiently^31^, followed by a familiarization period of 12 days (median) to remove the confounds of stimulus novelty on neural coding^5, 32, 33^. In each session, we presented identical visual stimuli of four major types (Fig. 1a): drifting gratings, static gratings, receptive field Gabors, and natural images.

**Fig. 1:**
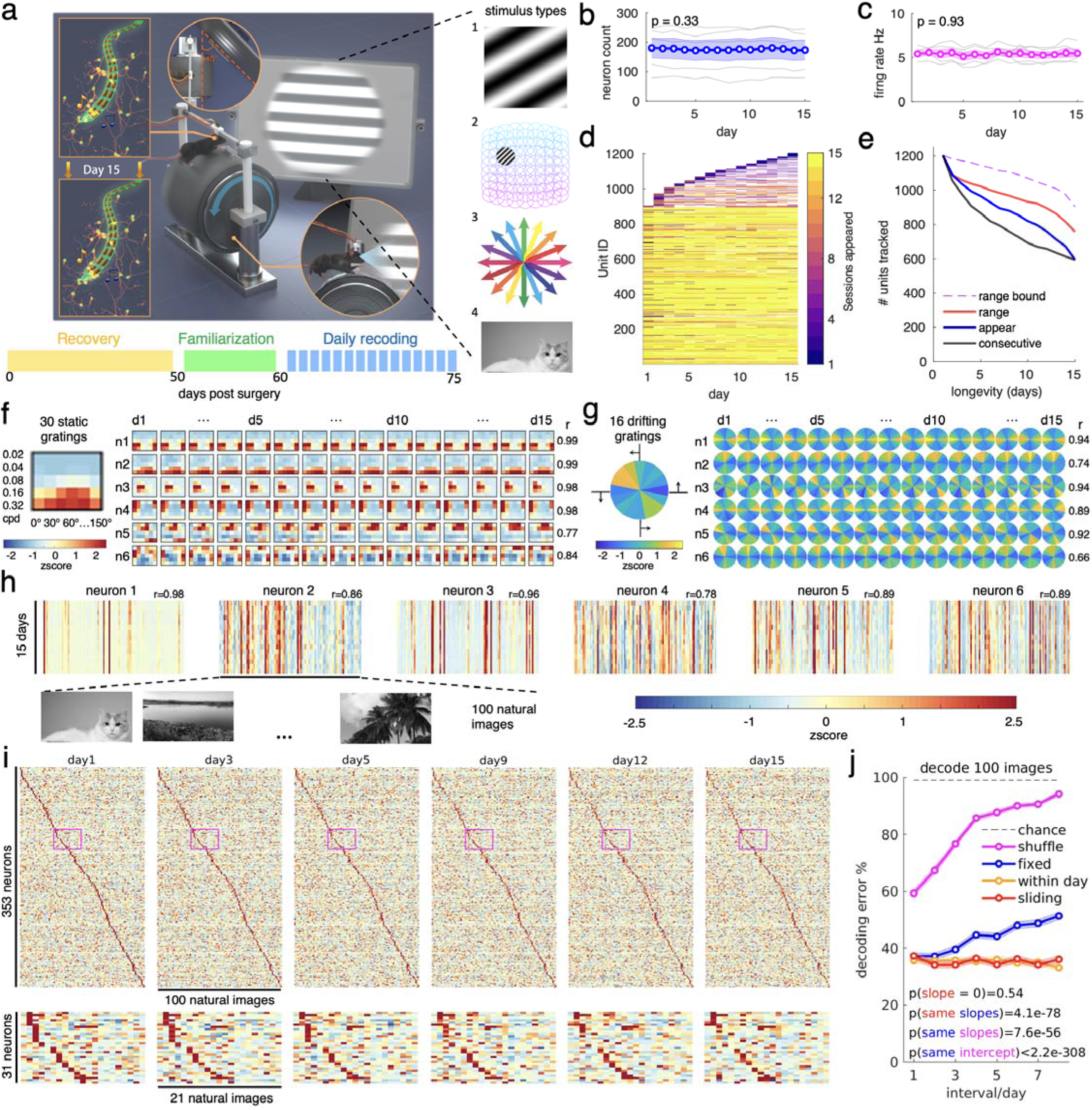
Longitudinal tracking of hundreds of single units under diverse visual stimuli reveals tuning stability and drift. **a**. Schematics (left) showing the measurement setup, Credit: Wang Wu, Shanghai. Ultraflexible NETs were implanted in V1 and LM for stable, tissue-integrated recordings under diverse stimuli (right). Bottom: experimental timeline of an example animal. **b-c**. The number of single units (b) and their average firing rates (c) from 5 animals (gray lines) recorded over 15 days did not change. Linear mixed effect, time as fixed effect, individual animal as random effect. Colored lines indicate mean ± s.e.m. **d**. A large fraction of single units tracked for extended periods, showing longevity tally, units sorted by first appearance session and color-coded for tracking longevity. **e**. Number of tracked single units for certain intervals based on 3 criteria: total number of sessions appeared (appear, blue), consecutive sessions appeared (consecutive, black), and the day range between the first and last seen session (range, red). The purple dashed line indicates the maximal possible range for a given unit (e.g., a unit first appearing on day 14 can only be tracked for a maximum day range of two days). See Supplemental Methods for more quantification explanations. **f-h**. Stable functional tuning to visual stimuli across all 15 days. Color denotes z-scored firing rate for 6 example neurons responding to 30 static gratings (f) of 5 spatial frequencies in cycles per degree (cpd) and 6 orientations; 6 example neurons responding to 16 drifting gratings with grating directions spaced 22.5°, with example 4 directions marked (g); and 6 example neurons responding to 100 natural images. Natural image examples from the authors, actual stimuli not shown due to copyright restrictions. (h). Tuning curve similarity after 7 days is denoted by the correlation coefficient “r” value. **i**. A large number of neurons (n = 353) had diverse, distinctive, and overall stable preferences for 100 natural images across 15 days, neuron orders were not resorted between days. Bottom: zoomed-in plot of pink-marked regions in the top panel, showing evidence of minor but clear feature preference drift across time in neuron populations. **j**. Neuron identity shuffled decoder approached chance at a higher rate than the fixed-in-time decoder when predicting 100 natural images. 10% of cells were shuffled per session, resulting in an expected (0.9)^day^ fraction of unshuffled neurons at any given day. The sliding-in-time decoders further reduced the rate of error increase compared to the fixed decoders for all stimuli. Sliding decoders did not show an error increase and approached within-day decoding performance. Colored lines indicate mean ± s.e.m. Linear mixed effect, time as fixed effect, decoding method (shuffle vs. fixed-in-time / sliding vs. fixed-in-time) as fixed effect, 500 animal-image pairs as random effect. Only tuned single neurons appearing for 15 days were included in the decoder of any given animal. See Supplementary Table 1 for additional reporting on sample size and statistics.

To verify stability of longitudinal recording, we quantified the number of recorded single units (Fig. 1b), their mean firing rates at each session (Fig. 1c) as well as the spatial distribution of unit amplitude and firing rate along cortical depth (Supplementary Fig1a-b). All these metrics were stable over the entire longitudinal period, confirming minimum electrode-tissue movements and permitting tracking of individual neurons throughout all sessions. We used an actively-learned ensemble of hierarchical clustering trees^34^ to match neurons across all sessions and detected a total of 1204 tracked neurons (Fig. 1d). They appeared for an average of 10.96 ± 0.15 days (mean + s.e.m.). Among which 118.80 ± 26.97 neurons per mouse showed up on all 15 days (mean ± s.e.m.). We cross-validated this process of matching units across days with other methods including dense neural network and discriminant analysis^35^ and found comparable results (Methods, Supplemental Discussion, Supplementary Methods). We tracked a large portion of units for an extended amount of time, for example, among the 903 neurons that first appeared on day 1, 594 (66%) of them appeared for all 15 days (Fig. 1e). Tracked units showed stable unit amplitude and session averaged firing rate across 15 days (Supplementary Fig1c-e). Furthermore, we computed the distribution of waveforms similarity and estimated position variations of same tracked units across days, against that of other units within day, confirming that the tracked units had much smaller changes in waveforms and locations (Supplementary Fig1f-g). These high-fidelity single neurons tracking represented substantially improved scale and longevity compared with prior chronic electrophysiological studies in rodent visual cortex^1, 36^, enabling us to examine large-scale neural functional stability over time at high temporal resolutions.

To study the functional stability of tracked neurons, we first computed their rate code tuning for all four stimulus types and observed both marked stability and noticeable drift. Large portions of neurons were significantly tuned to each stimulus (56 ± 3%, 61 ± 6%, 55 ± 4%, 83 ± 4% for drifting gratings, static gratings, receptive field Gabors, and natural images respectively on day 1), and the tuned portions remained unchanged over time (P > 0.26), indicating overall functional stability at the population scale (Supplementary Fig2a-b). Furthermore, many neurons significantly tuned to stimuli exhibited stable tuning across days, defined as having a trial-averaged tuning curve highly similar (e.g. correlation coefficient greater than 0.9) to itself after 7 days. This holds true for both low-dimensional stimuli such as static gratings (n = 533, 23% stable; Fig. 1f), drifting gratings (n = 481, 24% stable; Fig. 1g, Supplementary Fig2c), receptive field Gabors (n = 455, 9% stable; Supplementary Fig2d), and high-dimensional stimuli like natural images (n = 684, 13% stable; Fig. 1h). Across a population of 353 neurons that appeared and significantly tuned to natural images on all days, we observed visually consistent day-to-day most preferred natural images across 15 days (Fig. 1i, diagonal line). Conversely, we also detected longitudinal drifts in the tuning, including some units losing their tuning to the most preferred images (Fig. 1i, zoomed-in view). Consistent with these day-to-day changes in tuning, the error rate of a fixed-in-time population decoder (linear discriminant analysis, LDA) trained with the first 7 days increased with the time interval when predicting natural images of future days. However, applying a sliding-in-time decoder (trained with past 7 days and decoded the next day) removed this drift (P > 0.21), indicating that the day-to-day changes were gradual and trackable (Fig. 1j). Importantly, faithful mapping of visual representational drift required reliable tracking of neuron identity across days: shuffling 10% of neuron identity within mice per day as a simulation of unstable recording eroded the decoder performance significantly towards chance level (Fig. 1j). We observed similar decoding error indexed stability and drift for all other stimulus types (DG,SG,RFG stimuli. Supplementary Fig3).

### Neurons have varying representational stability which strongly correlates with tuning reliability

What impacts the stability of neural representation? To answer this question, we delved into the firing rate tuning of individual neurons and quantified the similarity between a single neuron’s tuning on one day (day x, x = 1 to 7) and 7 days later (day x+7). The tuning similarity across all four stimulus types, defined as the Pearson’s correlation coefficient between the tuning curves, displayed a wide range of variation, as depicted in Fig. 2a. To decipher the contributing factors to the stability, we related the across-day tuning similarity to the unit’s initial tuning metrics such as tuning reliability^10, 37^ (trial-to-trial variations), tuning strength^27^ (test statistics of unequal response to different stimuli across trials), and tuning selectivity^38^ (inequality coefficient of trial-averaged response to different stimuli). We found that tuning reliability (r > 0.8), tuning strength (r > 0.7), and tuning selectivity (r > 0.3) predicted long-term tuning stability (Fig. 2b, d, Supplementary Fig4a, c). In comparison, putative cell types^39^ (pyramidal vs interneuron) did not meaningfully differed in tuning stability (Fig. 2b, d, 4^th^ panel. Supplementary Fig4a, c). Finally, these contributors could also jointly explain tuning curve similarity across days (Fig. 2c, e, Supplementary Fig4b, d). Equally important, unit physical location changes played a minor role, suggesting that the observed tuning drift cannot be trivially explained by the possible existence of tracking errors (Supplementary Fig4e). These results indicate that neurons exhibit varying stability both within-session (reliability) and across daily sessions. A small portion of neurons responded highly reliably and stably to visual stimuli despite noise^40^ and mixed coding^41^ in the cerebral cortex. These units might serve as anchors^42^ for stable visual representations.

**Fig. 2:**
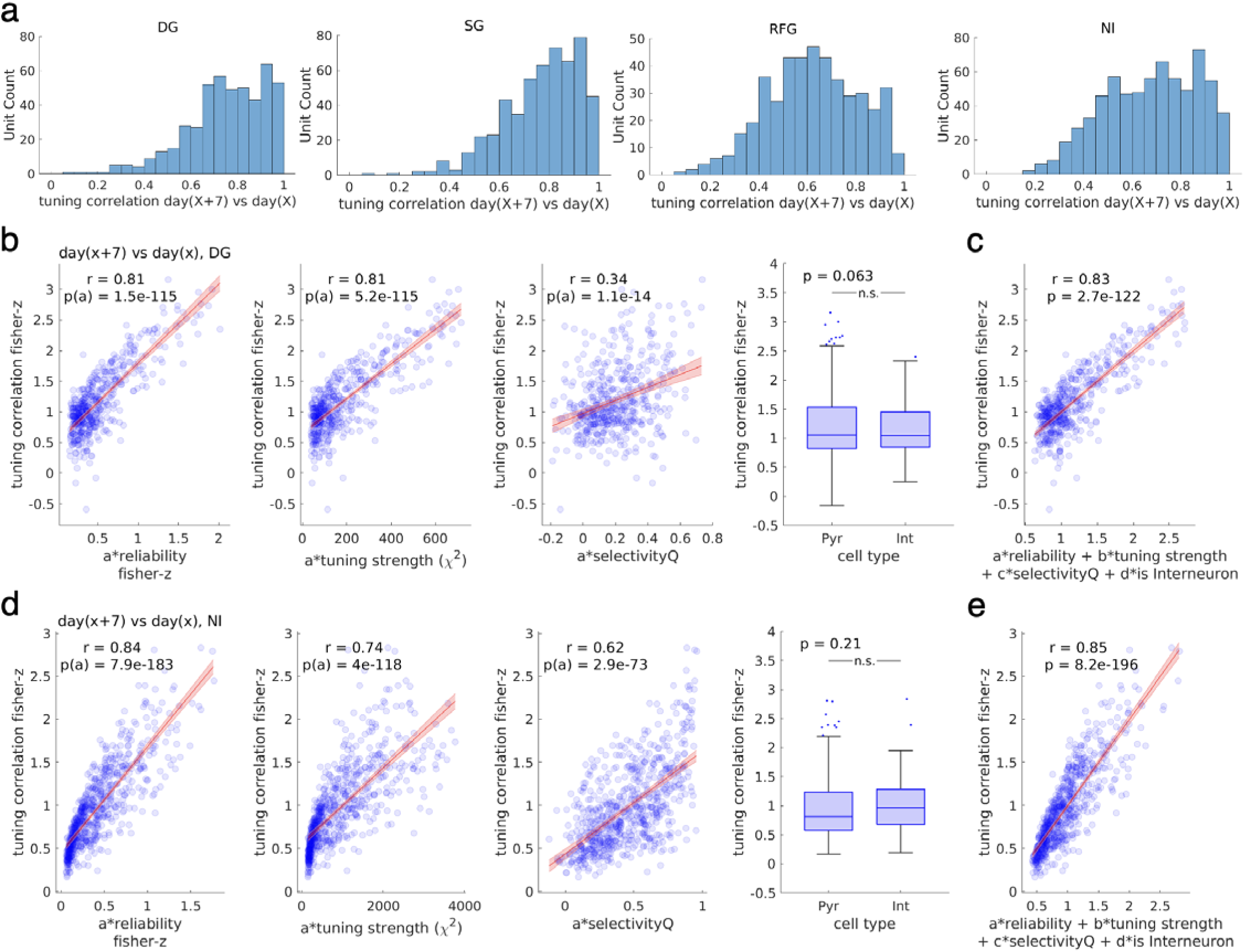
Neurons have varying representational stability which strongly correlates with tuning reliability. **a**. Across all four stimulus types, a large number of units had trial-averaged tuning curves that resembled themselves after 7 days, using stimulus-tuned single units appearing > 2 days and reappeared after 7 days. Each neuron contributed 1 data point: averaged of all possible instances tuning curve similarity after 7 days interval. N = 481, 533, 455, 684 for the 4 stimuli DG,SG,RFG,NI respectively. DG: drifting gratings; SG: static gratings; RFG: receptive field Gabors; NI: natural images. **b**. Tuning similarity after 7 days during drifting gratings was strongly correlated with tuning reliability (1^st^ column), moderately correlated with tuning strength (the Kruskal-Wallis test statistics, 2^nd^ column), and stimulus selectivity (3^rd^ column), while being weakly correlated with cell type (independent t-test, 4^th^ column, 300 pyramidal neurons, 181 interneurons). N=481 neurons. Correlation coefficient was Fisher Z-transformed. Red line: linear regression fit with 95% CI, text: correlation coefficient, significance level. **c**. Reliability, tuning strength, selectivity, and cell types could jointly explain tuning similarity after 7 days. Red line: linear regression fits with 95% CI, text: correlation coefficient, significance level. **d-e**. As in (b-c), but for natural image stimuli. N=684 neurons (451 pyramidal neurons, 233 interneurons). Box plots are in median, 25 to 75 percentiles. Whiskers represents 1.5-fold interquartile range below Q1 or above Q3. Outliers are indicated in scatters. See Supplementary Table 1 for additional reporting on sample size and statistics.

### Temporally resolved populational representation exhibits stable latency between stimulus categories

Our quantification of the stability of visual representations thus far, like previous studies^7, 9–11, 43^, relied on the average spiking rate over the entire stimulus presentation duration (stimulus duration: typically 1-5 seconds in past studies, 200-500 ms depending on the stimulus type in this work). However, we recognize that this approach overlooks the role of temporal patterns in the spiking activities, which may contain additional stimulus information^24, 26^. As a result, the stability of the neural code representing sensory information at higher temporal resolution (e.g., tens of milliseconds) remains unknown. Therefore, we studied the fine temporal resolution of electrophysiological recordings to explore the chronic stability of the temporal patterns of neural firing (Fig. 3) and how these patterns contributed to enhancing stimulus coding reliability and stability (Fig. 4).

**Fig. 3:**
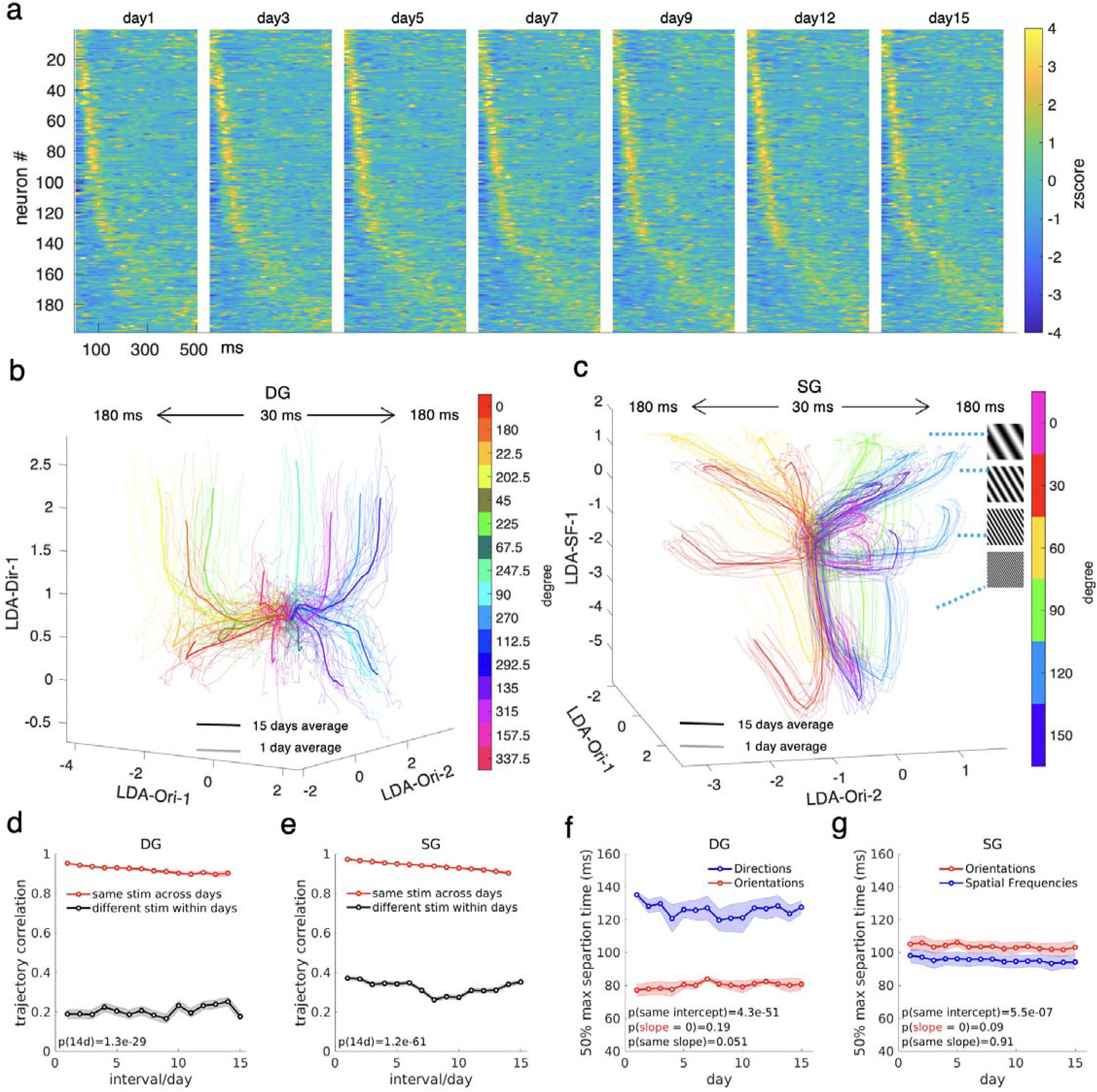
Temporally resolved populational representation exhibits stable latency between stimulus categories. **a.** The stimulus-evoked spiking time course to a single, specific stimulus (i.e. 0° drifting grating) of a population of neurons across multiple days. Neurons exhibited diverse temporal response profiles to this stimulus. Neurons were pooled from 5 mice over 7 different experiment days. Neurons that appeared for all 15 days and tuned to drifting gratings were selected. The neurons were sorted by maximum activation time and were not resorted between days. **b**. Disentangled population representation dynamics during drifting gratings from an example animal, showing daily trial-averaged (lighter lines) and 15-day-averaged (darker lines) traces, projected onto the first two linear discriminant orientation coding dimensions and the first direction coding axis. Moving angles of gratings having the same orientation but different directions (180° apart) were coded in similar colors. **c**. Same as in (a) but for static grating stimuli, projected onto the first two orientation coding dimensions and the first spatial frequency coding axis. Stimulus orientations were coded in color. **d**. The dynamical representations of same drifting grating stimuli (daily averaged trajectories in b) were more similar to themselves even after 14 days than other stimuli fitted from the same animals within days as measured by the correlation coefficient, lines are in mean ± s.e.m., independent t-test on 14 days interval, n=80, 9000 same/different mouse-pattern pairs respectively. **e**. Same as in (d) but for static gratings stimuli in (c), n=120, 20700 same/different mouse-pattern pairs respectively. **f**. The timing at which the mean inter-stimulus category distance reached 50% max for 5 mice across 15 days, all lines showing mean ± s.e.m. Orientations were consistently separated sooner than directions during the presentation of drifting grating stimuli across 15 days, linear mixed effect, time, stimulus category as fixed effect, individual animals (n=5) as random effect. **g**. Similar to (f) but during static grating stimuli. Spatial frequencies were separated sooner than orientations during static grating and were consistent across 15 days. See Supplementary Table 1 for additional reporting on sample size and statistics.

**Fig. 4:**
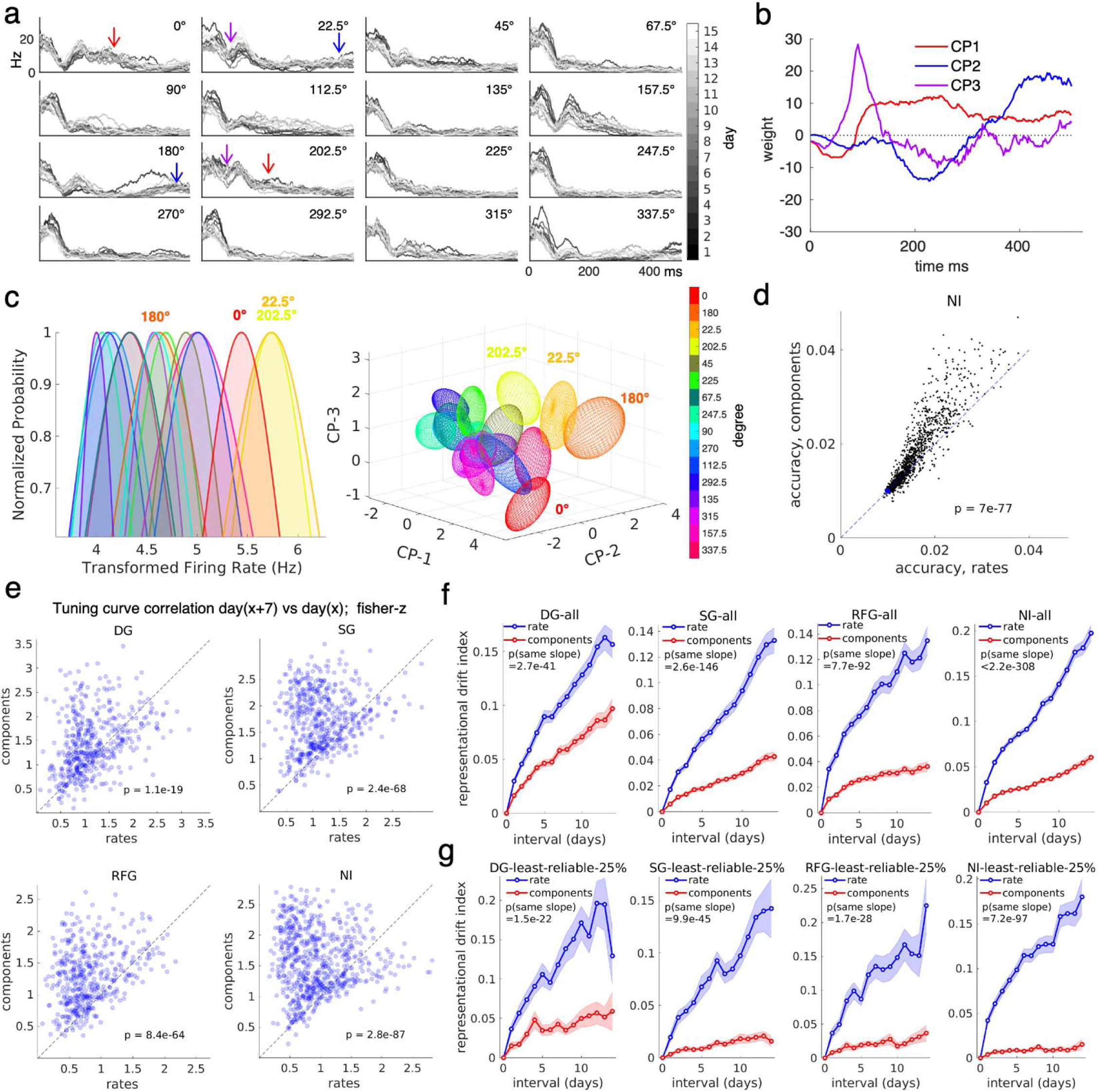
The temporal dynamics of individual neurons enhance tuning stability. **a**. Daily averaged, smoothed spiking time courses for one neuron during the stimulus-on period in response to different directions of drifting grating stimuli, instantaneous rate in Hz. Arrow coloring and implications: see (b) and main text. **b**. The first 3 LDA temporal components extracted for this unit showed a diverse tempor l profile that matches intuitions for stimulus separation with temporal dynamics in (a). **c**. Components provided greater separation between stimuli compared to firing rates. This is illustrated by an example neuron’s daily trial-averaged firing rates (left) and the distribution of the first three components (right) across 15 days under 16 different stimuli. The distributions (only showing 1 standard deviation region/ellipsoid) highly overlapped for firing rates (Left, Anscombe transformed), components-based representation reduced the overlap between highly similar stimuli (Right). **d**. Components increased decoding accuracy for nearly all single units (n=1204) during natural image stimuli, showing cross-validation decoding accuracy for all trials pooled from 15 days. Chance = 1/100 or 0.01. Paired t-test. **e**. Across all stimuli, temporal components-based unit tuning curves were more similar to themselves after 7 days than that of firing rate-based. Dots represent neurons in the population, n = 481, 533, 455, 684 stimulus-tuned single units that reappeared after 7 days for DG,SG,RFG,NI respectively. Paired t-test. Correlation coefficient was Fisher Z-transformed. **f.** Tuning curves across all 4 stimulus types drifted less over time when defined with the first 3 temporal components than when defined with firing rates. For each recording day and for each neuron, 50% of trials were randomly selected and averaged to form the tuning curve for within- and between-session similarity comparisons. Sampling was repeated 30 times, and the results were further averaged. All lines show the mean ± s.e.m. Linear mixed effect: time, definition method as fixed effect, tuned single units from all mice as random effect. DG: drifting gratings; SG: static gratings; RFG: receptive field Gabors; NI: natural images. N = 699, 719, 665, 831 stimulus-tuned single units for DG,SG,RFG,NI respectively. **g**. same as in Fig4f but only selecting 25% least reliable neurons in Fig4e for each stimulus type separately. Temporal components showed less drift for these neurons with a seemingly larger effect size. Firing rate-based reliability was used to shortlist neurons. See Supplementary Table 1 for additional reporting on sample size and statistics.

Is the millisecond dynamics of the populational representations stable over time? To answer this question, we extracted single-trial population dynamics from all 15 sessions. Responses to drifting gratings (Fig. 3a) and natural images (Supplementary Fig5a) appeared stable across two weeks. Neurons responding to a given stimulus were diverse and visually stable across all longitudinal sessions. To quantify the stability of the population dynamics that encode individual stimulus categories (e.g., 24 static gratings can be grouped into either 6 orientations or 4 spatial frequencies), we projected the population firing time course of each animal into a low-dimensional stimulus coding space with cross-validated LDA decoders^44^, pooling all trials across 15 days. We built the linear decoder based on rate code at high temporal resolution, i.e., firing rates computed between 70 and 160 ms. This temporal window accounted for the time delay of visual signal propagation and ensured different stimulus categories were readily decodable. The fixed decoder weight then projected neural data from 30 ms to 180 ms in 10 ms increments to evaluate representation differentiation at fine temporal resolution. In this coding space, nearby locations intuitively represented similar stimuli; the temporal evolution of stimuli-evoked population dynamics and their day-to-day stability were evident. The representations of the eight orientations of drifting gratings stimuli resembled an octagon formation, starting at the center and propagating radially outward from 30 ms to 180 ms (Fig. 3b, Supplementary Fig6a for breakdown plots of individual components, and Supplementary Movie 1-4). Notably, at approximately 90 ms, the two gratings having the same orientation but opposite drifting directions (e.g., 0° vs 180°, coded by similar color) bifurcated vertically, making these initially overlapping representations clearly separable (Supplementary Fig7). Similarly, the four spatial frequencies of static gratings stimuli were established in four parallel planes, where the six orientations evolved radially outward forming a hexagonal formation (Fig. 3c, Supplementary Fig6b, Supplementary Fig7, Supplementary Movie 5-7). Furthermore, the population dynamics representing the same stimulus were similar across days. For example, the temporal trajectory of the 24 static gratings across the 15 days formed 24 separable bundles, each of 15 traces (Fig. 3c). Statistically, trajectories of the same drifting gratings stimuli (Fig. 3d) overlapped more closely (P < 1.3e-29, independent t-test) with themselves (r = 0.90 ± 0.01) even after 14 days than with trajectories of different stimuli (r = 0.20 ± 0.01) recorded simultaneously. Similar results were obtained for static gratings (Fig. 3e).

Resolving the dynamics of populational representation allowed us to determine the exact timing and sequence of separation for coexisting stimulus feature categories (i.e., orientations versus directions for drifting gratings, orientations versus spatial frequencies for static gratings) and, importantly, the stability of such separation timing over days. In (Fig. 3f, Supplementary Fig6c), we showed that for drifting gratings, which consist of both orientation and direction features, the mean feature inter-stimulus feature separation between the eight orientations occurred about 50 ms earlier than the feature separation between the two directions. This latency was stable across 15 days. Similarly stable was the timing of representation separation under static gratings, except that it occurred at a different time point (Fig. 3g, Supplementary Fig6d). These results suggest that time-resolved populational representations encode rich dynamic features of the stimuli and hold remarkable longitudinal stability.

### The temporal dynamics of individual neurons enhance tuning stability

We next investigated if and how much temporal coding, compared to rate coding, carries additional information about external stimuli and its impacts on tuning stability. Using drifting gratings as an example, we noticed that different motion directions of the grating brought out distinct features in a neuron’s temporal spiking patterns (Fig. 4a). For instance, the grating moving at 202.5° triggered sustained activation between 100-300 ms (red), unlike the response to the 22.5° stimulus, which had the same orientation but opposite drifting direction, and lacked a ramp-up in activity after 400 ms (blue). These subtle yet clear differences in response patterns remained stable over 15 days, underscoring that average event rates alone do not fully capture these nuances.

To effectively extract temporal codes, we utilized cross-validated, regularized linear discriminant analysis (LDA) to determine the temporal components that best separated the different stimulus conditions. This supervised decomposition pinpoints variations that directly relate to stimulus decoding while being less indicative of spontaneous activity^45^ or variabilities induced by mixed coding^44, 46, 47^. Fig. 4b displays the top three components of the representative unit fitted with all trials over 15 days. They intuitively represented multiple aspects of spiking time courses that are not easily inferable from rate coding: the duration of activation (CP1), temporal contrast (CP2), and the preferred timing of activation (CP3). We observed diverse temporal profiles across all neurons (Supplementary Fig5b, see Supplemental Discussion). Importantly, we note that firing rate is also a special type of temporal components that has a temporally flat profile^48^. Therefore, to reduce the information overlap between rate codes and temporal codes, we discouraged LDA from finding temporally flat weights by normalizing each trial with the area under the peristimulus time histogram before fitting. This allows us to more clearly characterize the unique contribution between the two types of neural codes.

Considering these temporal components significantly improved the classification of different stimuli encoded by individual neurons. Using the responses of a representative neuron to drifting gratings as an example (Fig. 4a-b), the distributions of daily, trial-averaged firing rates showed high overlap among the 16 different stimuli (Fig. 4c). In contrast, when we represented the 16 stimuli using the daily averaged first three components, inter-stimulus separation notably improved, making similar stimuli such as 0°, 180°, 22.5°, and 202.5° clearly distinctive (Fig. 4c). Furthermore, we examined the decoding accuracy of individual neurons in single trials. Due to overlapping responses across different stimuli and high trial-to-trial variability^7, 49^, the accuracy was only 0.0158 ± 0.001 when using firing rates compared to the chance level at 0.01 (100 stimulus classes of natural images stimuli). Yet, temporal components increased decoding accuracy to 0.0178 ± 0.002 (P < 7e-77, paired t-test). This increase in accuracy when utilizing temporal code over rate code was consistent across all four stimulus types. (Fig. 4d, Supplementary Fig5c).

Next, we examined the contribution of the temporal code to longitudinal tuning stability. Across all stimuli, unit tuning curves were more similar to themselves after 7 days (Fig. 4e, similarity quantified by correlation coefficient after Fisher z-transformation, unbounded) when derived jointly from temporal components 1-3 than those derived from firing rates (i.e., those discussed in Fig. 2). Considering the entire recording duration, temporal components-based tuning had a smaller rate of change in representational drift index^10^ (a smaller slope, P < 2.7e-41) than firing rates-based (Fig. 4f). The drift reduction effect was particularly prominent for neurons having poorer firing rate-based reliability (Fig. 4g, Fig. 2). Importantly, merely increasing the dimension of firing rate-based tuning did not lead to decreased drift across time (Supplementary Fig5d). This is further strengthened by the fact that top three components extracted from a simpler method (PCA) failed to reduce drift (Supplementary Fig5d). Finally, high dimensional PSTH (before LDA or PCA dimension reduction) failed to meaningfully reduce drift (Supplementary Fig5d). Figure 2b showed that the stability of single-neuron firing rate-based tuning curve was associated with tuning reliability, strength, and selectivity. Similarly, these associations held true when stability, reliability, strength, and selectivity were derived from tuning curves of temporal components (Supplementary Fig8). Particularly, tuning reliability also increased when considering the top three components compared to using firing rates (P < 1.5e-29, Supplementary Fig9).

Together, these results suggest that time-resolved electrophysiological recording could empower us to use “time as a coding dimension.”^18, 24^ Importantly, our stable chronic recording revealed that temporal components-based tuning supports substantially greater stimulus representation reliability and stability at the single neuron level beyond what could be achieved by conventional firing event counting over seconds-long integration windows^7, 11^.

### Temporal dynamics enhance long-term populational representation discriminability against competing stimuli

We next extended the study from single neuron tuning to the longitudinal stability of visual representations formed by neuronal populations within each animal. We constructed populational representations on “super sessions,” which involved pooling tuning curves from neuron populations across all longitudinal sessions for joint dimension reduction. This approach allowed us to quantify the chronic stability of neural representation from different days in the same feature space, providing an intuitive reference for stability and stimulus distinctions.

Since linear methods cannot effectively reduce the dimensionality in V1^50^, we computed stimulus representations in Uniform Manifold Approximation and Projection (UMAP)^51^ space using both the firing rate tuning and the temporal component-based tuning (Fig. 5) and observed the following features: First, in both cases, the populational representations exhibited clear spatial structures that correlated with features of the stimuli (Fig. 5a-b, e-f and Supplementary Fig10a-d). For example, drifting gratings formed a ring manifold, with representations of nearby orientations as nearest neighbors, and representations of the same orientation but opposite directions (e.g., 0° and 180°) in similar angular positions on the ring (Fig. 5a). Static gratings arranged in five different arcs reflecting five spatial frequencies, and within each arc, nearby orientations formed clusters that were close to each other (Supplementary Fig10a-b). Representations of receptive field Gabors formed a continuous manifold that mirrored the geometry of the 9 x 9 grids (Supplementary Fig10c-d). These results confirmed that the recorded neural population was sufficiently large and functionally diverse to not only distinctly encode a wide range of stimuli but also form smooth representation manifolds^50^ in the neural space where similar visual stimuli were encoded to adjacent locations. Furthermore, representations of the same stimuli across all days were closely spaced and formed distinct clusters, indicating the longitudinal stability of the representation. For example, in natural image stimuli, the representations of 100 images across 15 days formed 100 distinct clusters, each of which consisted of 30 points representing the even/odd trial averages of 15 daily representations of the same image (Fig. 5e-f, Supplementary Fig11a-b)

**Fig. 5:**
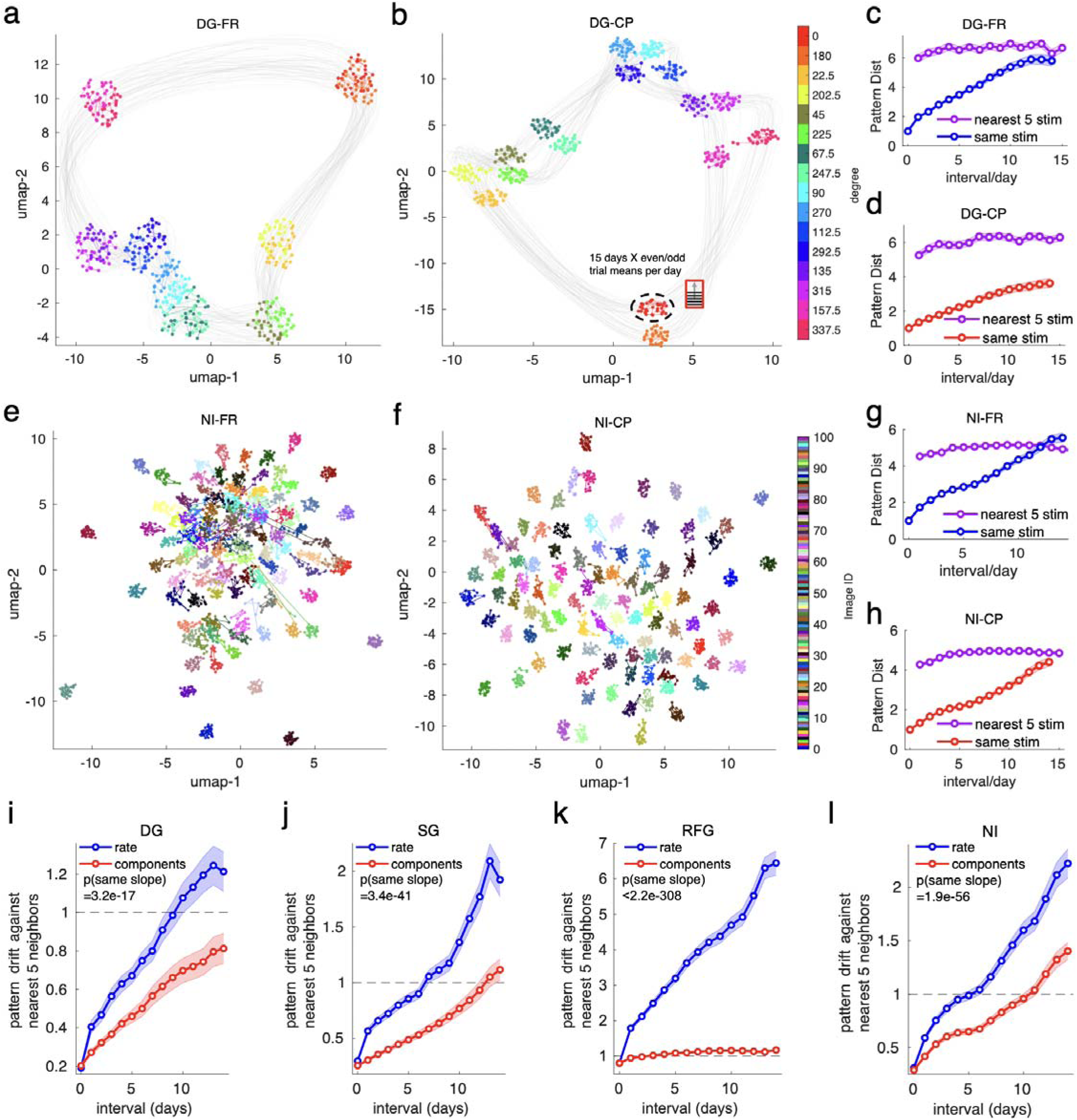
Temporal codes stably increase long-term populational representation discriminability against competing stimuli. **a-b**. For drifting gratings, different stimulus conditions (directions) were more distinctly and stably represented in low-dimensional UMAP space when represented using populational temporal components (b) compared to populational firing rate (a). The same stimulus formed distinct clusters of 30 points (15 days x even/odd trial averages, linked by lines). Clear spatial structures were observed with continuous geometry (i.e., nearby directions for drifting grating (a)). The original high-dimensional representation was constructed with all tuned neurons that appeared for 15 days in 1 example mouse. **c-d**. UMAP representations of the same drifting grating directions were more similar to themselves across sessions than to the nearest 5 stimuli from the same session, especially when represented by the top 3 temporal components (d), compared to that of firing rates (c). **e-h**. Same as in a-d but for natural images stimuli. e-f shows representations of center 97 images, for all 100 images, see Supplementary Fig11a-b. **i-l**. Statistics of Fig. 5c-d. Fig. 5g-h, and the other two stimulus types. Across four stimulus types (DG: drifting gratings; SG: static gratings; RFG: receptive field Gabors; NI: natural images), the separation between the same stimulus across days normalized by the distance to nearest five different stimuli within days (black dashed line) drifted at a lower rate for components-based representation than that of firing rate-based representation. All lines show the mean ± s.e.m. Linear mixed effect: time, representation definition method as fixed effect, animal-stimulus pairs as random effect (n=80,150, 405, 500 pairs for DG,SG,RFG,NI respectively). See Supplementary Table 1 for additional reporting on sample size and statistics.

Critically, using temporal components to construct the populational representation significantly improved the isolation between clusters of the same stimuli relative to those of competing neighbors (Fig. 5a-b, e-f, Supplementary Fig10a-b,c-d). To quantify this difference, we computed and compared the Euclidean distance of representations of the same stimuli across 15 days and that of the closest other stimuli within days in the low-dimensional UMAP feature space, all normalized by the average within-day distance between even and odd trials of the same stimulus. The distances within the same stimulus increased with time in both firing rate and temporal components-based representations. However, firing rate representations (Fig. 5c, g) approached or surpassed the distance to the five nearest stimuli at the end of the 15 days at a faster rate than that of temporal components-based representations (Fig. 5d, h). Statistically, compared with firing rate representations, temporal components better maintained the separation of a given stimulus across days against its nearest five stimuli within-days for the entire experimental period (smaller slope, P < 3.2e-17, Fig. 5i-l).

As another test of stability in visual representations, we investigated how much temporal information could increase single-trial causal decoder performance in future days without any recalibration. We fitted decoding models (regularized LDA) separately for each animal using simultaneously recorded neurons that appeared for most days (at least 12 days and must appear for all testing days). To causally incorporate temporal information, a new set of temporal components was fitted for each neuron only using trials from the first 7 days. Importantly, consistent with our findings for individual neuron decoding (Fig. 4d, Supplementary Fig5c), we found that at the population level, adding temporal components in addition to firing rate in the fixed decoder consistently reduced decoding error for all four stimuli across 8 future testing days when trained on the first 7 days (P < 0.0089, Supplementary Fig10e Supplementary Fig11, reduction by 5.0°, 2.4°, or 2.0 and 10.0 folds over chance for drifting gratings, receptive field Gabors, static gratings, and natural images respectively, with chance levels being 90°, 35.4°, 1/30, and 1/100 respectively). To depict the drift in decoding performance, we defined decoding drift index like that of representational drift index. We observed that adding temporal components reduced decoding drift for natural images and static images stimuli (P < 0.00038, Supplementary Fig10e), where no differences in decoding drift were evident in receptive field Gabors or drifting gratings stimuli, which might be because representation of natural stimuli drift more than that of artificial stimuli^10^. Together, these results revealed that populational representations of visual stimuli were better discriminated longitudinally when temporal coding was captured.

### Stability of temporal codes associated with network connectivity

We have uncovered the importance of temporal code in stabilizing visual representations against rate code. Yet, what makes a neuron respond stably to visual stimuli? The current literature suggests that many aspects of a unit’s tuning property stem from its connectivity in the neural network^42, 43, 52–55^. To test the association between temporal code stability and network functional connectivity, we measured and longitudinally tracked the functional connectivity among the population of neurons and examined their relationship to the tuning stability of temporal code, using that of rate code as a control.

We first quantified the stability of putative monosynaptic connections with pairwise cross-correlograms (CCGs) across days with a temporal resolution of 1 ms. This analysis not only relies on stable tracking of both neurons in any connection pair but also requires a large number of neurons to be recorded, as significantly functionally connected CCG pairs are rare, accounting for only about 1% of the total possible number of pairwise connections^28, 30, 56^. We showed an example of recurring jittered corrected CCGs^28, 57^ across sessions, manifesting consistent temporal lag, functional shape, and similar amplitude (Fig. 6a). Within an animal, the connections showed stimulus-specific patterns that were visually similar across adjacent days (Fig. 6b, Supplementary Fig12a, Supplementary Fig13). Quantitatively, we observed indications of overall network stability: the number of connections (P > 0.11) and average peak amplitude per animal (P > 0.39) remained stable for each stimulus, except that the peak amplitude during drifting gratings had a marginal rate of change at −8.5e-5/day (P = 0.005) (Fig. 6c-d, Supplementary Fig12b-c). Despite the overall network stability, we also observed gradual and subtle drift in the connectivity matrix similarity across days for all 4 stimuli (P < 3.7e-16, Supplementary Fig13)

**Fig. 6:**
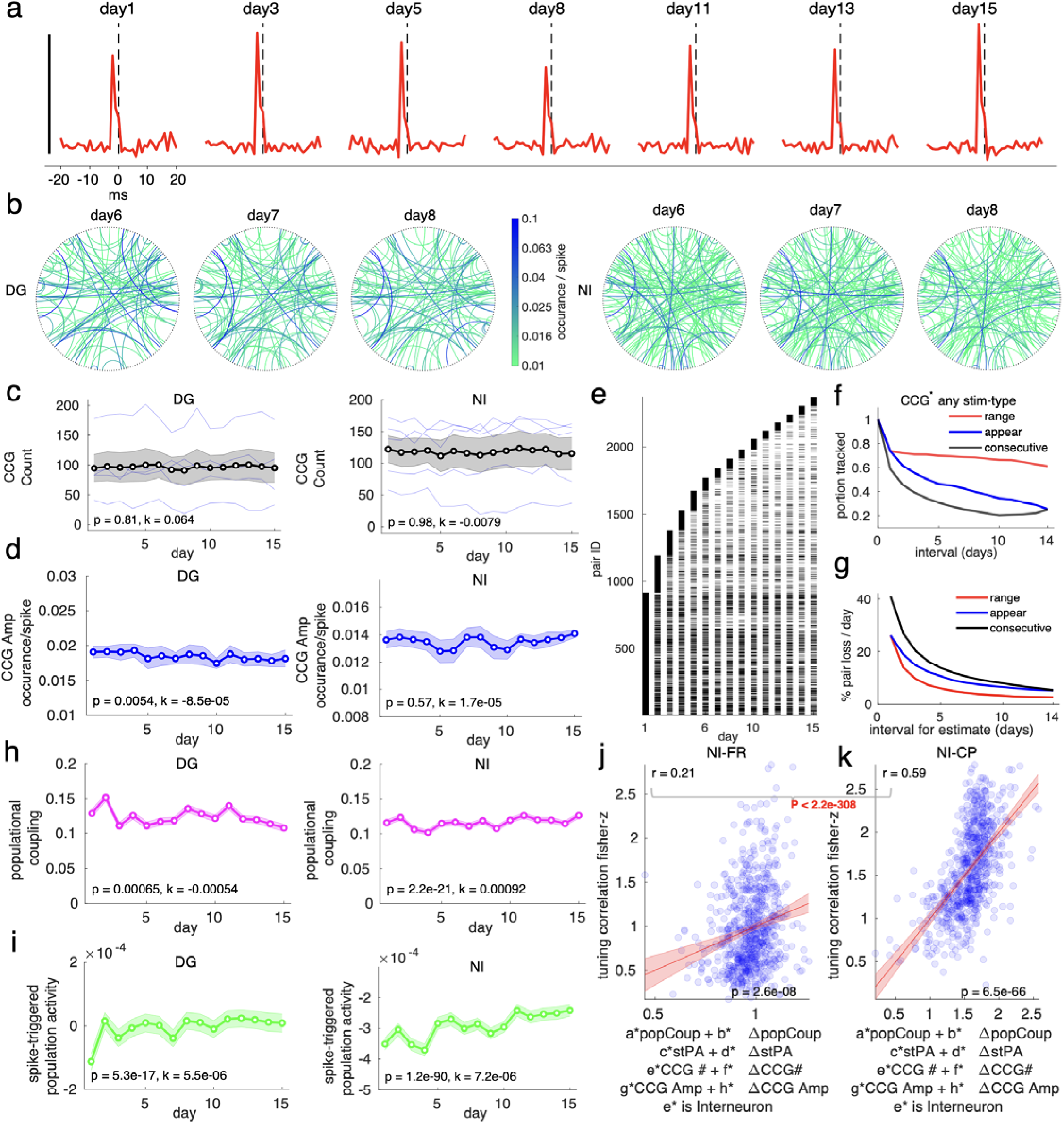
Stability of temporal codes associated with network connectivity. **a**. An example pair of CCGs having a similar shape to itself across 15 days; sharp peaks were found at lags within 4 ms; scale bar: 0.05 occurrence/spike. **b**. All significant CCGs (curved lines, color-coded by peak amplitude of the CCG) between neurons (black dots aligned on the big outer circles) in one mouse during the 2 stimuli (DG: drifting gratings. NI: natural images) showed similar connection patterns across days (columns), with neurons randomly placed on the circle but the order fixed across days and stimuli. Only neurons that had at least 1 significant connection during either DG or NI stimuli across 15 days were included (n=189). **c-d**. The CCG strength (c) and count (d) remained stable for the two stimuli. DG: drifting gratings. NI: natural images, showing the number of significant CCGs per animal (c) and mean CCG peak amplitude (d) across 15 days. Linear mixed effect, time as fixed effect, individual animals (n=5) as random effect (c), individual significant CCG pairs (n=786,905 for DG and NI, respectively) as random effect (d). Lines are in population mean ± s.e.m. **e**. Huge amounts of pairs were detected across 15 days. Pairs could be tracked for multiple sessions. Longevity tally for CCGs identified in response to any of the 4 stimuli. Pairs were sorted according to the initial session of appearance. **f-g**. Quantification of the fraction of trackable CCG pairs in (e), showing tracking probability based on 3 criteria, number of sessions appeared, consecutive sessions appeared, and the range between the first and last seen session (f), and the resulting estimated percentage pair loss per day over 14-day intervals (g). See Supplementary Fig13 for count data in (f) before taking the fraction. See Supplemental Methods for quantification explanations. **h-i**. Single neuron-to-population synchrony changed minimally across time for the two stimuli. DG: drifting gratings. NI: natural images. The population synchrony score was measured either by population coupling (h) or spike-triggered population firing at 0 ms (i) or for all single neurons that appeared for more than 2 days from all mice. showing mean ± s.e.m. Linear mixed effect model (LME), time as fixed effect, individual neuron (n=1037 single units appearing more than 2 days) as random effect. **j-k**. Functional connectivity metrics better explained temporal component-based natural images tuning stability after 7 days than that of firing rate-based. Correlation coefficient (y-axis) was Fisher Z-transformed. Stability during drifting grating stimuli was defined using either firing rate (j) or the first 3 components (k) and was jointly explained by 1. population coupling (popCoup), 2. spike-triggered population activity at 0 ms lag (stPA), 3. Total incoming CCG counts (CCG #), 4. total CCG strength (CCG Amp), 5. the rate of change “Δ” of variable 1-4 per day, 6. cell type. Red line: linear fitted response with 95% CI, text: correlation coefficient (square root of R^2^ of linear regression), significance level difference in correlation (Z-test). 684 stimulus-tuned single units (dots) that reappeared after 7 days. See Supplementary Table 1 for additional reporting on sample size and statistics.

Across all stimuli, we identified over 2000 connections across 15 days cumulatively (Fig. 6e). Yet, the probability of tracking the same pairs gradually declined over the duration of tracking (Fig. 6f, Supplementary Fig13), irrespective of the quantification methods of tracked duration. Notably, the day-to-day turnover of these significantly functionally connected pairs reduced with time interval (Fig. 6g), in line with a report that longer measurement durations reduce the estimated drift rate^58^. At the maximum measurement interval (Day 1 to Day 15), the average pair loss rate was 2.8% per day, which is a relevant comparison with the findings of imaging studies showing that the synaptic turnover rate is as high as 1% per day in the visual cortex^42, 59^.

It is plausible that pairwise CCGs do not fully represent the neuronal connections in the network. Therefore, to further characterize the stability of the associated network supporting individual neurons, we evaluated the neuron-to-population synchrony in terms of spike-triggered populational activity (stPA) at the 0 ms lag^54, 60^, and population coupling^42, 54^ (PC), the correlation between stimulus response from one neuron to the summed response of the rest of the population. In contrast to those in the CCGs, we found that PC and stPA themselves changed significantly over time (P < 0.0007) for all stimuli (Fig. 6h-i, Supplementary Fig12d-e). However, the rate of change was minimal, at no more than 0.001 per day for PC and 8e-06 (occurrence/spike) per day for stPA. Individually, more than 95% of the single units tracked for more than 2 days remained stable in the PC and stPA measures (P > 0.05, Bonferroni corrected, Supplementary Fig12f-g).

We next built linear models associating a unit’s tuning stability after 7 days with these network functional connectivity measures, including the magnitude and daily rate change in population coupling, spike-triggered population activity at the 0 ms lag, total incoming CCG counts, total incoming CCG strengths, and cell type as another proxy variable for the connectivity motif. We found that the firing rate stability was weakly to moderately associated with these factors combined (Fig. 6j). In contrast, this linear model predicted the tuning stability of temporal components to a greater extent (P < 9.1e-07) than that of firing rate for all stimuli but drifting gratings (Fig. 6j-k, Supplementary Fig14a-c, Supplementary Fig15a-d, Supplementary Fig16). These results thus showed that the stability of temporal codes was better associated with network functional connectivity than that of rate code.

## Discussion

A consistent neural activity landscape is instrumental in maintaining regular brain functions, such as enabling stable sensory representation of the external world, which forms a reliable basis for informing appropriate downstream actions. Understanding the baseline stability related to sensory experience at the single neuron, neural population and inter-neuron functional connectivity levels not only unravels the mechanism of such stability but also provides a solid reference point from which we can gain further insights into the brain’s ability to modify its internal structure to facilitate memory, learning, and adaptation in response to evolving environments.

Comprehensive Identification of the neural substrates that support consistent visual perception is challenging experimentally and requires 1) faithful, large-scale longitudinal monitoring of spiking patterns from the same sensory neuron populations^42, 61^ 2) a measurement method that induces minimal acute and chronic alterations to the neural circuit being measured; and 3) diverse^50^ and repeated^40^ visual stimulation.

Contemporary two-photon calcium imaging studies have revealed various amounts of representational drift under diverse types of stimuli, with the representation of naturalistic patterns^10, 11^ having more noticeable drift compared to those of artificial grating stimuli^7–10, 43^. Despite the merits in large-scale monitoring, cell-type specificity, and neuron reidentification confidence, calcium transients only provide a proxy for spiking with limited temporal resolution^13–17^. Furthermore, the use of high-power lasers carries inherent technical limitations by introducing the risk of phototoxicity, photobleaching, and photo-interference^62^, all of which may interfere with the very circuit whose stability we wish to characterize. In addition, cranial imaging window surgery could disrupt normal physiology like hemodynamics^63, 64^ and glymphatic flow^65^, change synaptic connections^66^, and activates immune cells like astrocytes and microglia^63, 65, 67^, which may take over one month to fully recover^63, 65^, while waiting for the recovery increases the chance of window clarity degradation^68^ from bone and dura regrowth^69^.

Two-photon imaging studies also have trade-offs between within-day experimental duration and across-day inter-session repetition frequency (to mitigate photobleaching), with both factors being crucial for characterizing visual representation stability. Within-session experimental time is vital for ensuring sufficient stimulus diversity and repetition of the same stimuli. Repetition is important because spike generation is a stochastic process^40^ and requires a decent number of repeats to form reliable trial-averaged responses and calculate trial-to-trial reliability. While one could focus on a few stimuli conditions while boosting trial count^68^, this approach misses the opportunity to capture stability in the high-dimensional stimulus response space^50^ specific to the visual cortex, where responses from one stimulus type cannot be easily predicted by responses from other stimulus types^70^. Across-sessions repetition frequency is also crucial for revealing the dynamics of chronic responses, particularly because stimulus familiarity^5, 32, 33^ has been shown to affect representation stability.

On the other hand, long-term electrophysiology offers intrinsic advantages such as high temporal resolution^15, 17^, absence of photobleaching, and increased freedom in terms of experiment duration and frequency^71^. However, electrical recording has conventionally suffered from scalability and longevity issues in tracking the same neuronal populations, attributed to the chronic neuroinflammatory response associated with traditional rigid electrodes^72^. The mechanical mismatch at the tissue-electrode interface could induce substantial micromotion and degradation, making longitudinal tracking of large neuronal populations highly unreliable and hence rarely available. In the literature, with heroic efforts, only a handful to a few dozen neurons are longitudinally followed^1, 36, 73^. Recently, high-density silicon probes have enabled chronic^27^ large-scale^28^ recording in mouse visual cortices, yet their ability to study longitudinal representation stability has not been clearly established.

We believe our platform could systematically tackle these challenges. We performed large-scale implantation of ultraflexible electrodes that integrate seamlessly with the surrounding tissue. Accordingly, we tracked more than 1000 neurons for an average of over 10 daily sessions. With largely unconstrained experimental duration and repetition frequency, we tested four visual stimulus types with 137 artificial and 100 natural patterns, for a total of 11930 trials per day for 15 consecutive days, providing a comprehensive evaluation of both within-session representation reliability and across-sessions visual representation stability across diverse stimulus types. This could inspire many exciting questions such as predicting the stability of one stimulus type from other stimulus types (Supplementary Fig17). Additionally, the long-lasting nature of NETs allowed us to perform these functional stability measurements at least 60 days after surgery to ensure a more complete recovery from implantation trauma^31^, minimizing the circuit instabilities induced by the measurement technology itself.

Armed with this tailored technical platform, we compared the visual representation stability between rate coding and temporal coding in the mouse visual cortex. We found that neurons had varying day-to-day rate code tuning stability to diverse types of stimuli, with across-session stability strongly associated with tuning reliability. In line with a report that tuning reliability may underlie the functional specialization of different brain regions^74^ and functional evolution of developmental stages^75^, our results further showed that response reliability could be an indicator of chronic rate code representation stability (Fig2).

Notably, we emphasized the time-resolved visual representations and temporal code. In addition to showing the existence of stable stimulus-evoked spiking dynamics across time at the individual neuron level^32, 71, 73, 76^, here we extended these studies into the evolution of population-level visual representations (Fig3) along stimulus-coding dimensions at 10 ms resolution across 15 days and found overall stable, distinct dynamical representations. Our results suggested the existence of consistent timing of separation between stimulus categories (e.g., directions versus orientations) across 15 days. The consistency among different stimulus types (drifting gratings versus static gratings) indicated that the observed stability in the timing of separation was not limited to a specific stimulus but was likely a generalized phenomenon. Further investigation could potentially unveil more important insights into the neural processing pathways that underlie the observed variations in activation timings of different stimulus features.

We next elucidated the benefit of incorporating temporal information longitudinally. Prior electrophysiological studies have provided evidence for the existence of temporal codes: individual neurons generate distinct spiking time courses in response to different stimuli^24, 36, 71, 73, 76^, with a possible underlying mechanism being the activation of multiple recurrent and feedback circuits^22, 23, 52, 77^. Here, we further showed that temporal components exhibited reduced drift in single neuron stimulus tuning compared to firing rates-based tuning. The reduction was particularly prominent for neurons with low firing rate-based reliability. In addition, when using stimulus-coding temporal components to form neural population representations, visual patterns were more distinct from competing stimuli and more stable across days compared to rate coding. Similarly, the stimulus identity of familiar scenes could be more stably identified from causal decoders trained with past recording days by incorporating temporal code. Thus, temporal coding carries more stable visual representations than rate coding.

The mechanisms underlying the enhanced stability of temporal codes over rate codes are likely multifaceted and complicated. We made the following observations while attempting to understand it: 1. A breakdown characterization for each individual component (Supplementary Fig18a) showed that the effect was not due to individual components themselves drifting less than firing rates. 2. the effect was not due to individual temporal components having a higher selectivity^38^ than that of averaged firing rate (Supplementary Fig18b) 3. We analyzed the representative neuron in Figure 4 and showed that the enhanced stability of component-based tuning compared to that of the average firing rate might have been achieved by “temporal gating,” which suppressed variations in non-coding dimensions. For instance, in the 15-day tuning curves to drifting gratings (Supplementary Fig18c), the neuron had a higher firing rate during 180° stimuli on day 1 than on other days. Yet, such potential non-coding excessive activation was only partly projected to the 1^st^ component while minimally affecting the 2^nd^ and 3^rd^ components, thus still preserving sufficient stability. Similarly, it is observed^12^ that drift is higher in “non-coding” dimensions, despite visual responses being studied on a different timescale and using a different neural code extraction mechanism. In the same vein, temporal codes were less explained by non-stimulus variables (Supplementary Fig18d). These finding might be consistent with a recent report^47^ that fast spiking dynamics could de-mix responses to multiple sensory inputs.

Finally, we found that network functional connectivity metrics explained tuning stability, especially when tuning was defined with stimulus-coding temporal components. This aligns with the view that visual neurons have many synaptic partners that are tuned to different stimulus features^53, 78–80^. They are also influenced by diverse feedback connections from downstream areas^22, 23, 77, 81^. Since these connections are likely activated at different timing during stimulus processing, they contribute to finer dynamic structures of a neuron’s tuning and induce the observed temporal coding^19, 21^. Therefore, network functional connectivity metrics better explained the stability of temporal code than rate code at the single unit level. Recently, it is proposed^82^ that drift can result from both experience dependent Hebbian plasticity and experience-independent synaptic volatility. Since the novelty of the visual experience was not explicitly manipulated in this study, our findings suggest that temporal codes may better capture synaptic volatility compared to rate codes. The role of temporal code in Hebbian plasticity awaits further studies.

In conclusion, we provided an elaborated survey of the day-to-day stability of visual representations with temporally resolved, stable, large-scale recordings using ultraflexible electrode arrays. We tracked the temporal spiking patterns of the same neuronal populations in the mouse visual cortex in response to diverse types of stimuli for 15 consecutive days. We found that temporal coding carries more stable visual representations than rate coding at the single -neuron and population levels compared to rate coding. The stability of the temporal code was better associated with network functional connectivity than that of rate code. We therefore posit that temporal coding might play a crucial role for the brain to maintain consistent visual representations across time.

### Limitations and Future Studies

The characterization of representation stability depends on understanding of the represented information itself. Contemporary studies have demonstrated the widespread phenomenon of mixed representation in the cortex^46^ including in visual areas^41^. These findings lead to the natural question of whether the day-to-day fluctuations in non-visual aspects of animal behavior (e.g. attention) contribute to the observed drift in visual representations^58, 61, 83–85^. We partially mitigated this effect by explicitly projecting spiking time courses on to stimulus coding dimensions (Fig. 3-5) via LDA. Future studies could thoroughly compare other ways of extracting temporal components/incorporating temporal information^18, 48, 86, 87^ to better represent sensory-specific inputs. Additionally, future studies could further build deterministic models of single neuron spiking^37, 88, 89^ to remove non-visual stimulus information and hence test if the residual stimulus encoding activity is even more stable across days. Furthermore, the visual cortex is far from homogenous, future studies could extend the stability characterization to specific cell types^33, 90, 91^ and the other higher visual areas. We also acknowledge limitations common to all electrical recordings, e.g., only neurons/cell types that fire sufficiently frequently are reliably sortable^42^. In addition, Calcium imaging still offers substantially higher throughput in terms of chronically trackable neurons per mouse. Finally, in this work, we studied the magnitude of representation stability under passive exposure to familiar stimuli. This serves as a critical reference point for future studies on the direction of visual representational drift, especially when steered by novel experiences^32, 82, 90, 92^ or active reward learning^93–95^.

## Methods

### Fabrication

The fabrication process has been previously reported^96^. 3nm Ti and 60 nm Ni were patterned on glass substrates as the initial layer, which was eventually sacrificed to release the flexible end of the probe. Next, we spin-coated and heat-treated ∼500 nm polyimide (PI) (PI2574, HD Microsystems, NJ, USA) to create the bottom layer of insulation. On top of that, a stack of 3 nm Cr, 100 nm Au, and 3 nm Cr was electron-beam evaporated (Sharon Vacuum Co., Brockton, MA) to create a layer of interconnects between electrode sites and backend bonding sites under photolithography. Bonding pads were further deposited with 3 nm Cr, 160 nm Ni, and 80 nm Au for improving soldering reliability. Another ∼500 nm PI layer served as the top layer of insulation and was identically fabricated as the bottom layer. To create the outline of the probe, vias to the electrodes and solder pads, RIE etching (Oxford Instrument) was performed with 9:1 O2/CF4 gas. Finally, in 20 of the probes, electrode sites were sputter coated with 10 nm Ti, 100 nm Pt, 10 nm Ti, and 300 nm IrOx (AJA ATC Orion Sputter System) and defined under photolithography. The other 8 probes were sputter-coated with Cr/Au at thicknesses of 5/120 nm, respectively. The total thickness of a probe was 900-1100 nm.

Three probe designs with different arrangements of the 32 channels were incorporated in this study. Design-1 electrodes were made of IrOx and were arranged into a 16×2 (depth x width) matrix, with both depth and width electrode spacing at 20 μm (center-to-center). The electrodes were rectangular in shape and sized 8 μm x 6 μm. The shank width tapered from 110 μm to 85 μm. Design-2 electrodes, made of Au, were arranged in a 16×2 (depth x width) matrix with depth and width electrode spacing at 31.5 μm and 30 μm center-to-center, respectively. The electrodes were rectangular in shape and sized 19.5 μm x 12 μm. The shank width tapered from 120 μm to 60 μm. Design-3 electrodes were made of IrOx, whose 32 channels were linearly spaced at a spacing of 25 μm center-to-center. They had rectangular contacts sized 12 μm x 6 μm. The shank width tapered from 105 μm to 80 μm.

After fabrication, a customized printed circuit board (PCB) was soldered onto the matching bonding pad sections of the device using stencil-aligned high-temperature solder balls. The implantable portion of the probe was released by immersing it in nickel etchant (Type I, Transene Inc., MA, USA) for 5 minutes at room temperature, and the corresponding section of glass substrate was subsequently cleaved. All probes were electroplated with PEDOT/PSS in phosphate-buffered saline, targeting a resistance of 30 - 100 kΩ. The released probes were grouped into four per layer and placed on four parallel 50 μm diameter tungsten wires (W5574 Tungsten Wire, Advent Research Materials, Oxford, England) and affixed with bio-dissolvable adhesive Polyethylene glycol (PEG).

### Animals

A total of 5 male mice were used in the experiments, including n=2 adult C57BL/6J-Tg (Thy1-GCaMP6s) GP4.3Dkim/J mice bred on-site and n=3 C57BL/6J mice acquired from Jackson Laboratories (Bar Harbor, ME). No mice were excluded. To rule out the potential effect of hormone fluctuations on representational drift and given we did not record at the time scale to measure these changes, all male mice was used. Further work could explicitly study the differences in representations of female mice at different hormone phases. Animals were housed in normal dark/light cycle (7a lights on, 7p lights off), with ambient temperature 68∼72°F and humidity 30∼70%. All surgical and experimental procedures in this study followed the National Institutes of Health Guidelines for the Care and Use of Laboratory Animals and were approved by the Rice University Institutional Animal Care and Use Committee.

### Surgical procedure

The surgical method has been previously published^31^. Briefly, animals were anesthetized with isoflurane (3% for induction and 1%-2% for maintenance) and administered extended-release Buprenorphine (Ethiqa TM) for analgesia and Dexamethasone (2 mg/kg, SC) for anti-inflammatory effects. The skull was exposed by removing surface hair and skin, followed by the removal of the fascia, and scoring of the skull in a crosshatch pattern to enhance cement grip. A circular craniotomy of 3 mm in diameter over the visual cortex was drilled in the skull for the NETs implantation, and a separate hole was drilled in the ipsilateral hemisphere +1.5 mm ML, +1 mm AP for implanting a Type 316 stainless steel grounding wire. Groups of probes were implanted simultaneously through the dura to the right hemisphere V1 and LM region targeting +(2-4) mm ML, −4 mm AP, within 1 mm depth, with inter-probe spacing about 700 μm, though variations existed to avoid surface vasculature. We inferred whether a shank is V1 or LM using the estimated target location: Given released probes were grouped into four per layer and placed on four parallel tungsten wires spaced about 700 μm before implantation, we typically expect the medial three shanks in V1 and the most lateral shank in LM. V1 and LM were targeted given they were reported to be among the areas having the least drift in within-session representation^11^. When a mouse received more than 4 probes, the second group targeted +(2-4) mm ML, −3.3 mm AP. Following implantation, the tungsten wires were removed after the PEG dissolved, and a sterile glass coverslip window (Harvard Apparatus, 1217N66) was placed over the craniotomy with its border sealed with Kwiksil (World Precision Instruments). Additional layers of cyanoacrylate and Metabond dental cement (Parkell, NY) further secured the window and allowed for the attachment of a head bar for head fixation. A total of 25 probes were implanted successfully, three animals received 4x, 4x, 3x Design-1 probes respectively. One animal received 4x Design-1 probes and 3x Design-3 probes. One animal received 4x Design-1 probes and 3x Design-2 probes. Following multiple reports that stable units were either recorded or emerged months after surgery^5, 35, 36, 97^ and that tissue surrounding the ultraflexible electrodes implantation continued to recover for at least a month^31^, animals were provided at least 50 days of recovery post-surgery and an additional 12 days (median) of familiarization to head restraint and visual stimulation before the beginning of consecutive daily visual stimulation.

### Large-scale electrophysiological recording and spike sorting

The electrophysiological data was then recorded using the Intan RHD USB interface board (Part #C3100) and the 128-Channel Recording Head stage (Part #C3316). Broken channels, typically with an impedance greater than 3 MΩ at 1 kHz measured by the same head stage, were excluded from further analysis.

The recordings were performed on awake head-fixed mice on a custom-made Styrofoam running wheel. The treadmill was fixed on an optical breadboard with optical posts and the whole setup was floated on an anti-vibration table (Thorlabs, PTT600600). Passive visual stimulation was administered for 15 consecutive days. Each recording session lasted about 2.5 hours.

Spike sorting was performed in Mountainsort4^98^. Common median referencing^99^ was applied to the high-pass filtered data from each shank to reduce common mode noise such as motion artifacts. The spike detection threshold was set to positive and negative 3.5 times the standard deviation (default: 3), with a detection interval of 12.5 samples (default 10 samples), and the adjacency radius was set to 100 μm. Each session was sorted individually, and only common channels available for all 15 days were used to track putative same units (Supplemental Methods).

Noise clusters were rejected when the following criteria were not met simultaneously: Mean waveform peak-to-peak amplitude (P2P) > 25 μV; Firing rate (FR) > 0.01 Hz; Full width at half-maximum (FWHM) of the mean waveform (top channel) at 0.064 ms < FWHM < 0.9 ms; Trough to peak time (TP) of the mean waveform at 0.08 ms < TP < 1.02 ms; Mean top-channel normalized amplitude of 2^nd^ through 20^th^ largest channels should be smaller than 0.5; The peak of mean spike waveform should be within ± 0.12 ms of spike-sorter reported peak time.

Putative single units were identified using the criterion that the fraction of spikes with inter-spike intervals under 2 ms were less than 1.5% of the total firing events. This criterion is comparable to those used for two long-term tracking publications (1.5 ms, 2%)^5^ and (2 ms, 1%)^99^.

### Location estimation

The mean waveform of units was used for estimating their locations. The classical estimate of unit locations from oversampled electrodes is the weighted centroid method^100, 101^. Given recent advances in spike source modeling, we weighted a point source model estimate^102^ and a weighted centroid estimate at a 70%:30% ratio for better visualizing the displacement of units across 15 days in the Supplementary Fig1a,f. This weighted scheme was used for calculation of firing rate drift over depth and amplitude drift over depth as well in Supplementary Fig1b. For each probe on each recording day, a depth histogram of unit firing rates or unit amplitudes over the entire recording was calculated at 0.1 μm bin. The histogram was smoothed with a moving mean filter of −5 to 5 μm. In other statistical analyses of chronic location changes over time, only the weighted centroid was used to be consistent with past literature^5^.

### Deep learning validation of labeling consistency

Tracking was also performed with dense neural networks to validate the consistency of the human labeling. For each sample neuron pair, the dense neural network took the 42 similarity-metrics together with the Autocorrelogram (ACG) similarity as input and outputted a probability that the two units should be combined (Supplementary Methods). ACG was calculated by Cell-Explorer^39^ at 1 ms resolution for 500 ms. Representative time bins were selected to reduce its dimension, and Pearson’s correlation coefficient between selected ACG bins was used to quantify the similarity of ACG for pairs of units. The time bins were [1, 2…20;21, 23…49;51,53,56,60,65,71,81,94,112,136,171,218,282,372,495] ms. The network consisted of 1 input layer, 2 hidden layers, and 1 single output node^103^. The activation functions for the 4 layers were rectified linear unit (relu), hyperbolic tangent (tanh), tanh, softplus. Adam optimizer with a default learning rate was selected to minimize the categorical cross-entropy of true versus decoded labels. Twenty models of different numbers of hidden units per layer (50∼500), portion of drop out (0∼0.5), number of training epochs (2∼15) were searched during the model fitting, and the best model was selected using Bayesian optimization with Gaussian process prior. The data were divided into 25 folds, each containing 24 probes of data as the training set while the remaining 1 probe as the testing set. Only unit pairs with d(WC2) <1.5 and Top4Xcorr<0.3 (See Supplementary Methods) were incorporated for training and testing, with those violating the thresholds automatically inferred as different units. The model fitting and hyperparameter search were repeated each time when we tracked units from another fold. The decoding quality was defined as the percentage of correctly recovered user decisions in the masked adjacency matrix for the test set at each probe. The same process was repeated using LDA as the machine learning model for comparison.

### Visual stimulation

Visual stimulation was conducted on awake, head-fixed mice. The visual stimulation was shown from a wide screen of 71 cm x 42 cm in size, which was placed 21 cm away from the left eye at a 45° angle to the animal’s rostral caudal axis and was refreshed at 60 Hz (frames per seconds). The screen was calibrated to have a luminance range of 0.01∼10 cd/m^2^. Visual stimulation was administered using the Matlab Psychtoolbox with full contrast. The stimulus patterns were not spherically warped, instead, a centered 60° aperture^104^ was applied to reduce changes in the viewing angle toward the edge of the screen from the animal’s perspective, while the rest of the screen remained in full-field gray. Four sets of stimuli were included in the study: drifting grating (DG), static grating (SG), natural images (NI), and receptive field mapping Gabor^28^ (RFG).

Each block of DG patterns consisted of 16 directions evenly spaced at 22.5° at a spatial frequency of 0.05 cycles per degree (cpd) and a temporal frequency of 2Hz. The initial spatial phase at the starting time of each stimulus was not modulated and kept fixed across sessions. Each stimulus trial lasted for 500 ms followed by a 500 ms gray screen. 50 blocks were presented resulting in a total of 800 trials per day.

Each block of SGs consisted of 60 patterns which factorize into 6 orientations (0°, 30°, 60°, 90°, 120°, 150°), 5 spatial frequencies (0.02cpd, 0.04cpd, 0.08cpd, 0.16cpd, 0.32cpd), and two spatially complementary phases (e.g., black-white-black vs white-black-white). Each stimulus trial lasted at least 250 ms followed by a 50 ms gray screen. 45 blocks were presented, resulting in a total of 2700 trials per day. Patterns of the same spatial frequency and orientation were grouped for further analysis, resulting in 30 patterns, each with 90 trials.

Each block of NIs consisted of 100 natural images from the ImageNet dataset^37, 105^. Each stimulus trial lasted for 250 ms (17 frames) with no gray screen in between. 60 blocks were presented, resulting in a total of 6000 trials per day.

Each block of RFGs^28^ stimulus was formed by spatially sparse drifting gratings of approximately 7° in size, 45° oriented with a spatial frequency of 0.05 cpd and a temporal frequency of 2 Hz. The Gabor patches showed up in 1 of 81 locations (9 rows and 9 columns). The exact azimuth, elevation, and size of each pattern was summarized in Supplementary Fig2. Each stimulus trial lasted for 245 ms with 50 ms gray screen in between. 30 blocks were presented, resulting in a total of 2430 trials per day.

For all stimulus types, the presentation order of the stimulus conditions was randomly initialized at the start of each block. Each recording session lasted about 2.5 hours. A light-to-frequency converter TAOS (part number: TSL235R) was attached to a corner of the screen so that visual stimulation onsets could be synced with electrical recording through Intan’s digital input. It later was observed through the photo-diode timing signal and a high-speed camera shooting at the screen that visual patterns did not disappear precisely at the programmed stimulus stop time, which might be due to the screen not refreshing precisely at 60Hz. We eventually used a stimulus on time of 520 ms (∼31 frames at 60Hz), 260 ms (∼16 frames), 245 ms (∼15 frames), 280 (∼17 frames) ms for DG, SG, RFG, NI stimuli respectively and consistently across all trials for accumulating firing rate or searching for temporal components.

### Visual response processing (rate code) and tuning significance

The single trial response of each recorded unit to a certain stimulus (e.g., a direction of the drifting gratings) was calculated by counting the total number of spikes when the trial pattern was presented (Fig. 1-2). Significantly tuned neurons (rate code) were defined as those that survived 3 statistical tests of visual responsiveness, at level of P < 0.05 simultaneously. One-way analysis of variance (ANOVA) tested whether a neuron responded differently (had a different mean firing count) to all the individual stimuli (e.g., gratings of a specific angle) in a stimulus type^106^. Similarly, Kruskal-Wallis tests^27^ were applied to assess such differential activation across individual stimuli (different median firing counts). The third test was a chi-square test of independence^28^; briefly, chi-square statistics 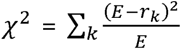 measures the total amount of unexpectedness across all *k* stimuli, between the empirical response (empirical trial averaged firing count *r_k_*) relative to the expected response, i.e. a neuron responds equally strongly to all individual stimuli, 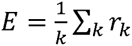 (same number of trials per stimulus). The null distribution for calculating the P value of the test-statistics was simulated by shuffling the stimulus identity and recalculating *r_k_*, resulting χ^2^ 1000 times. Visual responsiveness was assessed for each neuron for each stimulus type individually.

### Visual decoding

Regularized linear discriminant analysis was consistently used. Either the total spike count (rate code) or temporal components during stimulus presentation were used as inputs. The decoding performance was defined as the difference between the true angle and decoded angle for DG and RFG in Fig. 5, or the portion of correctly/incorrectly classified trials for all other figures. For RFG stimuli, only 25 (5 x 5) spatially non-overlapping (sampling every other pattern in both azimuth and elevation axes) stimuli out of the 81 patterns shown were used for decoding. Overall decoding error in any day was derived for each stimulus condition (e.g., 0-degree drifting gratings) and each mouse separately. Chronic decoding performance was evaluated based on these same mouse-stimulus pairs across days. Decoding drift index as defined as (*Acc_wd_* − *Acc_bd_*)/(*Acc_wd_* + *Acc_bd_*), where *Acc_wd_* or Accuracy within-day indicates the accuracy of 1/7 held out trials in the first 7 days when the decoder was trained. *Acc_bd_* or Accuracy between-day indicates the decoding accuracy on a future testing day. For DG and RFG stimuli, decoding angular error (the smaller the better) were converted to “accuracy” (the larger the better) by subtracting from maximal possible angular error any pairs of true vs decoded stimuli could obtain.

### Tuning reliability and selectivity

Similar to past literature^37^, the reliability of a neuron’s rate code or temporal code to a particular stimulus type on any given day was defined as the correlation coefficient between its tuning function derived from one trial versus the mean tuning function computed from the rest of the trials, averaged over all such single trials. Tuning selectivity measures the sparsity^38^ of neural responses to all stimuli in a stimulus type. This measurement of stimulus selectivity has been shown to be generalizable to all stimulus types, capable of considering the influence of the number of unique stimuli per stimulus type and offers unique insight beyond conventional spareness measures. A total of 1000 equally spaced firing count threshold values *T*_1_ ∼ *T*_1000_ were chosen to span the minimal to maximal trial-averaged firing count *r_k_* across all *k* stimuli. S was then defined as 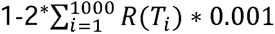 * 0.001, where *R*(*T_i_*) is the fraction of stimulus whose *r_k_* is greater than *T*_i_. We took the absolute value of S as the final selectivity value. When a unit fires equally to all stimuli, S = 0, and when it fires predominantly to one stimulus or fires in response to all but one stimulus, S approaches 1.

### Classification of cell types

Using metrics exported by ^39^, we defined narrow interneurons as having trough to peak width < 0.48 ms and wide interneurons as having trough to peak width > 0.48 ms and ACG rise time > 6 ms. Both narrow and wide interneurons were later grouped into one interneuron class.

### Temporal Components

LDA temporal components were identified for each neuron and under each of the 4 types of stimuli separately. The spiking timestamps during stimulus-on period of a neuron were smoothed with a 49 ms moving mean window. Trials from all days (Fig. 4) or the first 7 days (Fig. 5) for a particular stimulus type was then pooled before fitting a cross-validated, regularized LDA model (searching for best gamma with Bayesian optimization using MATLAB’s fitcdiscr function) to classify stimuli identity, e.g., which of the 16 directions for DG stimuli. To reduce the information overlap between rate codes and temporal codes, we discouraged LDA from finding temporally flat weights by normalizing each trial with the area under the peristimulus time histogram before fitting. The final LDA axes were extracted using a separate model fitted with the optimal parameter found in the cross-validation process. LDA is a supervised dimensionality reduction technique that finds axes maximizing the linear separability of data based on class labels^107^. LDA computes eigenvectors (axes) corresponding to the eigenvalues of the matrix (*S_W_*)^−1^ *S_B_* where *S_W_* is the within-class scatter matrix, representing the variance within each class. *S_B_* is the between-class scatter matrix, capturing the variance of class means relative to the global mean. However, when working with high-dimensional data, LDA can suffer from overfitting^108^. To address this, a regularization term is added to stabilize the computation. Instead of directly computing eigenvectors from (*S_W_*)^−1^ *S_B_*, they are now computed from 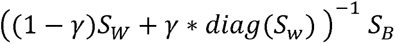 where *diag*() is the diagonal matrix operation, *y* is the regularization factor between 0 and 1. (determined through cross-validation).To compare the stability of rate-code based tuning and temporal components-based tuning, for each recording day and each neuron, 50% of the trials were randomly selected and averaged to form the tuning curve for between-session similarity comparisons. For within-day cases, the 50% sampled trials were averaged and compared to the average of the complementary 50% of the trials. Sampling was repeated 30 times, and the results were further averaged. Tuning curve similarities between days (tuning stability) were quantified with Pearson’s Correlation Coefficient (CC) before further converting to the representational drift index (*CC_wd_* − *CC_bd_*)/(*CC_wd_* + *CC_bd_*) as in ^10^.

### Population dynamics

We projected the population firing time course into a low-dimensional stimulus coding space with cross-validated LDA decoders^44^, pooling all trials across 15 days. Cross-validated decoders were constructed with populational spike count using a 90 ms integration window at 160 ms after stimulus onset (from 70 ms to 160 ms) to ensure that different stimulus categories were readily decodable. Four different decoders were fitted to separately classify the orientation, direction of drifting gratings and orientation, and spatial frequency of static gratings. The decoder optimization procedure was similar to that described in the visual decoding section. The corresponding decoder weights were then used to project neural data from 30 ms to 180 ms in 10 ms increments. To determine the feature separation timing between different stimulus categories (such as orientations versus directions), the pairwise inter-condition distance matrices were calculated with upper-triangle elements averaged at each time bin. The separation timing was then defined as the time (after interpolation) at which the inter-condition separation reached halfway to the maximal value from the initial starting distance (Supplementary Fig6 for graphical explanation).

### Representation in UMAP space

As reported in past literature, V1 populations have high-dimensional geometry, at least when the dimension is linearly reduced^50^. We thus followed multiple literature to use nonlinear neighbor embedding methods^11, 49^ when reducing the dimension of the neural populational responses. Even/odd trials-averaged population firing counts for all stimuli in a stimulus type for 15 days were pooled to be jointly dimension-reduced. (e.g., 16 grating directions x 15 days x even/odd trials = 480 samples). Uniform Manifold Approximation and Projection^51, 74^ (UMAP) was performed for each stimulus type separately with the following parameters:

DG: correlation distance, 2 dimensions, 149 neighbors, 0.9 minimum distance
SG: Euclidean distance, 2 dimensions, 134 neighbors, 0.7 minimum distance
RFG: cosine distance, 2 dimensions, 134 neighbors, 0.9 minimum distance
NI: correlation distance, 2 dimensions, 119 neighbors, 0.9 minimum distance

When quantifying if the representations of the same stimulus conditions were closer to themselves across days than to different stimuli within days in the dimension reduced space, all distances computed were normalized by the mean of within-day distances (even versus odd trials) for the same conditions across all days. When quantifying separation between the same stimulus across days against the nearest five different stimuli within days, we used 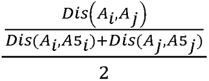, for day pair *i* and *J* of stimulus *A*, where *Dis*(*A_i_, A_j_*) is the average distance between 4 pairs (even/odd trial means x two days) of within stimulus between day Euclidean distances. *Dis*(*A_i_, A*5*_i_*) is the average distance between 20 pairs (even/odd trial means x 5 neighbors and their even/odd trial means) of between stimulus Euclidean distances within day *i*. UMAP generates low-dimensional embeddings from the original high-dimensional space, preserving the probability of connections (transformed from distance measures) between data points and their nearest neighbors. While a single distance measurement in the UMAP space is unitless and less meaningful on its own, we believe the distance ratio between the same stimulus across days and nearby competing stimuli is more informative. Conceptually, this ratio resembles an odds ratio in statistics. We posit that when representational drift is low, stimulus representations from one day are more likely to be connected to the same stimulus from another day rather than to competing stimuli.

### Preferred spatial frequency/orientation

For each neuron, the preferred spatial frequency/orientation of SG stimuli was defined as the trial-averaged firing count weighted spatial frequency/orientation, marginalizing over orientation/spatial frequency respectively.

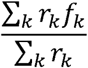

### Functional Connectivity

We followed published methods carefully to calculate the jitter-corrected cross-correlogram^28, 57^ CCG as 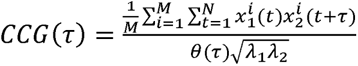 between the spike trains of pairs of units 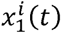 and 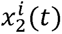. CCGs were derived individually for each recording day and separately for each stimulus type (DG, SG, RFG, NI) at 1 ms temporal resolution. Only unit pairs whose mean firing rate *λ*_1_, *λ*_2_ were both higher than 2 Hz for that specific stimulation type were included in the calculation. Only the stimulus-on period was used for calculation. The CCGs were then normalized by the geometric mean firing rate of the two units as well as a triangular function *θ(τ)* to correct overlapping sliding window edges. The CCG was then averaged across all *M* repeats of the same stimulus *i*. Individually normalized CCGs were calculated for each stimulus pattern (e.g., drifting grating directions, static grating orientations *x* spatial frequencies) separately before averaging across all available patterns of that stimulus type. A peristimulus time histogram-preserving jittered version of the CCG was computed and subtracted from the original CCG to generate the final CCG. A pair was considered significantly functionally connected if there was a peak in the −10 ms to 10 ms region of the correlogram, and this peak was above six standard deviations of cross-correlogram flank (defined as −112 ms∼62 ms, together with 62 ms∼112 ms). CCG peak amplitude was defined as the connectivity strength, and the temporal location of the peak is defined as the lag.

### Population Coupling

Population coupling^54, 55^ between a specific neuron and the rest of the neuronal population is the correlation coefficient between per-trial-based the response (firing count) of that neuron with the summed response of the rest of the population in the same animal. All trials of a specific stimulus type were directly concatenated without considering stimulus identity for this calculation. Population coupling was computed for each neuron-stimulus type combination.

### Spike-Triggered Population Activity

A separate measure that describes the neuron-to-population coupling is spike-triggered population activity^54, 60^. The spike train of a target neuron and the summed spike train of the rest of the neuronal population were summarized on a per-trial basis at 1 ms resolution. For each trial, the target neuron triggered population firing histogram of the same trial was calculated at different lag. A shuffled version of the spike-triggered histogram was computed using population activity from different trials (permuted 100 times) and averaged across permutations. After subtracting the shuffled histogram, the per-trial-based spike-triggered population firing histogram was averaged across trials. The strength of the coupling was defined as the final histogram value at the 0 ms time lag.

### Statistics

All the statistical analyses were performed in MATLAB (MathWorks, MA). All tests were two-sided. Kruskal-Wallis, ANOVA and Chi-Square test of independence were used to test the visual responsiveness of the units as reported in the methods section. Normality was assumed but not explicitly tested whenever applicable. To quantify if units were more like themselves across days than other units in the same probe within days, for both biophysical and tuning properties, independent t-tests were used. They were also used to compare functional tuning stability/reliability between cell types. Welch’s paired t-tests were employed for the pairwise comparisons between the portions of recovered user responses for different tracking schemes for units from each electrode shank. It was also used to compare firing rate versus component-based single neuron decoding accuracy. The superiority of using tracked (paired sample) objects rather than unpaired population measures to measure population stability has been discussed previously^61^. To quantify such longitudinal stability measures for tracked (paired sample for longitudinal statistical study design) animals, probes, single units, single unit-pairs, and mouse-pattern pairs, we used linear mixed effect models^10^. Objects tracked for less than 3 days were typically excluded from such analysis following the 3 time points minimal guidance in^42^. A recent study also reported that true stability could be confounded by a short tracking duration^58^. To investigate individual unit, individual unit pair, individual animal, individual probe’s evolving direction and magnitude for any measures of stability (changes over time), a separate linear regression over time model was fitted for each member in the population. Bonferroni’s correction for multiple comparisons was applied. To determine the significance of the differences between two correlation coefficients, the MSTATS toolbox was used.

### Data Availability

Data supporting the main conclusions (e.g., neuron population spiking timestamps across different trials X stimulus types X recording days) will be available upon publication.

### Code Availability

- Mountainsort4 for spike sorting is available at https://github.com/magland/ml_ms4alg
- LDA decoders are available with MATLAB functions “fitcdiscr”, sample code for temporal components fitting will be available upon publication.
- Linear mixed effect, multiple linear regressions were performed with MATLAB function “fitlm,” “fitlme.”
- To test different in correlation coefficient, MSTATS toolbox was used https://www.mathworks.com/matlabcentral/fileexchange/47235-mstats-a-random-collection-of-statistical-functions.
- UMAP for dimension reduction is available at https://umap-learn.readthedocs.io/en/latest/
- Cell explorer for ACG, CCG, cell type classification, is available at https://cellexplorer.org/
- Jitter corrected CCG is available at https://github.com/jiaxx/modular_network
- Shuffle corrected spike triggered population activity is available at https://github.com/VeronikaKoren/struct_pop_public
- Dense neural network used for validation of labelling consistency is available at https://github.com/KordingLab/Neural_Decoding
- DeeplabCut for animal pose labelling is available at https://github.com/DeepLabCut/DeepLabCut
- YOLO for animal pose labelling is available at https://docs.ultralytics.com/

## Acknowledgements

We thank Dr. Cristopher Niell, Dr. Nuo Li for feedback. We thank Dr. Valentin Dragoi for suggestions on population coupling analysis. We thank Dr. Ngoc M. Tran for input on tracking the same neurons. We thank NIH R01NS102917 (to C.X.), U01NS115588 (to C.X.), R01NS109361 (to L.L.) and Rice University internal awards (L.L. and C.X.) for funding.

## Author Contributions Statement

Contributions: Conceptualization, C.X., H.Z., L.L.; Supervision, C.X., L.L.; Investigation and formal analyses, H.Z. with input from C.X., A.S.T., L.L.; Methodology - electrode preparation: P.Z., F.H. supervised by C.X.; Methodology - surgery: F.H. with input from C.X., L.L.; Methodology - experiment setup: H.Z., S.P. with input from C.X., A.S.T.; Visualization H.Z., F.H. with input from C.X., L.L.; Writing - original draft, H.Z., L.L., C.X.; Writing - review & editing, L.L., C.X., A.S.T., H.Z. all authors reviewed the manuscript; Funding acquisition and project administration, C.X., L.L., A.S.T.; Resources, all authors.

Lead contact: C.X.

These authors jointly supervised the work: C.X., L.L.

These authors contributed equally: H.Z., F.H.

Present Address:

Fei He. Shanghai Institute of Optics and Fine Mechanics, Chinese Academy of Sciences, Shanghai, China

