## Supplemental Information for "Temporal coding carries more stable cortical visual representations than firing rate over time"

**Affiliations:**

### Supplemental Discussion

Our unit tracking algorithm outperformed mutual nearest neighbor[1] alone in the longevity of tracking. To evaluate manual labeling consistency, we predicted manual decisions with linear and non-linear models (Supplementary Fig20c-d). Extending from single-channel-based work[2], we found that 97.31 ± 0.29% (mean + s.e.m.) of manual decisions from a given probe could be recovered by a deep learning (DL) model trained on other probes using multi-channel features. In comparison, a discriminant analysis-based[3] linear model recovered 96.39 ± 0.36%, indicating that the labeling was objectively reproducible with sophisticated models(Supplementary Fig20e)

We investigated whether the extracted temporal components across different neurons have universal or diverse profiles. We visualized shapes of top three components pooled from all single units with UMAP embedding (Supplementary Fig5b) when mice viewed drifting grating stimuli. Two somewhat separable clusters were observed, representing temporal integration (where either positive or negative peak weight dominates) and temporal contrast (with near equal magnitude of peak positive versus negative weight), respectively. Within each cluster, the peak activation timing of components tiled the entire stimulus duration, highlighting the diversity of stimulus-evoked response time courses across the neuronal population. While the major shape clusters seemed to align with a report on gaze shift induced temporal dynamics[4], having either monophasic or biphasic response, we observed a much more diverse distribution of component peak timing with more blurred boundaries to define “early” or “late” activation clusters, unless “early” is defined by sooner than 20 ms (the black/dark blue clusters in the peak timing panel) , which is typically before any stimulus information reaches visual cortices[5]. Such differences might be due to different dimension reduction techniques for identifying temporal components, which could be the subject of future research.

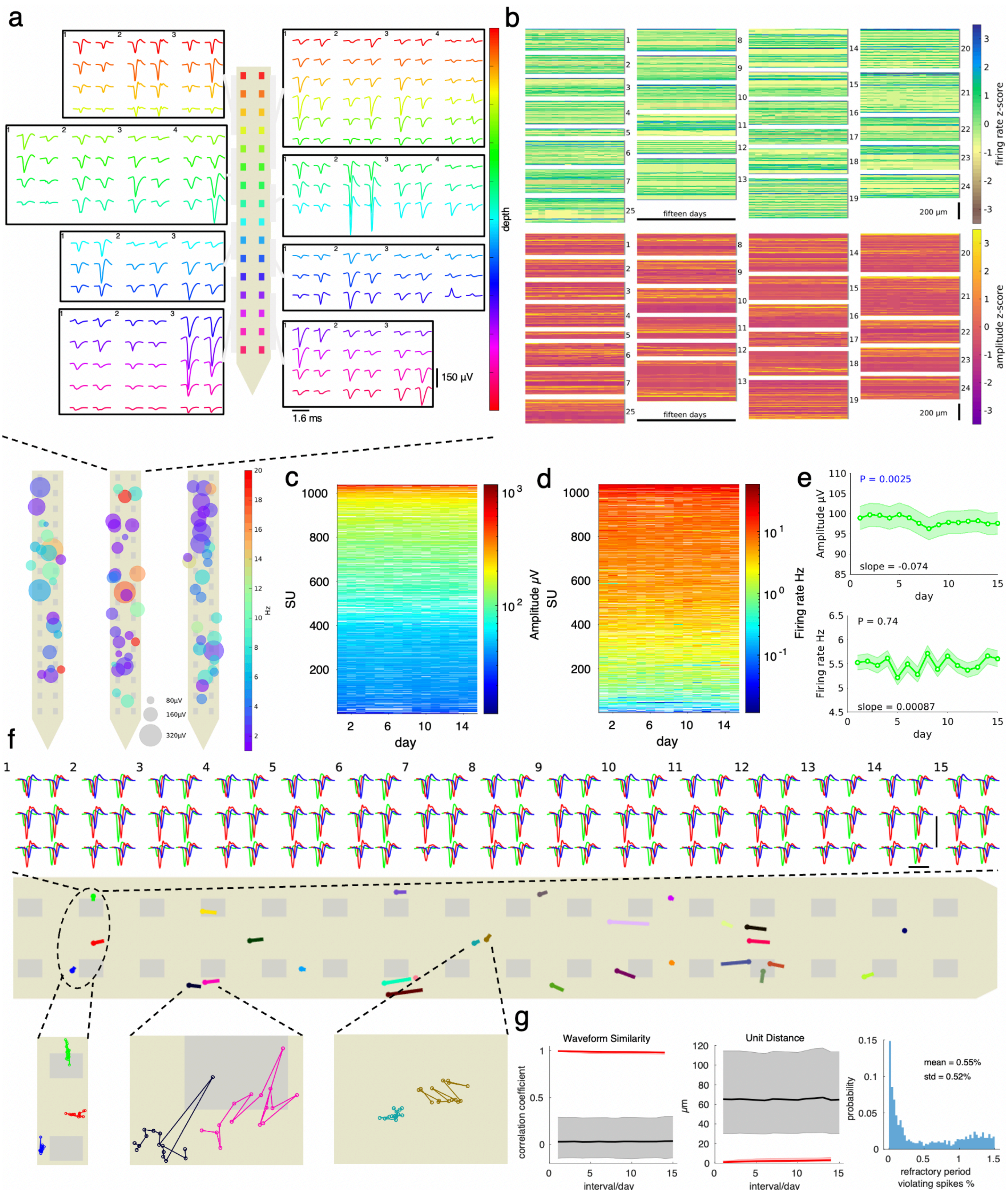

**Supplementary Fig1 Stability of overall recording and stability of tracked units' longitudinal biophysical signatures, related to Fig. 1.**

**a.** The closely spaced recording sites on NETs facilitated the separation of spatially adjacent neurons and allowed estimation of neuron locations relative to the recording sites (Top: Multi-channel spike waveforms from one NET recording with 28 units (the middle probe in the bottom panel). Each unit, denoted by number, spanned across multiple spatially adjacent channels. Channels are color-coded by depth along the probe (inter-electrode distance: 20 μm). Bottom: Representative recordings from 3 NETs, showing many single units and their estimated locations. Amplitude is coded in size and firing rate coded in color. **b.** Minimal electrode-tissue movement was observed. Smoothed depth histogram of firing rate (top) and amplitude (bottom) from all units across time. All 25 NETs from n = 5 animals were vertically stacked. No visible drift along depth from day 1 to day 15. **c-e.** Tracked units had stable amplitude (c) and firing rate (d) across sessions. Showing heat map (c-d) of all single units tracked > 2 days from all mice and corresponding population mean ± s.e.m. over time (e). Linear mixed effect model, time as fixed effect, units as random effect. (n=1037 single units appearing more than 2 days) **f.** Representative tracking of single units in one probe showed high waveforms similarity (top) and minimal position drift (bottom) across 15 days. scale bar vertical: 150 μV. Horizontal: 1.6 ms. Zoom-in images show the day-to-day movement between trackable neurons and recording sites (gray rectangles, inter-electrode distance 20 μm), with positions of each day marked by circles and of adjacent days connected by lines. **g.** Single units' waveforms (left), locations (middle) were more like themselves across sessions than other simultaneously recorded units from the same probe. Showing median, 25 percentiles and 75 percentiles. $p<0.0001$ with values reported in Supplementary Table 1, independent t-test. n=757, 311845 same/different unit pairs respectively at 14-day interval. Right: Probability density of refractory period violations for single units included in the study. (refractory period is defined as an inter-spike interval <2 ms) See Supplementary Table 1 for additional reporting on sample size and statistics.

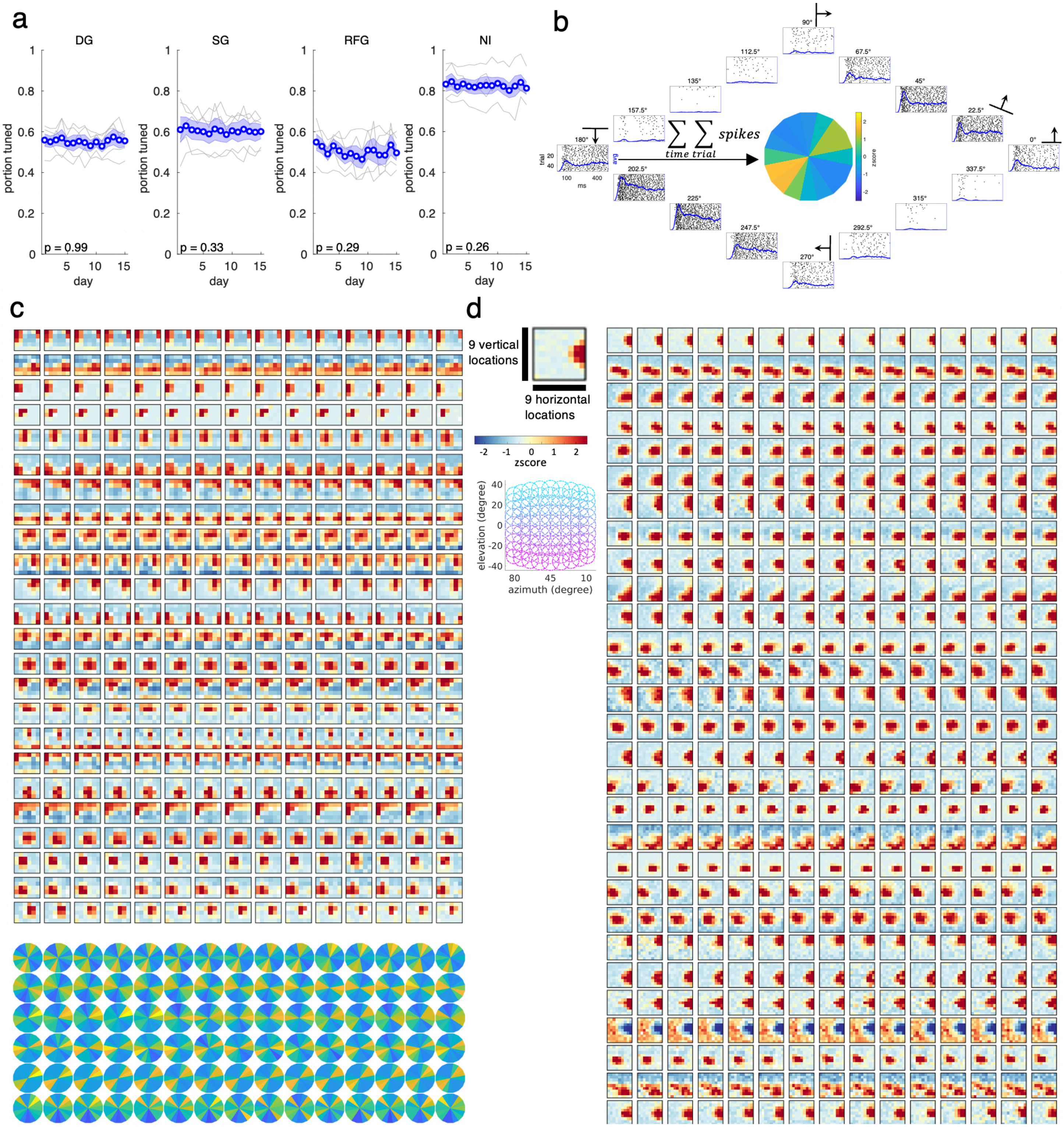


**Supplementary Fig2 Stable functional tuning across 15 days for diverse stimulus types, related to Fig. 1.**

**a.** The fraction of neurons tuned to stimulus remained stable ($p > 0.05$) for all four stimulus types, linear mixed effect with time as fixed effect, individual mouse as random effect (n=5) **b.** Explanation of firing rate-based tuning curve calculations related to panel c, d, and Fig. 1 f-h. A tuning curve of a neuron was formed by summing the number of spikes across trials for any given condition while z-scored across conditions. Rasters (dots) show spike timing for each stimulation trial aligned to visual stimulation onset for one neuron in one day. Trials (n=50) were collected during drifting grating stimuli, with 16 grating moving directions. Smoothed trial averaged spiking time courses (blue, min-max normalized across conditions) were added as visual guides to compare inter-stimulus differences. **c-d**. Units showed stable tuning curves across 15 days in four stimulus types, related to Fig.1, showing additional neurons for static gratings (c, top), drifting gratings (c, bottom), and representative neurons during receptive field mapping (d). The exact elevation and azimuth mapped, and sizes of the Gabor patterns are presented as gridded circle plots. See Supplementary Table 1 for additional reporting on sample size and statistics.

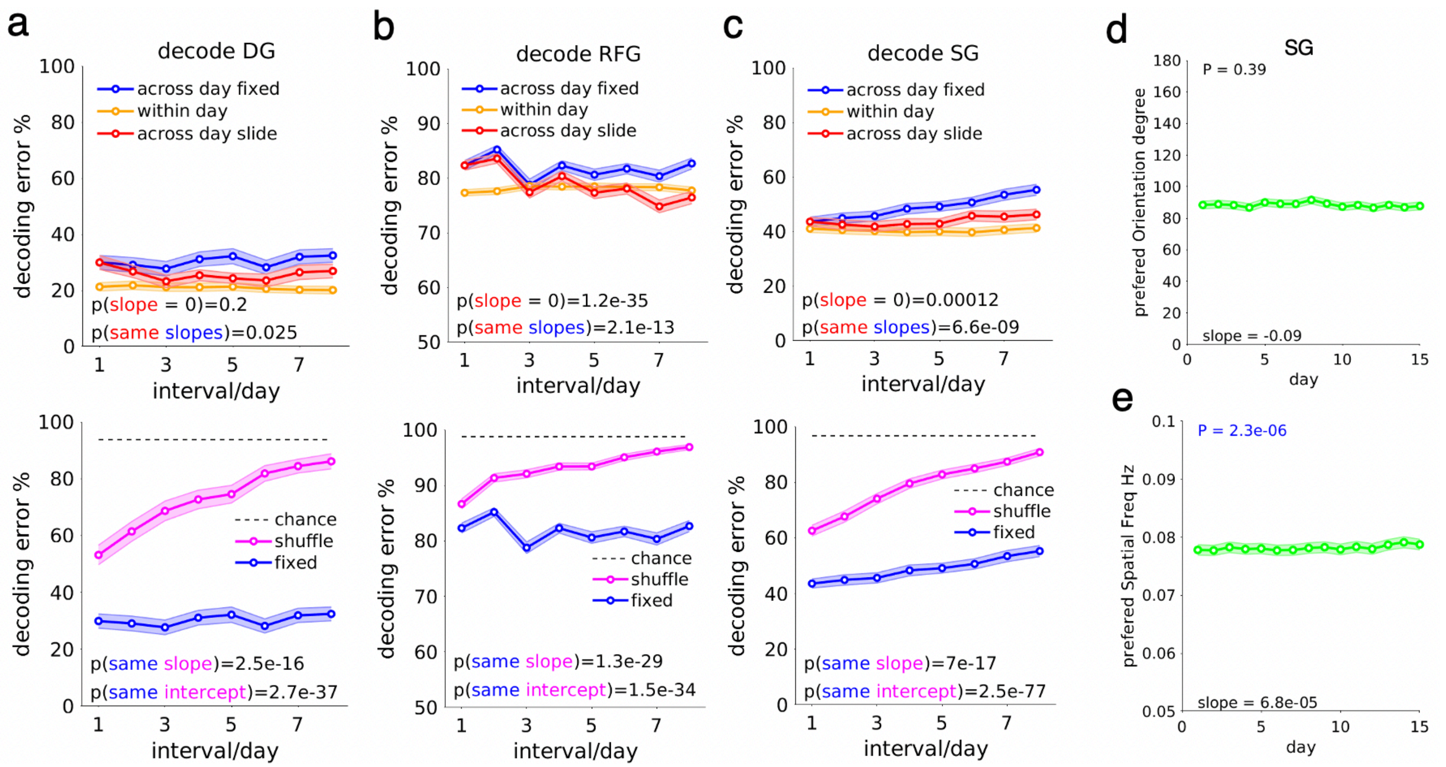


**Supplementary Fig3 Drift in visual representations cannot be trivially attributed to unreliable tracking of neuron identity, related to Fig. 1**

**a-c.** The decoder with shuffled neuron identity approached chance at a higher rate than the fixed-in-time decoder across all stimuli (DG: drifting gratings; SG: static gratings; RFG: receptive field Gabors). Sliding decoders reduced the rate of error increase compared to the fixed decoder for all stimuli. Sliding decoders did not have an error increase and approached the within-day decoder for all stimuli but SG, which may be related to an inherent increase in neuronal preferred spatial frequencies over time[6]. Plots are in mean + s.e.m. Linear mixed effect, time as fixed effect, decoding method (shuffle vs fixed-in time / sliding vs fixed-in time)) as fixed effect individual animal-stimulus pairs as random effect (80,405,150 animal-image pairs for DG, RFG, SG, respectively). **d-e.** Preferred orientation (d) did not change yet preferred spatial frequency (e) increased during static grating stimuli. The plots were derived from all tuned single neurons that appeared for 15 days (n=478) from all mice. Plots are in mean + s.e.m. linear mixed effect model (LME), time as fixed effect, individual neuron as random effect. See Supplementary Table 1 for additional reporting on sample size and statistics.

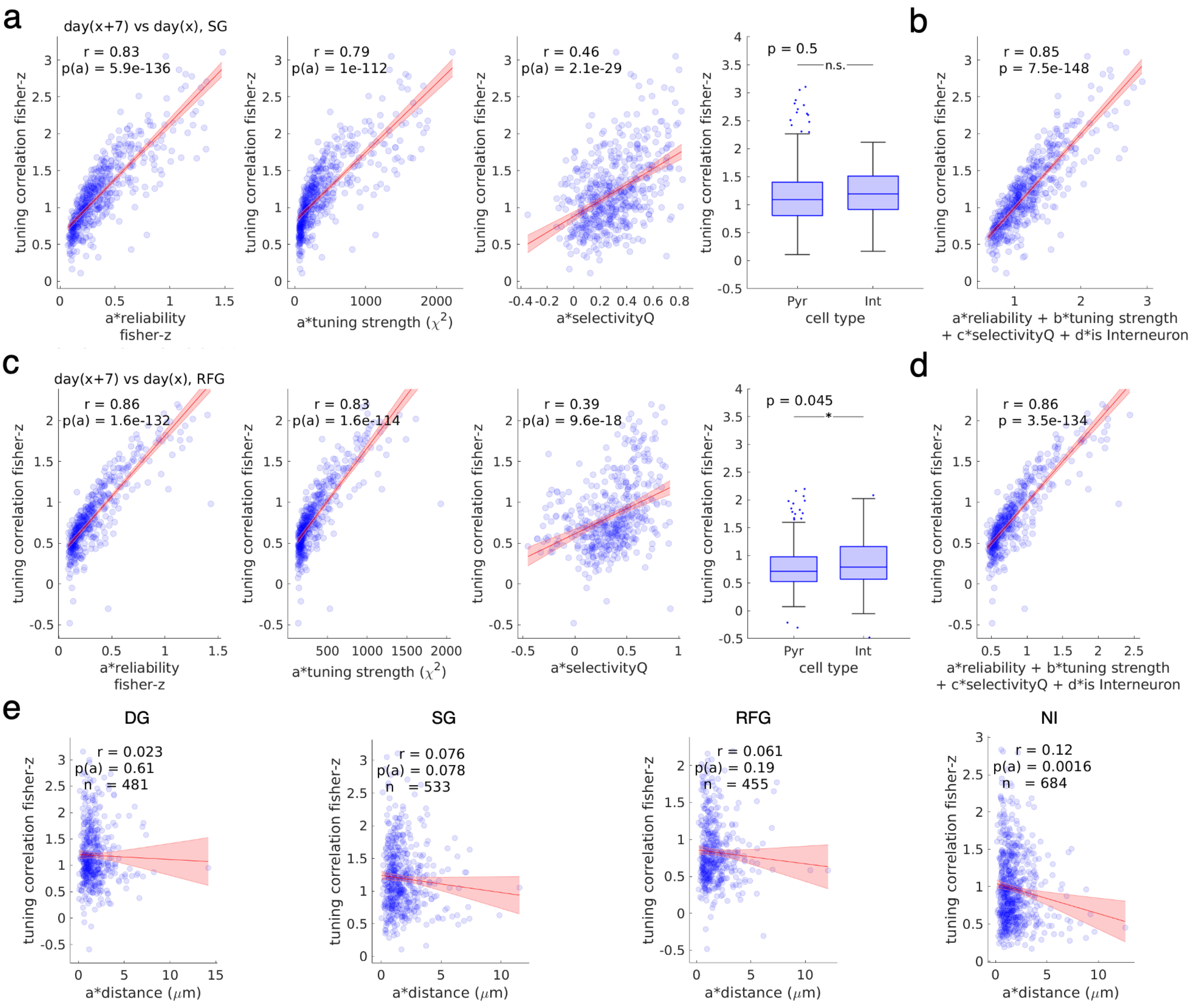


**Supplementary Fig4 Representational stability of individual neurons strongly correlates with tuning reliability, related to Fig. 2.**

**a**. Tuning similarity after seven days for static gratings was strongly correlated with tuning reliability (1st column), moderately correlated with tuning strength (Kruskal Wallis's test statistics, 2nd column) and stimulus selectivity (3rd column), while being weakly correlated with cell type (independent t-test, 4th column). Red line: linear regression fit with 95% CI, text: correlation coefficient, significance level. Correlation coefficient was Fisher Z-transformed. N=533 stimulus-tuned single units appearing > 2 days and reappeared after 7 days,(338 pyramidal neurons, 195 interneurons).

**b**. Reliability, tuning strength, selectivity, and cell types could jointly explain tuning similarity after 7 days. Red line: linear regression fit with 95% CI, text: correlation coefficient, significance level. **c-d**. As in b. but for receptive field Gabors stimuli. N=455 neurons (277 pyramidal neurons, 178 interneurons). **e**. Unit physical location changes marginally if significantly explained tuning stability for the four stimuli (DG: drifting gratings; SG: static gratings; RFG: receptive field Gabors; NI: natural images). Box plots are in median, 25 to 75 percentiles. Whiskers represent 1.5-fold interquartile range below Q1 or above Q3. Outliers are indicated in scatters. See Supplementary Table 1 for additional reporting on sample size and statistics.

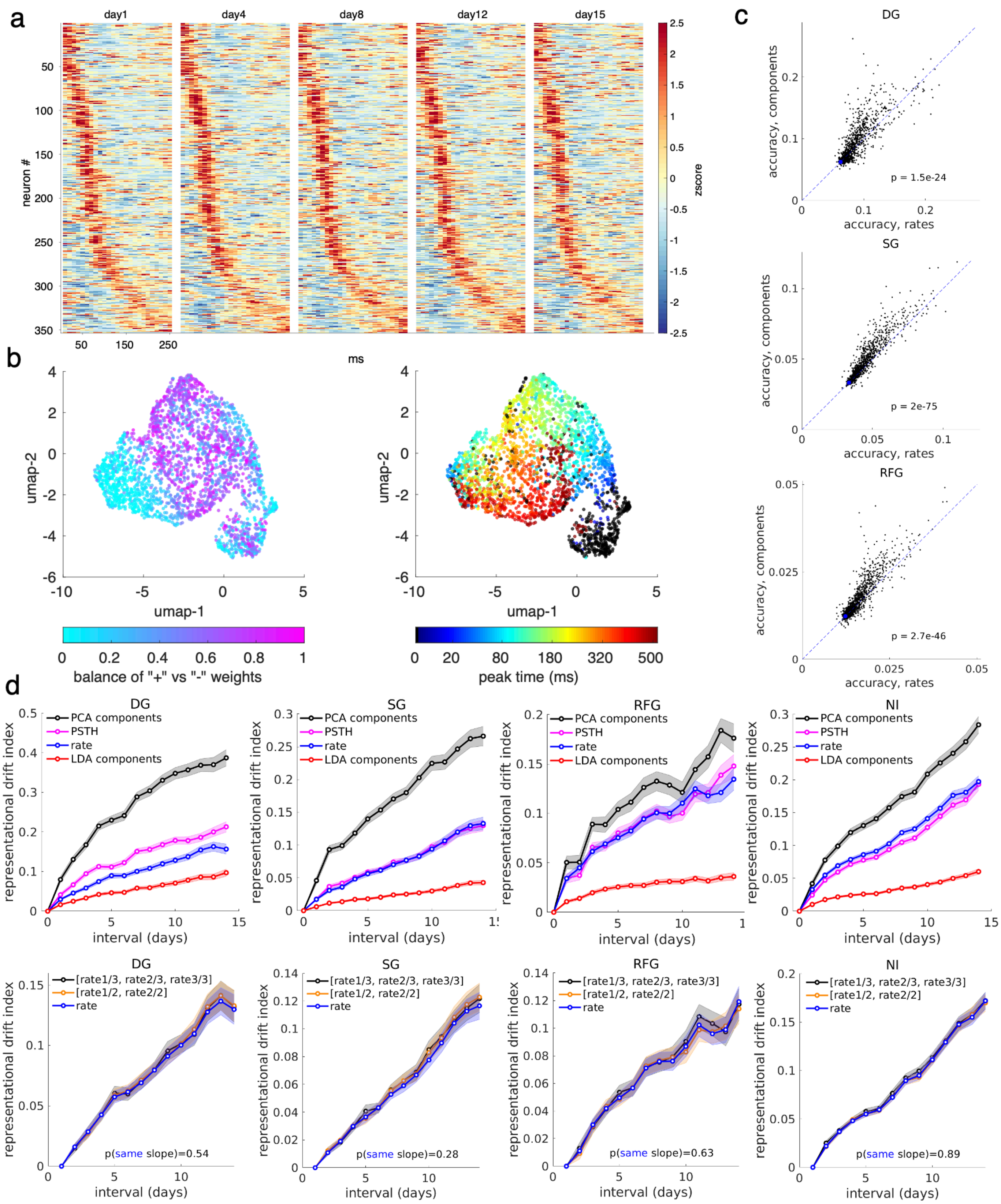


**Supplementary Fig5 Additional information on extracting temporal components from visual evoked spiking time courses, related to Fig. 3 and Fig. 4**

**a.** The stimulus-evoked spiking time course to a single, specific nature image of a population of neurons across multiple days. Neurons exhibited diverse temporal response profiles to this stimulus. Neurons were pooled from 5 mice over 7 different experiment days. Neurons that appeared for all 15 days and tuned to natural images were selected. The neurons were sorted by maximum activation time and were not resorted between days. This further illustrates that the temporal profile might remain stable across days. **b.** Components differed in the balance of positive versus negative weights in time (left) and in peak timing (right). UMAP embedding of temporal components during drifting grating (first three components pooled) from all single neurons collected across all mice, color-coded by components positive/negative weight balance ratio (left, 0 means imbalance), or component absolute peak timing (right) within the stimulus presentation period. Balance is defined by min(a, b) / max(a, b) where a and b are the magnitudes of cumulative (temporal integration) positive weights and negative weights of a temporal component respectively. **c.** As in Fig. 4d, temporal components achieved higher single neuron decoding accuracy than firing rates, n=1204 single units. Shown are the results for the other three stimuli: drifting gratings (top, chance level 1/16), static gratings (middle, chance level 1/30) and receptive field Gabors (bottom, chance level 1/81) **d.** Merely increasing the dimension of the tuning curve did not necessarily lead to increase tuning similarity across time for all four stimulus types (DG: drifting gratings; SG: static gratings; RFG: receptive field Gabors; NI: natural images), as exemplified by 1. Using top 3 PCA components extracted from the PSTH or the PSTH themselves to form the tuning curves did not meaningfully reduced drift than rate coding based tuning curves (top row). and 2. dividing each day's trials into 2 halves (even vs odd trials) or 3 one-thirds (e.g.,1,4,7…; 2,5,8…; 3,6,9…) before calculating their mean firing rates, thus increasing the dimension of each day's firing rate-based tuning by 2X or 3X respectively (bottom row) did not lead to an decrease in tuning drift as exemplified by LDA temporal components in Fig. 4f. N = 481, 533, 455, 684 stimulus-tuned single units that reappeared after 7 days for DG,SG,RFG,NI respectively. See Supplementary Table 1 for additional reporting on sample size and statistics.

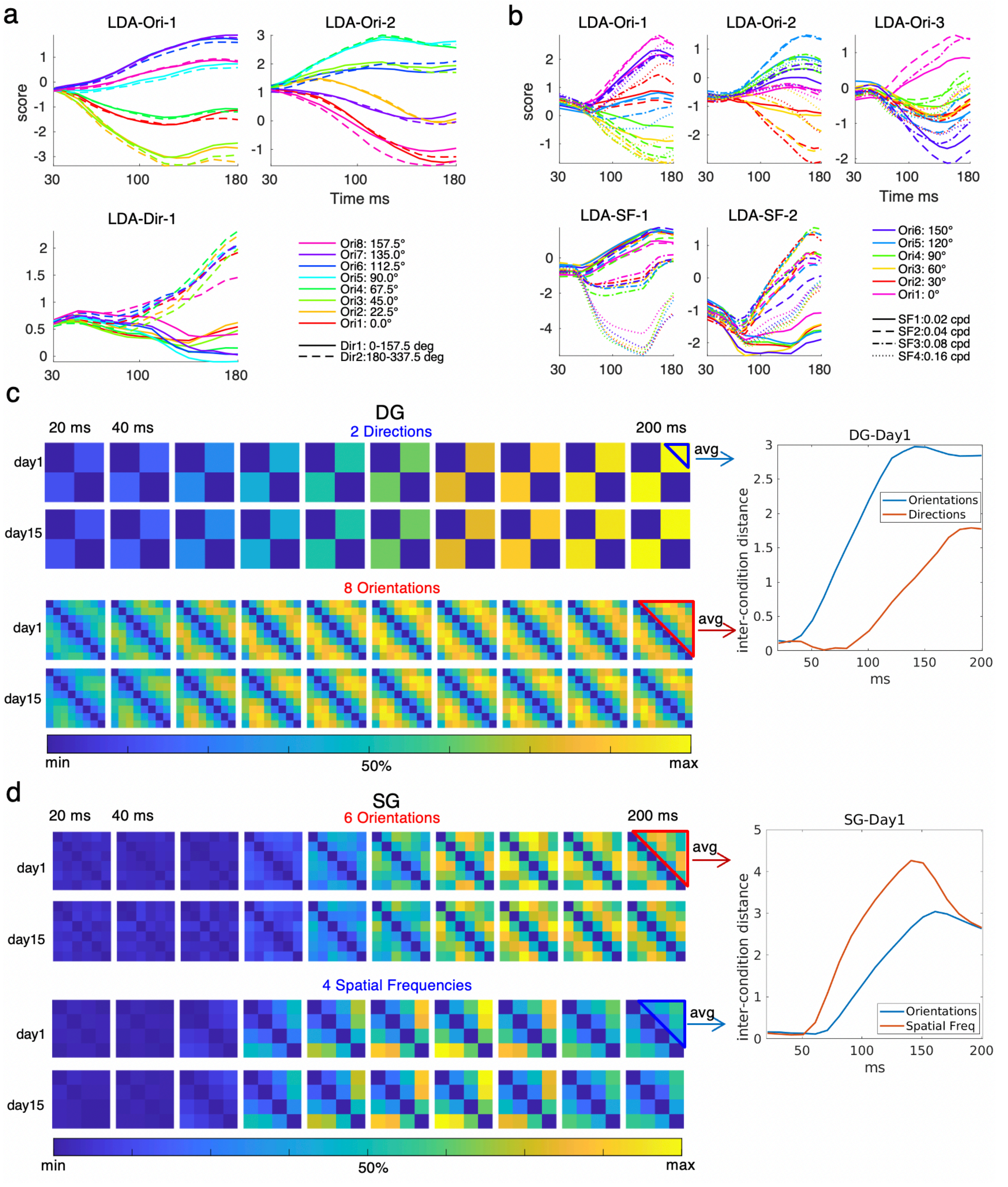


**Supplementary Fig6 Explanation of estimating the timing of stimulus categories separation from population dynamics, related to Fig. 3**

**a**. Per-component breakdown view of disentangled population representation dynamics during drifting gratings from an example animal. Traces represent 15-day averages in Fig. 3b, projected onto the first two linear discriminant orientation coding dimensions and the first direction coding axis. Moving angles of gratings with the same orientation but opposite directions (180° apart) were coded in similar colors. **b**. Same as in (a) but for static grating stimuli from Fig. 3c, projected onto the first three orientation coding dimensions and the first two spatial frequency coding axes. Stimulus orientations were coded in color, while spatial frequencies were coded in line style. **c**. Orientations were separated sooner than directions during drifting grating stimuli with consistent results across 15 days. Left: Between-condition distance matrix of population dynamics in two days for grating directions (top two rows) and grating orientations (bottom two rows) during the first 200 ms of stimulus presentation in one mouse. Coordinates of population dynamics were extracted from 1-day averaged traces as in Fig. 3b, using the first direction encoding LDA axis (top two rows), or the first two orientation encoding LDA axes (bottom two rows). The 8 orientations were derived from 16 initial angles, averaging across an angle and its corresponding angle + 180 degrees. The two directions were an angle versus (angle + 180 degrees). Euclidean distances were normalized by the maximum distance across time. Right: Evolution of average inter-stimulus distance. **d**. Spatial frequencies were separated sooner than orientations during static grating stimuli, with consistent results across 15 days. Same analysis as in (c) but repeated for static gratings; distance was calculated using the first three orientation coding axes and the first two spatial frequency coding axes.

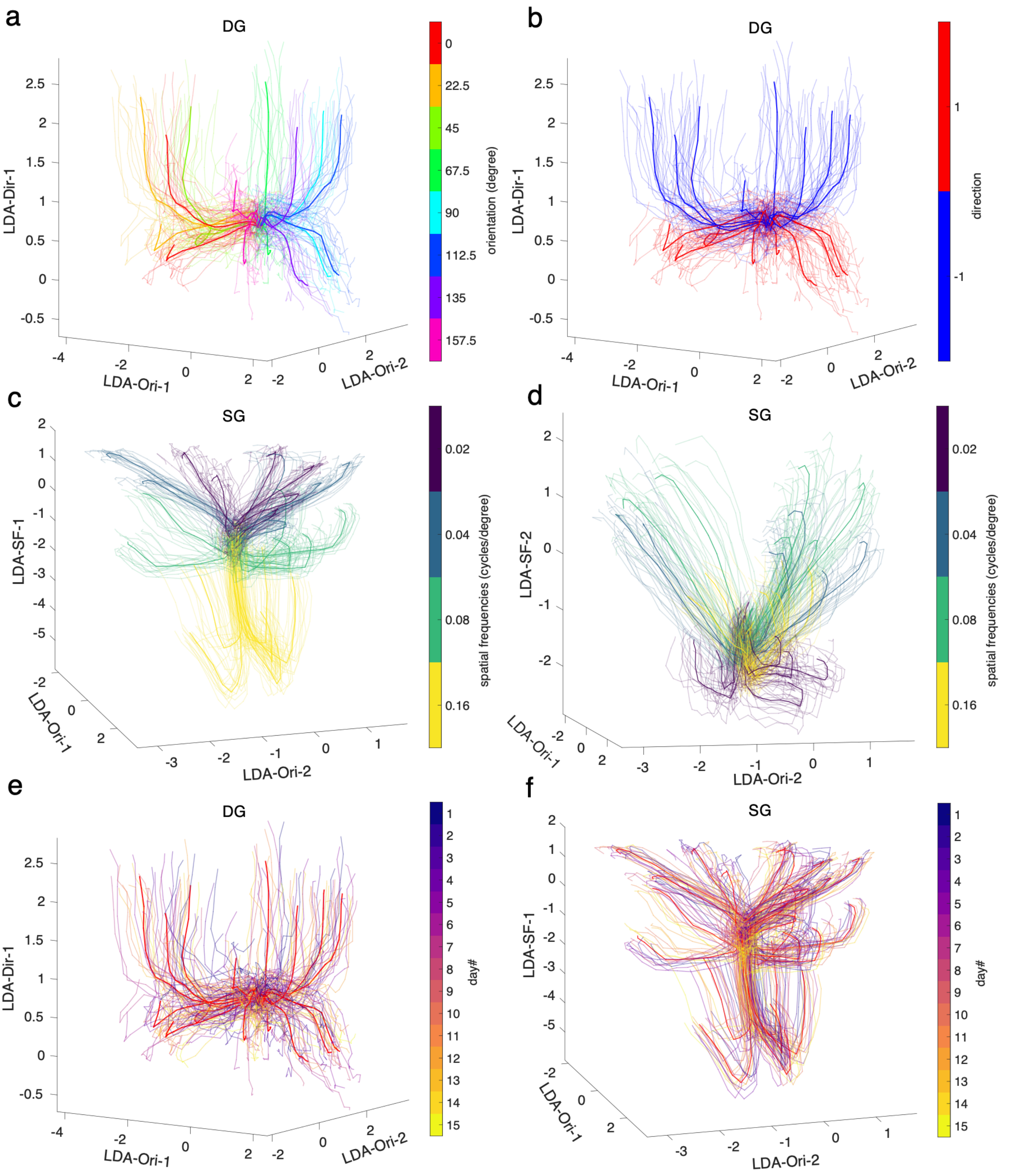


**Supplementary Fig7 Additional visualization on temporally resolved populational representation, related to Fig. 3**

**a-b**. Disentangled population representation dynamics during drifting gratings from an example animal. The plot shows daily trial-averaged (lighter lines) and 15-day-averaged (darker lines) traces, projected onto the first two linear discriminant orientation coding dimensions and the first direction coding axis. Orientations (a) or directions (b) were used to color-code the traces. **c-d**. Same as in (a) but for static grating stimuli, projected onto the first two orientation coding dimensions and the first (c) or second (d) spatial frequency coding axis. Stimulus spatial frequencies were coded in color. **e-f**. same as in (a) and (c) but color-coded by day number.

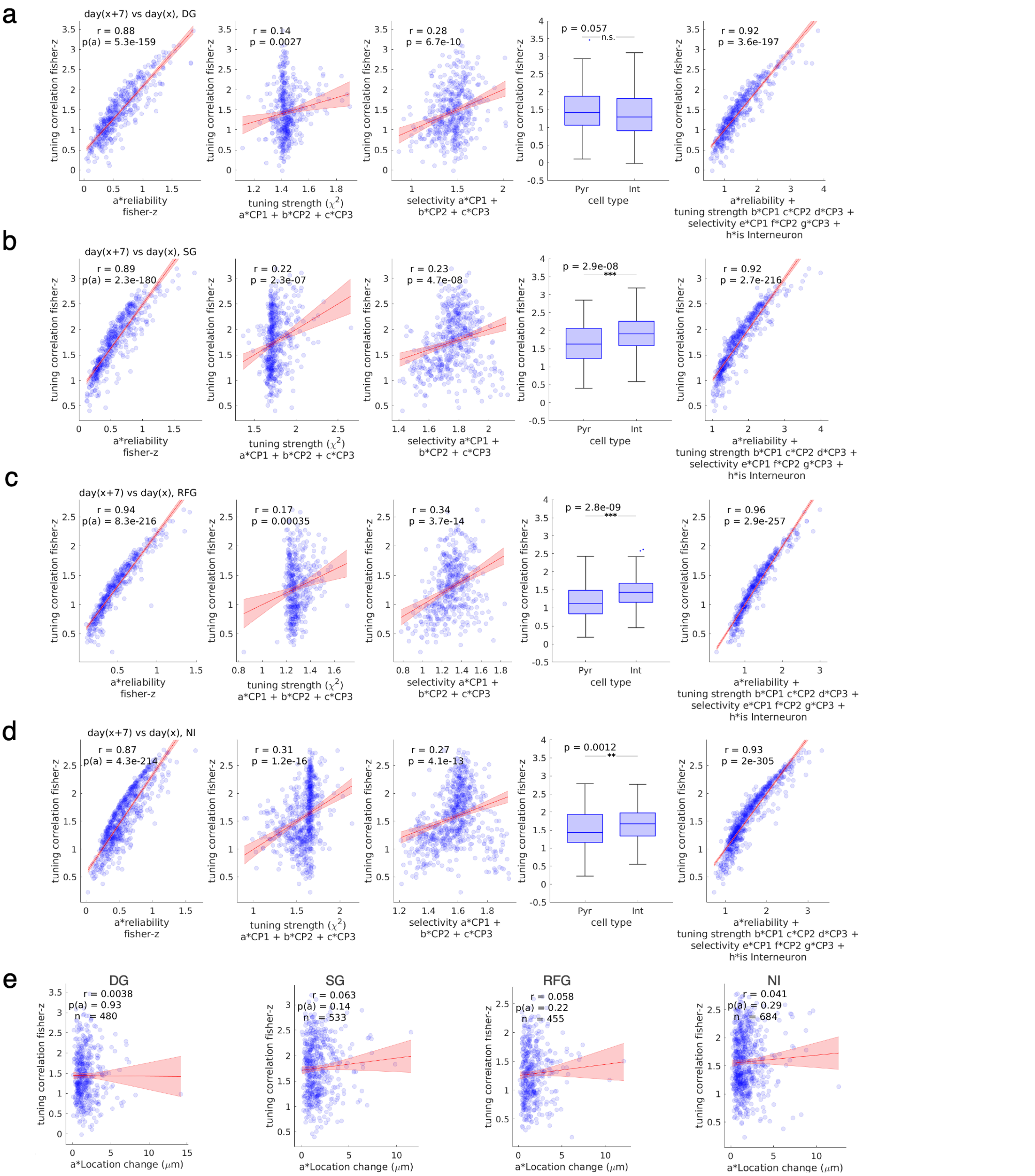


**Supplementary Fig8 Temporal components-based single neuron tuning stability also explained by tuning reliability, strength, and selectivity, related to Fig. 4**

**a-d.** Across all four stimulus types (rows, DG: drifting gratings; SG: static gratings; RFG: receptive field Gabors; NI: natural images) temporal components-based tuning similarity after 7 days strongly correlated with tuning reliability (1st column), weakly or moderately correlated with tuning strength (Kruskal Wallis's test statistics, 2nd column) and stimulus selectivity (3rd column). Interneurons exhibited higher components-based tuning stability for all stimuli except for drifting gratings (d, 4th column, independent t-test). Correlation coefficient was Fisher Z-transformed .Reliability, tuning strength, selectivity and cell type could jointly explain tuning similarity after 7 days (5th column). Red line: linear regression fit with 95% CI, text: correlation coefficient, significance level. DG: N=481 stimulus-tuned single units appearing > 2 days and reappeared after 7 days (299 pyramidal neurons, 181 interneurons). SG: N=533 neurons (338 pyramidal neurons, 195 interneurons). RFG: N=455 neurons (277 pyramidal neurons, 178 interneurons). NI: N=684 neurons (451 pyramidal neurons, 233 interneurons). **e.** Unit physical location changes did not significantly explain temporal components based tuning stability for the four stimulus types. Box plots are in median, 25 to 75 percentiles. Whiskers represent 1.5-fold interquartile range below Q1 or above Q3. Outliers are indicated in scatters. See Supplementary Table 1 for additional reporting on sample size and statistics

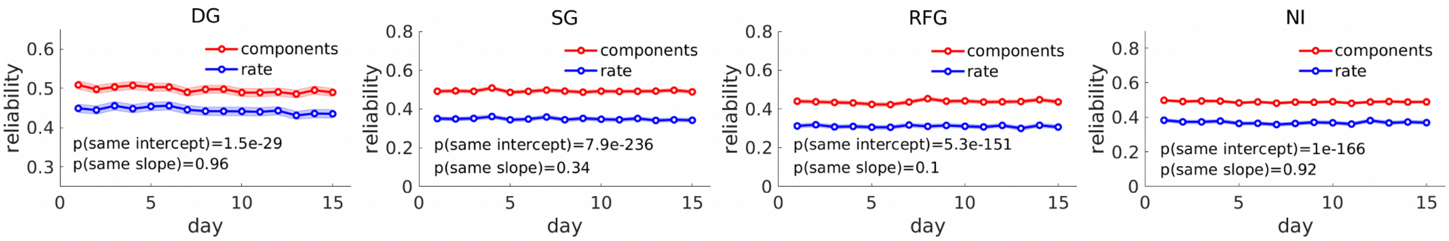


**Supplementary Fig9 Temporal code increased tuning reliability across all 4 stimulus types compared to firing rates, related to Fig. 4**

Tuning reliability increased across all 4 stimulus types in a way that is stable across 15 days when defined with the first 3 temporal components compared to firing rates. All lines show the mean ± s.e.m. Linear mixed effect: time, tuning definition method as fixed effect, tuned single units from all mice as random effect. N = 699, 719, 665, 831 stimulus-tuned single units for DG,SG,RFG,NI respectively. See Supplementary Table 1 for additional reporting on sample size and statistics.

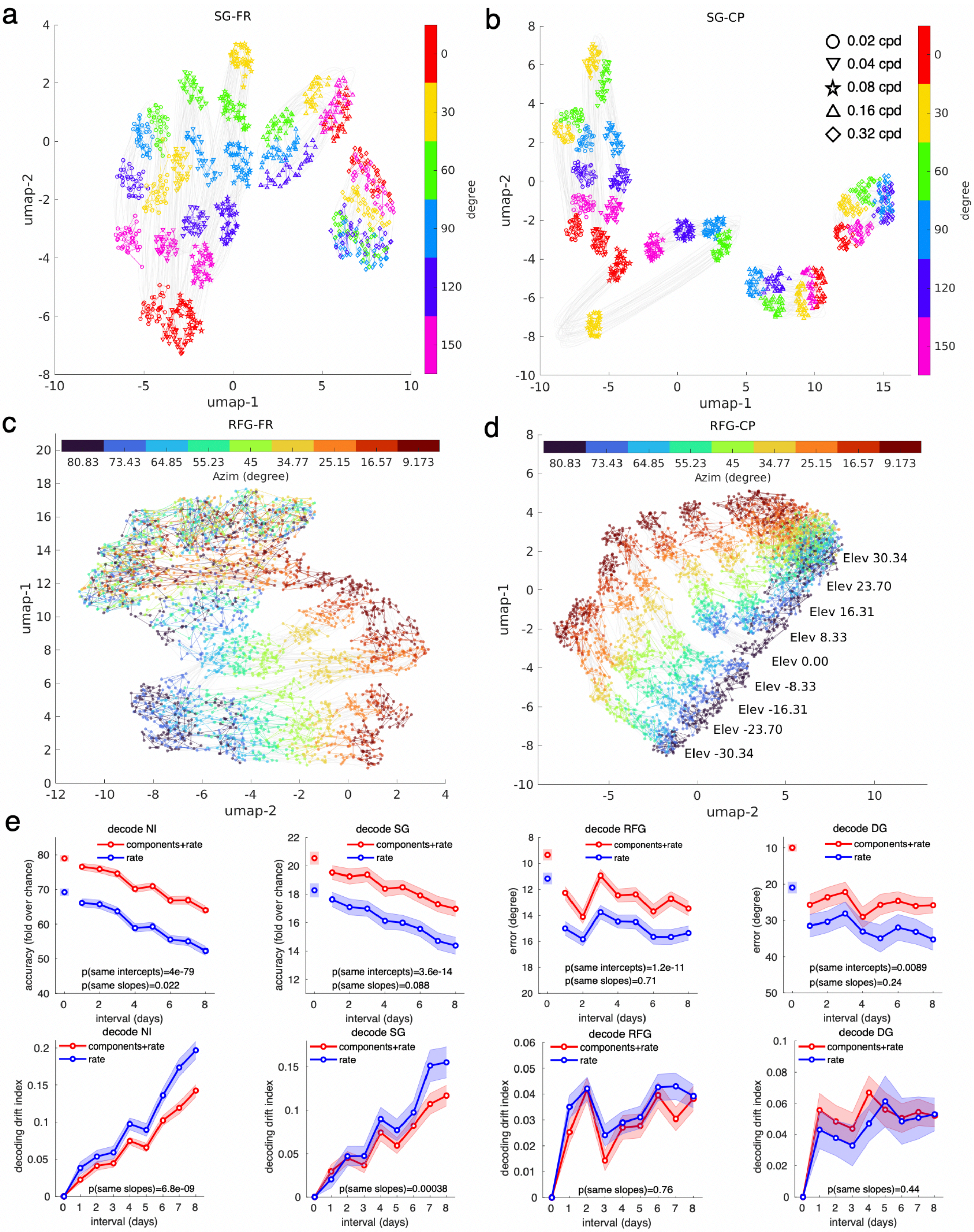

a
SG-FR
umap-1
umap-2
degree
0
30
60
90
120
150
b
SG-CP
0.02 cpd
0.04 cpd
0.08 cpd
0.16 cpd
0.32 cpd
umap-1
umap-2
degree
c
RFG-FR
80.83
73.43
64.85
55.23
45
34.77
25.15
16.57
9.173
Azim (degree)
umap-2
umap-1
d
RFG-CP
Azim (degree)
Elev 30.34
Elev 23.70
Elev 16.31
Elev 8.33
Elev 0.00
Elev -8.33
Elev -16.31
Elev -23.70
Elev -30.34
umap-2
umap-1
e
decode NI
components+rate
rate
accuracy (fold over chance)
p(same intercepts)=4e-79
p(same slopes)=0.022
interval (days)
decode SG
accuracy (fold over chance)
p(same intercepts)=3.6e-14
p(same slopes)=0.088
decode RFG
error (degree)
p(same intercepts)=1.2e-11
p(same slopes)=0.71
decode DG
error (degree)
p(same intercepts)=0.0089
p(same slopes)=0.24
decode NI
decoding drift index
p(same slopes)=6.8e-09
decode SG
p(same slopes)=0.00038
decode RFG
p(same slopes)=0.76
decode DG
p(same slopes)=0.44
interval (days)

**Supplementary Fig10 Temporal dynamics enhance long-term populational representation discriminability against competing stimuli during static gratings and receptive field Gabors stimuli, related to Fig. 5**

**a-d**. For both static gratings (a-b) and receptive field Gabor (c-d) patterns, different stimuli were more distinctly and stably represented in low-dimensional UMAP space when using temporal components (b, d) compared to populational firing rate (a, c). Most stimulus conditions formed distinct clusters of 30 points (15 days x even/odd trial averages, linked by colored lines). Clear spatial structures were observed with continuous geometry. For example, those of the same spatial frequencies (linked by gray lines) or same orientations (colors) were represented in similar locations during static grating stimuli. Similarly, those of nearby azimuths (colors) or elevations (linked by gray lines) were represented in similar locations during receptive field Gabors stimuli. For all panels, the original high-dimensional representation was constructed with all tuned neurons that appeared for 15 days in one example mouse. Either daily averaged odd/even trial populational firing rates (a, c) or daily averaged top three components (b, d) were used. **e**. Compared with those of firing rate alone, decoding performance of and natural images (column 1), static gratings (column 2), receptive field Gabors (column 3) and drifting gratings (column 4) improved significantly when the top 7 temporal components were incorporated (top row). Interval 0 indicates the accuracy of 1/7 held out trials in the first 7 days when the decoder was trained. Chance level equals to 1/100, 1/30, 35.4°, and 90° respectively). Decoding drift index exhibited smaller (NI and SG) or equal (DG and RFG) increase over time (bottom row). The plots are presented in mean ± s.e.m. Linear mixed effect: time, decoding strategy as fixed effect; Individual mouse-stimulus pair as random effect n = 500, 150, 125, 80 mouse pattern pairs for the 4 stimuli respectively, only 25 non-overlapping stimuli out of the 81 shown in RFG were used for decoding. See Supplementary Fig13c for the same decoding analysis but with all decoding quantification reported in terms of accuracy. See Supplementary Table 1 for additional reporting on sample size and statistics.

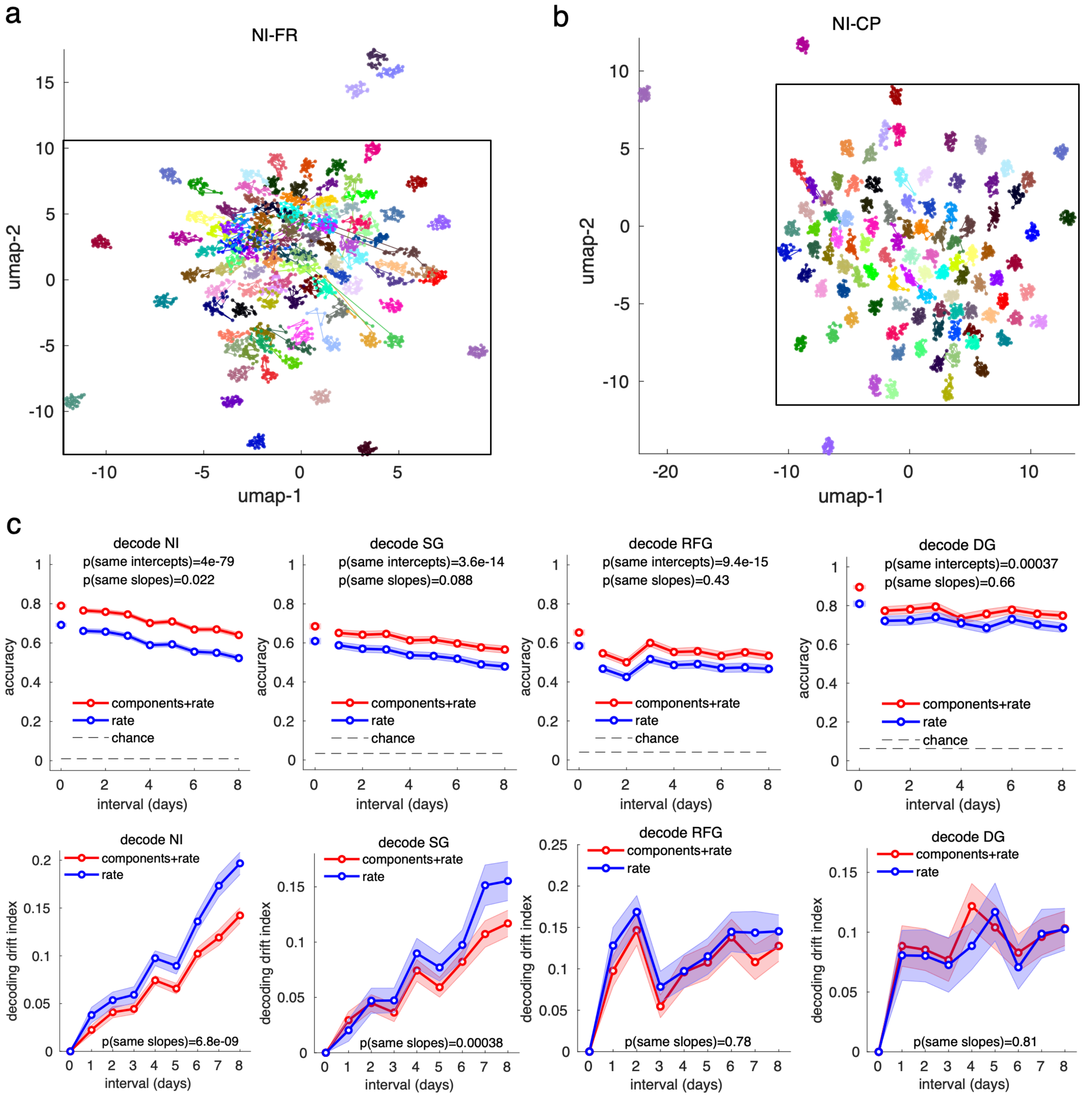


**Supplementary Fig11 Temporal information stably increases populational representation discriminability and visual decoding accuracy, Related to Fig. 4, Fig. 5**

**a-b.** Zoomed-out plot of Fig. 5e-f, black rectangle indicates the selected clusters corresponding to the 97 natural images in Fig. 5e-f. **c.** Same decoding analysis as in Supplementary Fig10 e, but with all decoding quantification reported in terms of accuracy.

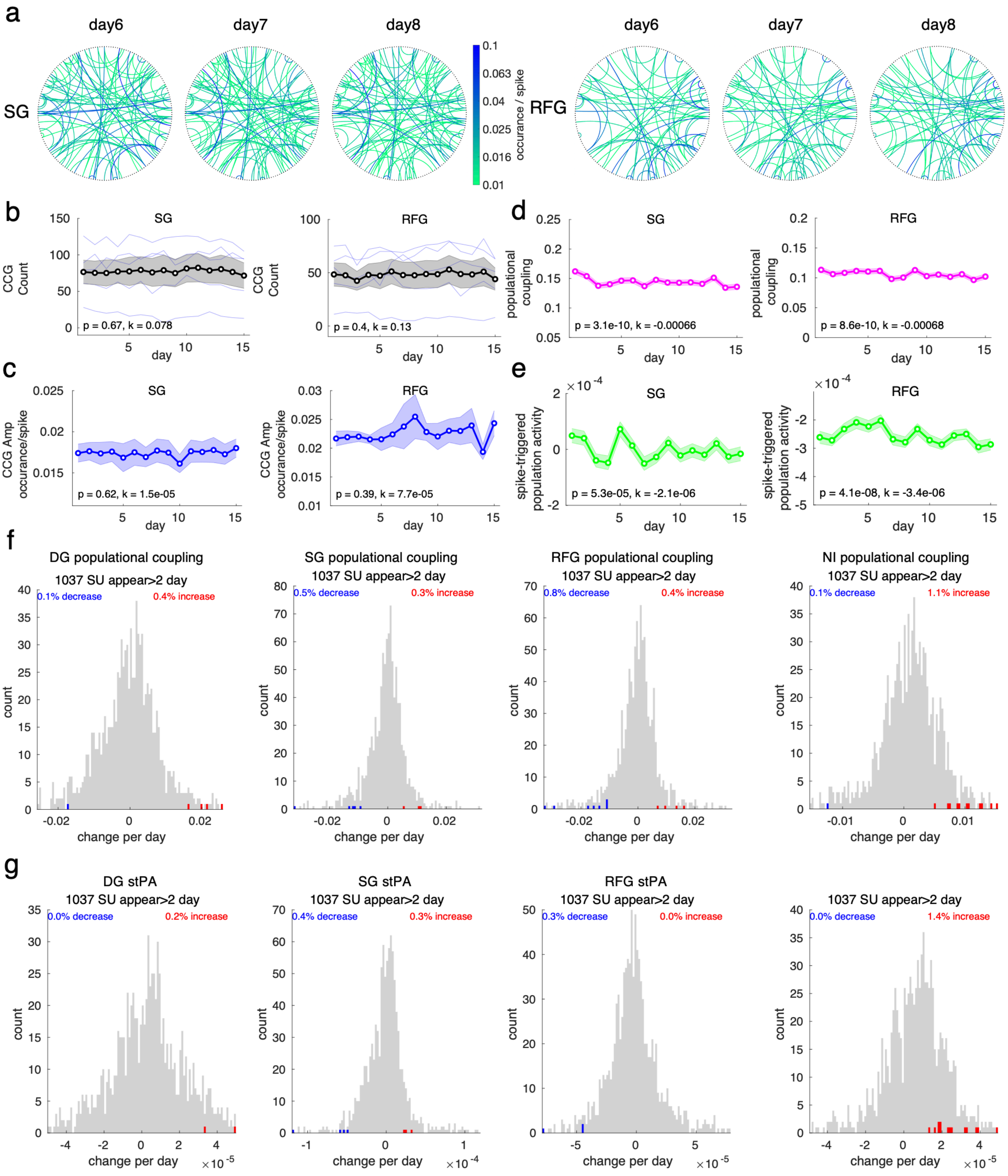


**Supplementary Fig12 Stability of functional neuronal network, related to Fig. 6**

**a**. All significant CCGs (curved lines, color-coded by CCG peak amplitude) between neurons (black dots aligned on the big outer circles) in one mouse during the two stimulus types (SG: static gratings and RFG: receptive field Gabors) showed similar connection patterns across days (columns), with neurons randomly placed on the circle but the order fixed across days and stimuli. Only neurons that had at least one significant connection across 15 days during either DG or NI stimuli were included (n=189). **b-c**. The CCG count (b) and strength (c) remained stable for the two stimulus types, showing the count of significant CCGs (b) and the mean CCG peak amplitude (c) per animal across 15 days. Linear mixed effect, time as fixed effect, individual animals (n=5) as random effect. individual significant CCGs (n=613,410 for SG and RFG, respectively) as random effect. Lines represent population mean ± s.e.m. **d-e**. Single neuron to population synchrony changed minimally across time for the two stimuli. The population synchrony score across time was measured either by population coupling (d) or spike-triggered population firing at 0 ms (e) or for all tuned single neurons that appeared for more than 2 days from all mice, showing the mean, 95% CI of all pairs across time. Linear mixed effect model (LME), time as fixed effect, individual neuron as random effect (n=1037 single units appearing more than 2 days). **f**. Few percentages of neurons significantly increased or decreased their population coupling for the four stimuli (columns, DG: drifting gratings; NI: natural images), showing a histogram of rate of change estimate for individual neurons of those significantly increased (red), decreased (blue), or non-significantly changed (gray). Linear regression, $P < 0.05$, Bonferroni corrected. **g**. Same as in f but for spike-triggered population firing. See Supplementary Table 1 for additional reporting on sample size and statistics.

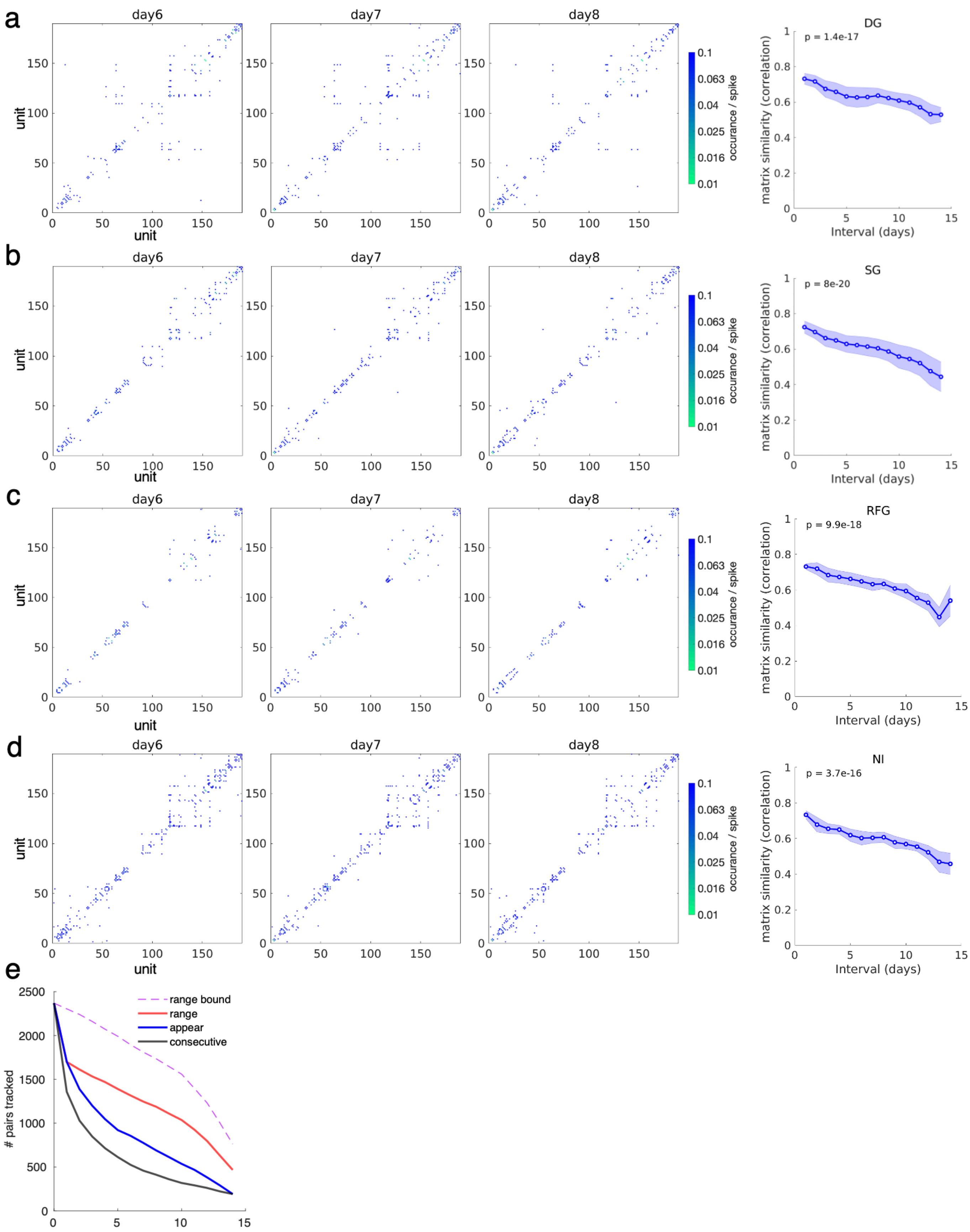


**Supplementary Fig13 Stability of functional neuronal network, related to Fig. 6**

**a-d.** Left: All significant CCGs between neurons in one mouse during the 4 stimuli (DG(a) , SG(b), RFG(c), NI(d)) showed similar connection patterns across days. Neurons were sorted by depth locations along each electrode shank and appended across all shanks in one animal. The neuron order was fixed across days. Only neurons that had at least 1 significant connection during either DG or NI stimuli across 15 days were included (n=189). Right: similarity (correlation coefficient) of connectivity matrices dropped gradually across days. Linear mixed effect, duration as fixed effect, individual animals (n=5) as random effect. Lines are in population mean ± s.e.m.
**e.** Quantification of trackable CCG pairs, like Fig. 6f. This plot shows the number of pairs tracked without calculating the tracked proportion. Tracked duration was determined based on three criteria: the number of sessions appeared, consecutive sessions appeared, and the range between the first and last seen session. The first and last co-appearing sessions for the two units forming a given pair of CCG determine the maximal possible trackable days for that pair (range bound).

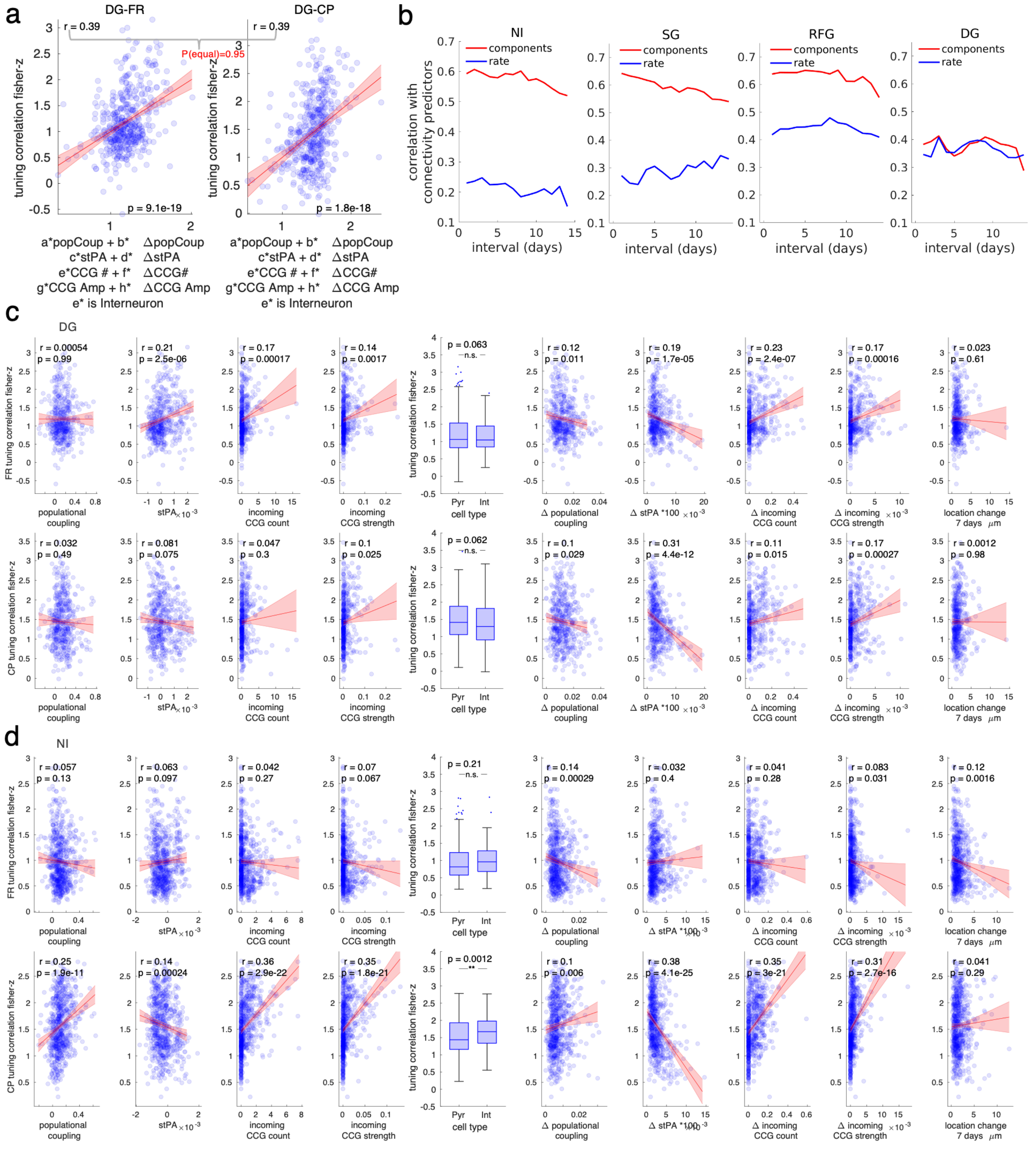


**Supplementary Fig14 Functional connectivity and single neuron-to-population synchrony better explain components-based tuning stability than that of firing rate-based during drifting grating and natural image stimuli; related to Fig. 6, Supplementary Fig12**

**a.** Stability of static grating stimuli after 7 days, defined either using firing rate (left) or first three components (right), was jointly explained by 1. population coupling, 2. spike-triggered population activity at 0 ms lag, 3.

Total incoming CCG counts, 4. total CCG strength, 5. the rate of change of variables 1-4 per day, 6. cell type. Correlation coefficients between tuning curves were Fisher Z-transformed. Red line: linear regression fitted response with 95% CI, text: correlation coefficient (square root of $R^2$ of linear regression), significance level for differences in correlation (Z-test). **b**. The plot shows correlation coefficient (square root of R2 of linear regression) between tuning curve similarity and assorted connectivity metrics over all time intervals. For 3 out of the 4 stimuli, correlation between network connectivity and tuning curve correlation is higher when based on temporal components compared to rate code. **c.** Tuning stability during drifting grating stimuli defined using firing rate (top row) and first three temporal components (bottom row) were separately explained by 1. population coupling, 2. spike-triggered population activity at 0 ms lag, 3. Total incoming CCG counts, 4. total CCG strengths, 5. the rate of change of variable 1-4 per day, 6. cell type 7. unit location changes 7 days apart. Red line: fitted linear regression response with 95% CI, text: correlation coefficient, significance level. N=481 stimulus-tuned single units appearing > 2 days and reappeared after 7 days (300 pyramidal neurons, 181 interneurons) **d.** As in c. but for natural images stimuli. N=684 neurons (451 pyramidal neurons, 233 interneurons). Box plots are in median, 25 to 75 percentiles. Whiskers represent 1.5-fold interquartile range below Q1 or above Q3. Outliers are indicated in scatters. See Supplementary Table 1 for additional reporting on sample size and statistics.

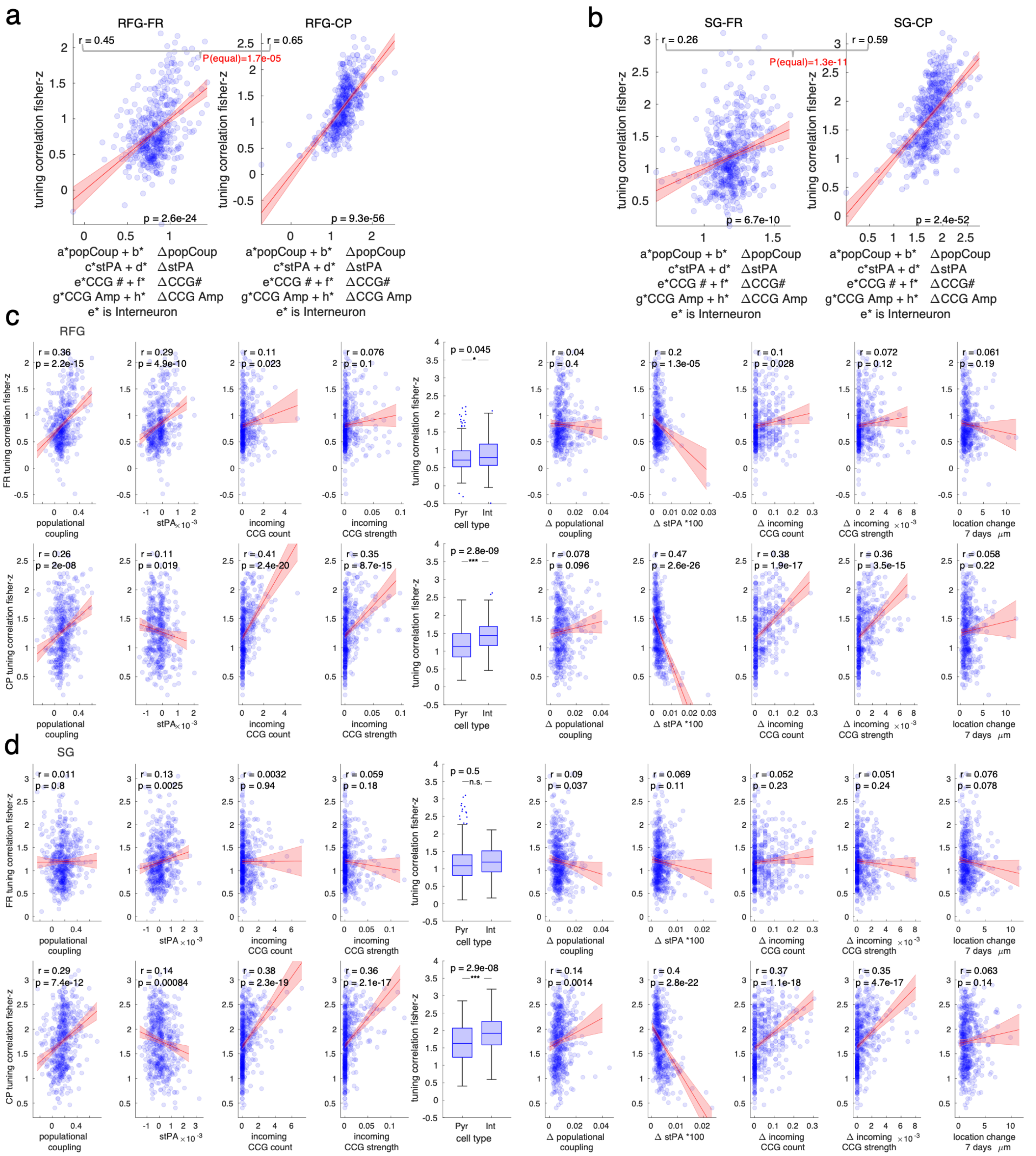

**Supplementary Fig15 Functional connectivity and single neuron-to-population synchrony better explained component-based tuning stability than that of firing rate-based during receptive field Gabors and static gratings stimuli; related to Fig. 6, Supplementary Fig12**

**a.** Stability of receptive field Gabors stimuli after 7 days defined using either the firing rate (left) or the first three components (right) was jointly explained by 1. population coupling, 2. spike-triggered population activity at 0 ms lag, 3. Total incoming CCG counts, 4. total CCG strength, 5. the rate of change of variable 1-4 per day, 6. cell type. Correlation coefficients between tuning curves were Fisher Z-transformed. Red line: linear regression fitted response with 95% CI, text: correlation coefficient (square root of $R^2$ of linear regression), significance level for differences in correlation (Z-test). **b.** As in a. but for static gratings stimuli. **c.** Tuning stability during receptive field Gabors stimuli defined using firing rate (top row) and first three temporal components (bottom row) were separately explained by 1. population coupling, 2. spike-triggered population activity at 0 ms lag, 3. Total incoming CCG counts, 4. total CCG strengths, 5. the rate of change of variable 1-4 per day, 6. cell type 7. unit location changes 7 days apart. Red line: fitted linear regression response with 95% CI, text: correlation coefficient, significance level. N=455 stimulus-tuned single units appearing > 2 days and reappeared after 7 days (277 pyramidal neurons, 178 interneurons). **d.** As in c. but for static gratings stimuli. N=533 neurons (338 pyramidal neurons, 195 interneurons). Box plots are in median, 25 to 75 percentiles. Whiskers represent 1.5-fold interquartile range below Q1 or above Q3. Outliers are indicated in scatters. See Supplementary Table 1 for additional reporting on sample size and statistics.

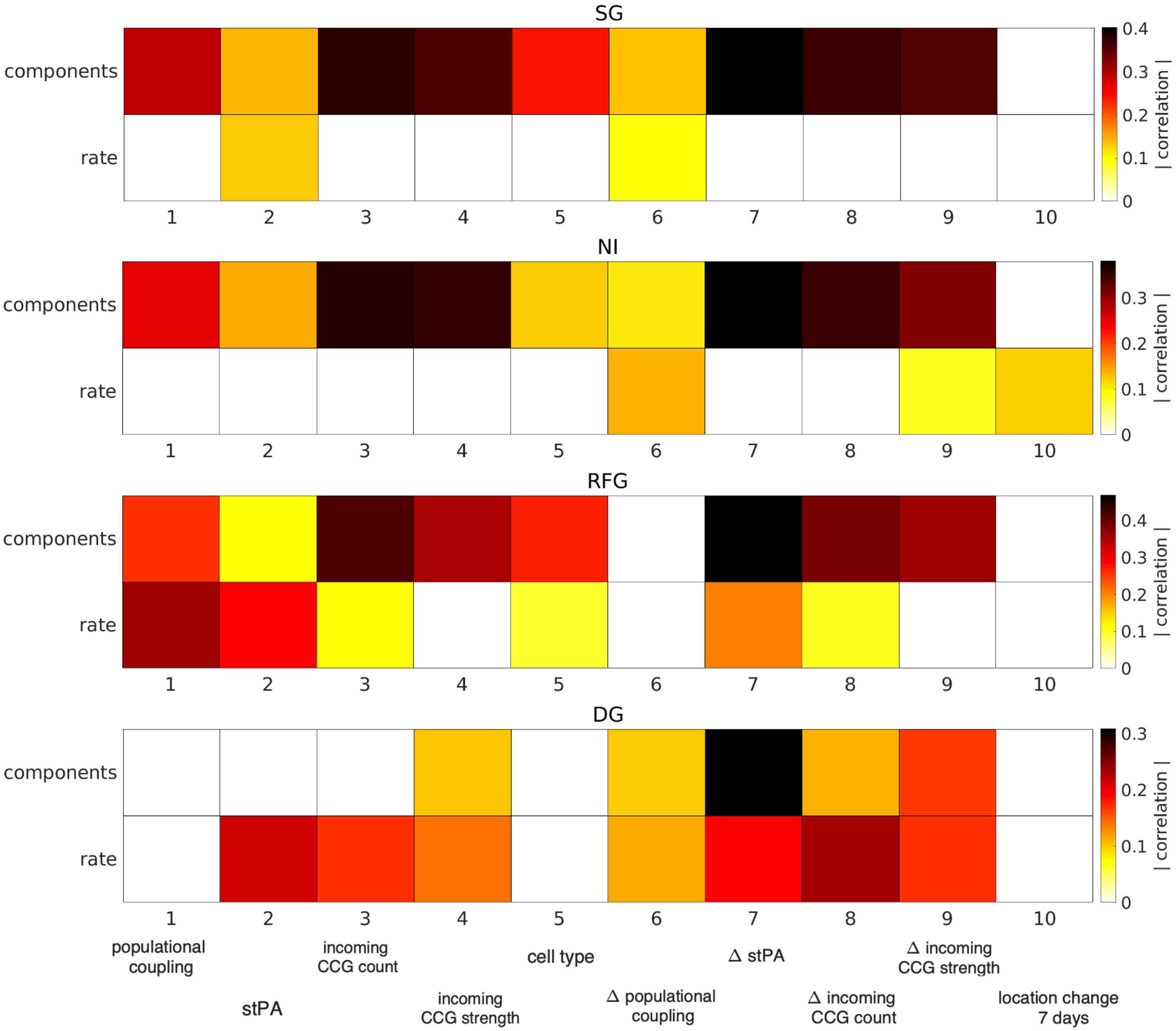


**Supplementary Fig16 A summary of the correlation strength between tuning curve similarity after 7 days and different variables tested in Supplementary Fig14 and Supplementary Fig15**

Insignificant correlations were set to 0. Temporal components based tuning curve similarity were better associated with many connectivity metrics than firing rate based for 3 out of 4 types of stimuli tested (visual inspection, statistics not performed).

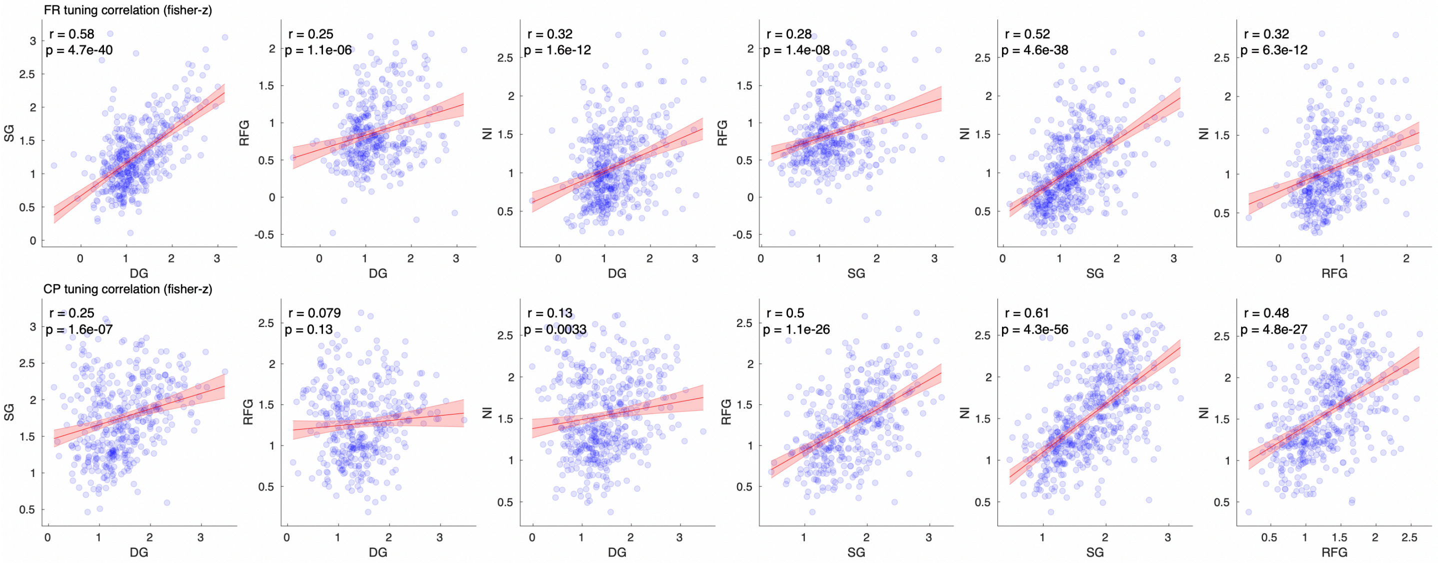


**Supplementary Fig17 Predicting the stability of one stimulus type with stability of other stimulus types**
Tuning curve similarity after 7 days for one stimulus type is correlated with tuning curve similarity after 7 days for other stimulus types, showing all combinations of stimulus type pairs (4 choose 2) for firing rate-based similarity (top row) and temporal components-based similarity (bottom row). Red line: linear regression fit with 95% CI, text: correlation coefficient, significance level. Across all pairs of comparison, the stability seems positive correlated between different stimulus pairs, with particularly strong prediction power between static grating versus natural image stimuli, which may be due to the similarity in the visual pattern themselves (both static and full field). N = 429, 371, 476, 398, 529, 446, 429, 371, 476, 398, 529, 446 (left to right, top to down) single and tuned (tuned to both stimuli) units appearing > 2 days and reappeared after 7 days. See Supplementary Table 1 for additional reporting on sample size and statistics

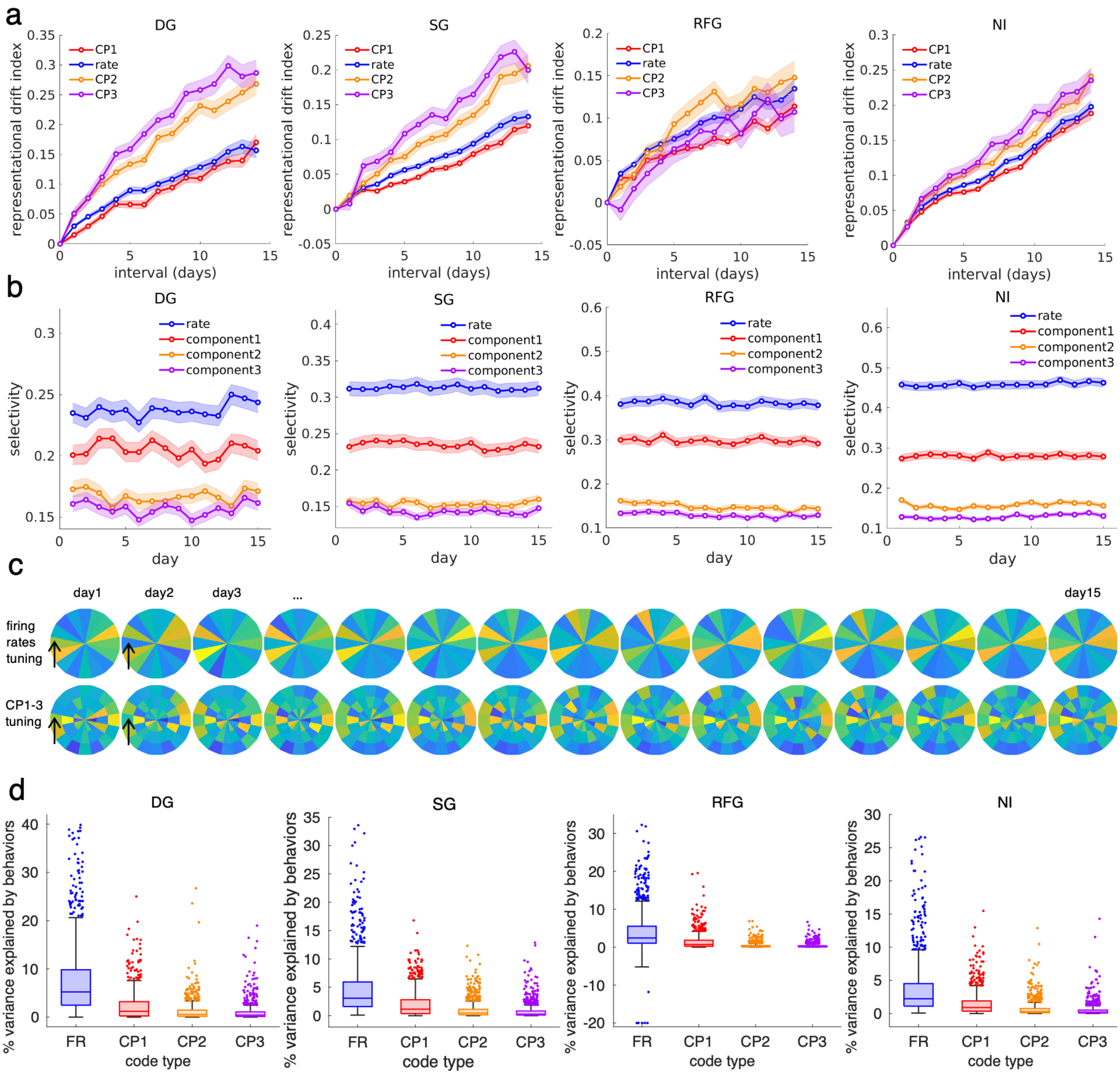


**Supplementary Fig18 Comparison of tuning and selectivity stability of firing rate-based tuning curve versus individual component-based, more explanations on the functionality of components, related to Fig. 4**

**a.** Comparison of representational drift index computed using different features (firing rate versus individual temporal components). **b.** temporal components were less stimulus-selective over time than firing rate for all four stimulus types (DG: drifting gratings; SG: static gratings; RFG: receptive field Gabors; NI: natural images). A selectivity of 1 means the neuron is modulated by only one stimulus, while a selectivity of 0 means it responds equally to all stimuli. All lines show mean + s.e.m. **c.** The example (Fig. 4a-c) neuron's firing rate-based tuning curve (top) versus top 3 components-based tuning curve (bottom) across 15 days. The component one is on the outermost ring. See Fig. 1 for color-scale. **d.** Firing rates code are better explained by non-stimulus variable. The percentage variance in different neural code (firing rates, temporal components 1-3) was jointly explained by behavioral non-stimulus variables including but not limited to pupil size, animal motion(limb speed or treadmill rotations)[7], LFP estimated EMG[8], LFP average envelopes magnitude[9] from multiple bands (e.g., 3-7Hz[10]), phase of delta band[11], trial number within-day[12]. GLM was fitted for firing rates (Poisson kernel) or components (Gaussian kernel) following the guidance in[13]. (Values below -20% were set to -20% for ease of visualization, 5 instances adjusted for FR during RFG) Box plots are in median, 25 to 75 percentiles. Whiskers represents 1.5-fold interquartile range below Q1 or above Q3. Outliers are indicated in scatters.

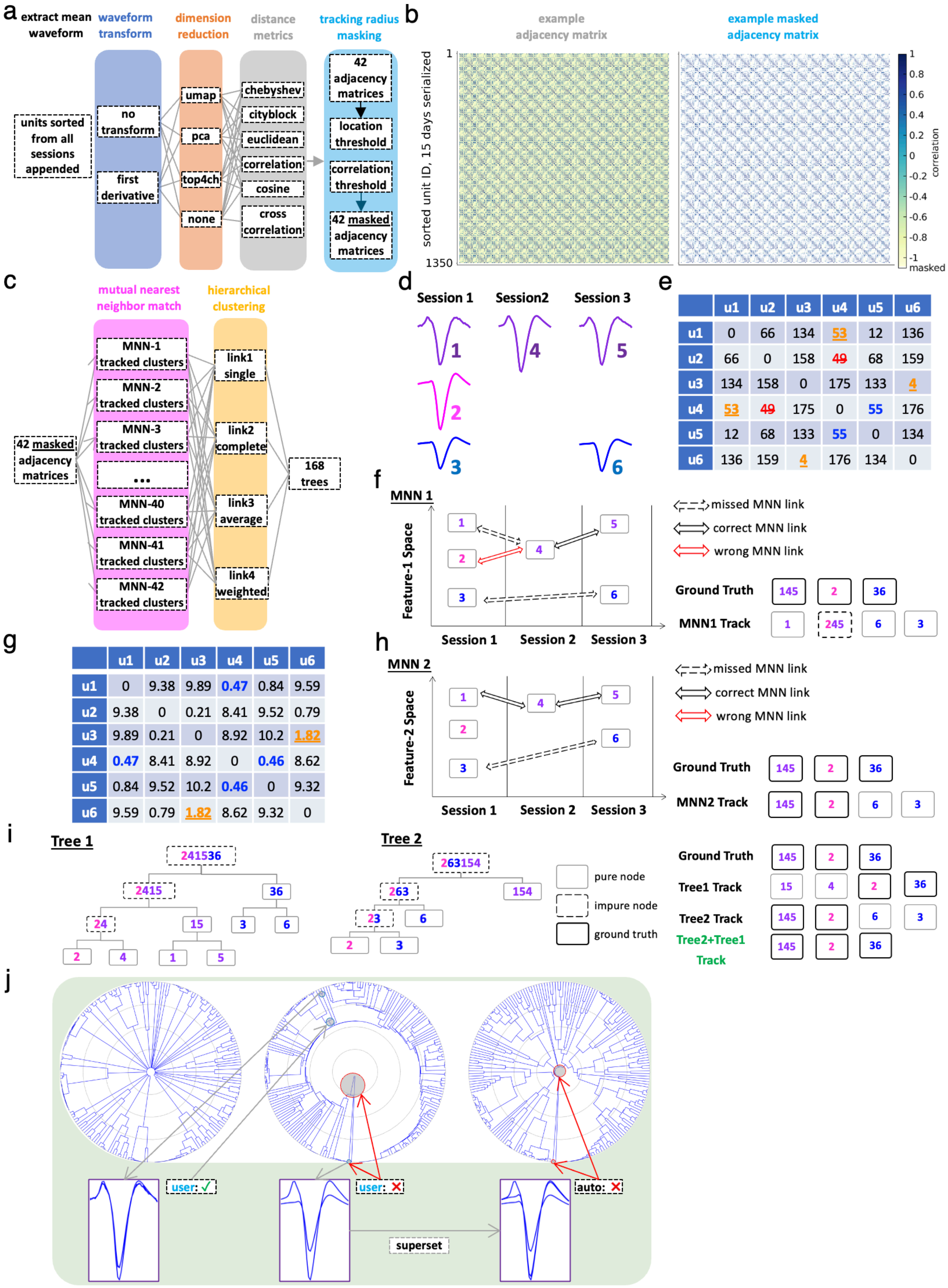
a
extract mean waveform
waveform transform
dimension reduction
distance metrics
tracking radius masking
units sorted from all sessions appended
no transform
first derivative
umap
pca
top4ch
none
chebyshev
cityblock
euclidean
correlation
cosine
cross correlation
42 adjacency matrices
location threshold
correlation threshold
42 masked adjacency matrices
b
example adjacency matrix
example masked adjacency matrix
sorted unit ID, 15 days serialized
1
1350
correlation
masked
c
mutual nearest neighbor match
hierarchical clustering
42 masked adjacency matrices
MNN-1 tracked clusters
MNN-2 tracked clusters
MNN-3 tracked clusters
...
MNN-40 tracked clusters
MNN-41 tracked clusters
MNN-42 tracked clusters
link1 single
link2 complete
link3 average
link4 weighted
168 trees
d
Session 1
Session2
Session 3
e
u1 u2 u3 u4 u5 u6
u1 0 66 134 53 12 136
u2 66 0 158 49 68 159
u3 134 158 0 175 133 4
u4 53 49 175 0 55 176
u5 12 68 133 55 0 134
u6 136 159 4 176 134 0
f
MNN 1
Feature-1 Space
Session 1
Session 2
Session 3
missed MNN link
correct MNN link
wrong MNN link
Ground Truth
MNN1 Track
g
u1 u2 u3 u4 u5 u6
u1 0 9.38 9.89 0.47 0.84 9.59
u2 9.38 0 0.21 8.41 9.52 0.79
u3 9.89 0.21 0 8.92 10.2 1.82
u4 0.47 8.41 8.92 0 0.46 8.62
u5 0.84 9.52 10.2 0.46 0 9.32
u6 9.59 0.79 1.82 8.62 9.32 0
h
MNN 2
Feature-2 Space
Session 1
Session 2
Session 3
missed MNN link
correct MNN link
wrong MNN link
Ground Truth
MNN2 Track
i
Tree 1
Tree 2
pure node
impure node
ground truth
Ground Truth
Tree1 Track
Tree2 Track
Tree2+Tree1 Track
j
user: ✓
user: ✗
auto: ✗
superset

**Supplementary Fig19 Principles of the semi-automatic method for tracking the same neurons across sessions**

**a.** A flow chart for the initial tracking process. An ensemble of diverse adjacency matrices was created. **b.** Example raw adjacency matrix formed by units from one probe pooled from 15 days using correlation as the distance metric. Units were sorted by session. Within each session, units were sorted by depth locations along the shank. (14*2)+1=29 stripes can be seen, representing potentially repeatedly identifiable units that were similar to themselves between sessions. A large number of elements in each adjacency matrix were masked after applying simple location and correlation thresholds. **c.** A flow chart for later tracking procedures. The pairwise adjacency matrices were initially flattened (forming clusters) using the mutual nearest neighbor algorithm between all adjacent sessions. The initial clusters were further linked recursively to form 168 hierarchical clustering trees. See (d) to (j) for expanded explanation of these processes. **d.** A simulated scenario demonstrating the tracking procedure: six hypothetical units sharing the same top channel were detected/sorted in 3 individual sessions. Unit group 1-4-5 represented the same unit across time, while unit group 3-6 were another neuron which did not appear in the second session. Unit 2 was a different unit from groups 1-4-5 and 3-6, appearing only in session 1. For simplicity, only the top channel waveform of each unit is shown; in actual tracking, all available channels from a 32-channel shank were used. Unit 2 was obtained from a real recording, while unit group 1-4-5 (characterized by a positive peak before depolarization with slight amplitude fluctuation across sessions) and unit group 3-6 (major amplitude change from unit 2 with slightly different depolarization and repolarization slopes) were derived from Unit 2 to simulate a difficult tracking scenario. **e.** Distance matrix comparing the mean squared error (distance feature 1) between all pairs of units. This serves as the starting point of the past unsupervised method and the initialization portion of the current method. Mutual nearest neighbor (MNN) pairing across adjacent sessions was performed on this matrix. Red strikethrough: False positive pairs detected by the MNN. Orange: Missed pairs by the MNN. Blue: Correctly detected MNN pairs. **f.** The distance matrix in (e) was transformed into a one-dimensional feature space (e.g., using a multidimensional scaling algorithm) and plotted across time to illustrate the MNN matching result. Unit 2 incorrectly prevented Unit 1 from linking with Unit 4, as Unit 2 was closer to Unit 4 than Unit 1. Similarly, Unit 3 failed to link with Unit 6 due to the absence of a mutual nearest neighbor in session 2, preventing a bridge between Unit 3 (session 1) and Unit 6 (session 3). Consequently, MNN tracking based on this feature resulted in three fragmented units and one incorrectly tracked unit group. **g.** Another distance matrix comparing correlation distance (distance feature 2) between all pairs of units. For ease of presentation, the data was transformed into (1-correlation coefficient)*1000. Mutual nearest neighbor (MNN) pairing across adjacent sessions was performed on this matrix. Orange: Missed pairs by the MNN. Blue: Correctly detected MNN pairs. **h.** Same as (f) but for summarizing the MNN result in (g). Using this distance feature resulted in different tracking outcomes (two correctly tracked units and one fragmented unit) but still failed to reach the ground truth. **i.** Hierarchical clustering trees ignore session labels and attempt to form multiple possible links, ultimately joining all units. This approach solves the issue of missing session. (e.g., Tree 1, where unit3 is successfully linked with unit6). Leaf nodes in Trees 1 and 2 were initialized based on all valid MNN pairs found in (f) and (h) respectively. Invalid MNN pairs (identified after visual inspection) in (f) and (h) were broken into individual elements before tree formation. As trees progress from leaf nodes (bottom) to root nodes (top), they become increasingly impure (containing incorrect merges). However, since users manually verify tree nodes, tracking terminates before reaching impure root nodes, ensuring that no false linkages are automatically generated. Nevertheless, hierarchical clustering trees, like MNN, face limitations as no single distance metric fits all cases. Tree 1 failed to correctly track Unit 1-4-5, while Tree 2 failed for Unit 3-6. Green: Combining results from both Tree 1 and Tree 2 produced the ground truth track. This justifies the approach outlined in (a) and (c), where multiple trees were formed using different combinations of distance metrics and hierarchical clustering methods. **j.** The procedure for cutting multiple trees at appropriate heights. Units in (b) were used to create Hierarchical clustering trees (only 3 shown for simplicity; 168 trees were used in total). The trees were visualized in circular layouts with root nodes at the center. Waveforms of the tracked units were overlaid and presented to users for confirmation. (Insets: top channel waveforms of units contained in tree nodes; actual tracking used all electrode channels.) User confirmation of unit merging results in selecting the parent node as the next query target. Conversely, rejecting a merge leads to automatically cutting all parent nodes in that tree. Nodes from other trees containing a superset of the rejected node's units are also automatically cut. This process continues until all nodes across all trees are verified, ultimately yielding a final set of correctly tracked units.

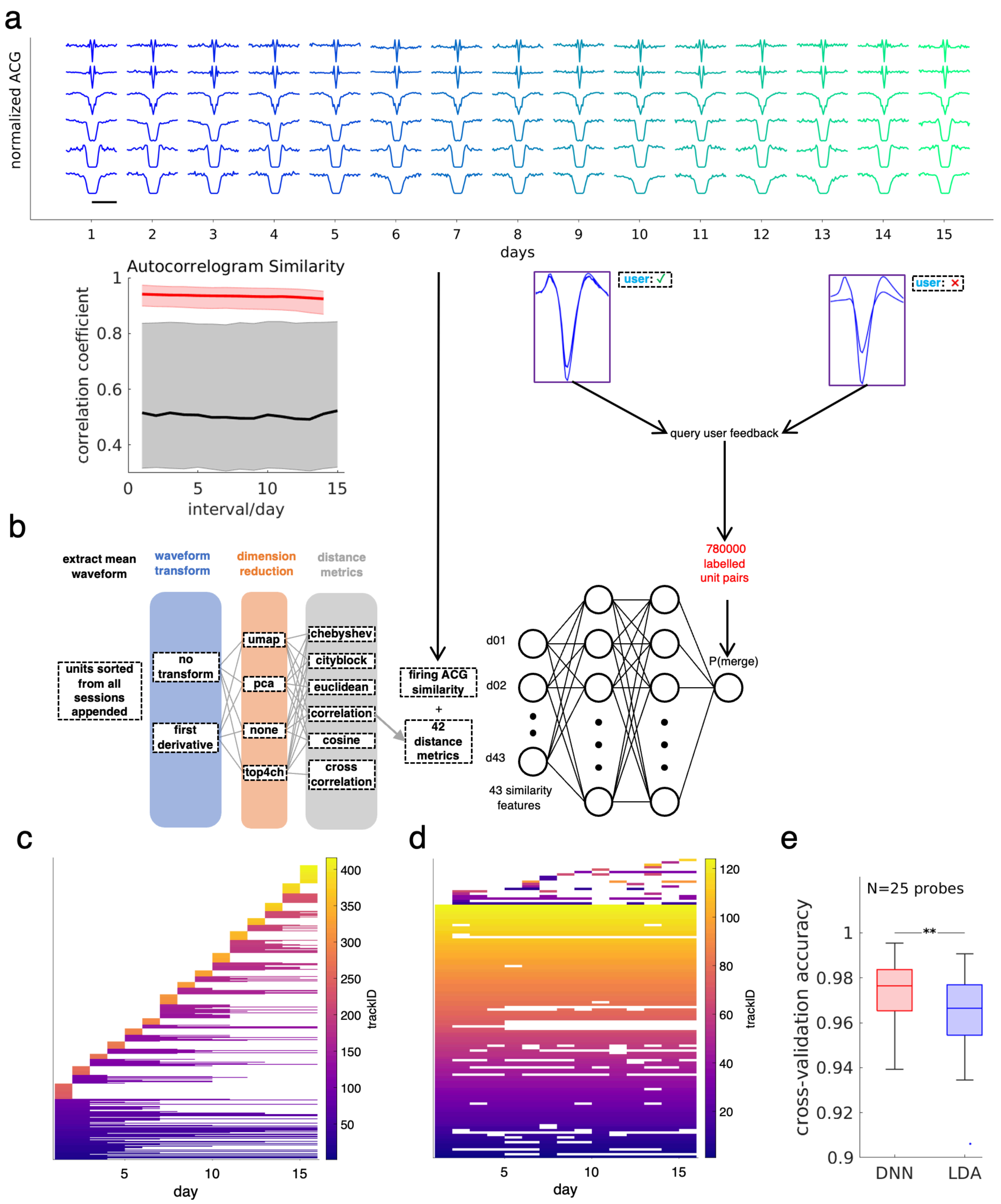


**Supplementary Fig20 Validation of tracking the same neurons, related to Fig. 1 and Supplementary Fig1**

**a.** Validation of using unit autocorrelogram (ACG) as an additional feature for tracking the same neurons. Top: Example ACGs from 6 units (rows) across 15 days (columns). ACGs maintained nearly identical shapes over time. Scale bar: 50 ms. Bottom: Single-unit ACGs were more like themselves (red) across sessions than to other units from the same probe within the same session (black, all valid single units across all mice. $P<0.001$ see Supplementary Table 1). **b.** Flowchart of the tracking validation process. Diverse pair-wise distance metrics (see Supplemental Methods and Supplementary Fig19) were computed for all sorted units pooled from individual days in a probe. 780,000 user-labeled unit pairs (Supplemental Methods and Supplementary Fig19) were collected across 25 probes as ground truth. Each pair contained 43 distance features and a binary user label of "same" or "different" units. A dense neural network was trained to reproduce user decisions outputting the probability of merging any two units. **c-d.** Comparison of tracking performance between Mutual nearest neighbors-based tracking[1] (MNN, Supplemental Methods and Supplementary Fig19, Fig. 1d) and the current method. MNN method (c) was applied to units in an example probe. The horizontal lines (tracks) corresponded to the same units. Units could be tracked for limited duration. The resulting unit longevity tally plot fragmented into multiple short pieces. **d.** Unit longevity tally for the method: The number of fragmented tracks was drastically reduced, and average longevity increased compared to (c). **e.** Percentage of manual decisions of a given probe that were reproduced by either the deep learning method or a simple linear discriminant analysis method trained with labels from the other 24 probes. High reproducibility indicates strong internal labeling consistency. Paired t-test $p < 0.01$. Box plots are in median, 25 to 75 percentiles. Whiskers represents 1.5-fold interquartile range below Q1 or above Q3. Outliers are indicated in scatters. See Supplementary Table 1 for additional reporting on sample size and statistics.

### Supplemental Methods

#### Tracking

The mean waveform for all available channels in a probe was extracted from -0.8ms to 1.12ms of the spike-sorted timestamp for each sorted unit. It was used for the calculation of assorted pairwise unit distance metrics (Supplementary Fig19a). Units from all sessions were directly appended and treated as coming from one session. The goal of the tracking was to form a manual ground truth adjacency matrix, which means for each pixel (each unit pair) in the matrix, label whether the pair can be merged (same units) or should be kept separated (different units). After which, the labeled adjacency matrix was converted to flat clustering (clusters/tracks of repeatedly identified same units) by finding connected components. We opted for the seemingly "painstaking"[14] visual inspection supervision to ensure the best possible reliability of tracked units. Consistent with past literature, such final manual curation steps for cluster merging after initial tracking still play the last line of correction despite numerous steps of automation[15, 16]. To reduce the amount of manual effort, three strategies were applied: 1. Distance thresholding (masking) of invalid pixels in the adjacency matrix (Supplementary Fig19). 2.Unsupervised Mutual nearest neighbor[1] match (MNN) between units from temporally adjacent sessions for track initialization (Supplementary Fig19). 3. Active learning (user feedback) with an ensemble of hierarchical clustering trees whose leaf nodes were initialized by MNN (Supplementary Fig19). Finally, we checked manual labeling consistency with cross-validated deep learning and linear discriminant analysis methods (Supplementary Fig20).

Here we further explain the 3 core principles behind the tracking.
1. Distance thresholding: a large number of the $\binom{n}{2}$ pairwise decisions in the adjacency matrix did not merit further visual inspection. They were automatically masked as 0 following distance thresholds, such as having correlation coefficient between unit waveforms smaller than 0.7. Supplementary Fig19b
2. Mutual nearest neighbor match: Unit X in session N is matched/linked to another unit Y in the next immediate session N+1 if a) Y is the closest unit in session N+1 relative to unit X and b) unit X is the closest unit in session N relative to unit Y. The process was repeated until all possible links/tracks of units were formed. An example matching process for 6 units across 3 sessions using 2 different distance metrics were shown in Supplementary Fig19.d-h . Despite the algorithmic simplicity, MNN could not link units that show up on non-consecutive days (Supplementary Fig19f,h). In addition, false positive matches could still be generated (Supplementary Fig19f).
3. For each MNN matched unit tracks, a hierarchical clustering tree initialized by those tracks ignored session labels and attempted to form increasingly larger clusters/tracks that eventually merged all units across all sessions. (Supplementary Fig19i). This solved the issue of MNN which fails to link units showing up on non-consecutive sessions. Users were prompted to stop the merging at appropriate heights of each tree branches. Finally, combining tree cutting results from multiple trees improved the final tracking performance compared to only using individual trees (Supplementary Fig19i). To improve reviewing efficiency, the next node to be prompted to the user was intelligently selected (actively learned) given the user's feedback on the current node. (Supplementary Fig19j).

Distance thresholding was performed with two types of features: fixed-threshold distance features and adaptive-threshold distance features. Three fixed-threshold distance features were used across all iterations of the tracking process, whereas adaptive-threshold features were only used to mask distance matrices in the preliminary MNN round.

The fixed-threshold distance features included Weighted Center distance d(WC), top 4 channels cross-correlation (Top4Xcorr), Cross-Peak-Percentage-Difference-Sum (XPPS). d(WC) features were computed as the Euclidean distance between the spatial weighted centroid[15] of the two units. When computing the weighted centroid location, P2P amplitude of each channel was raised to the power N. N=2 gives d(WC2) distance feature, and N=10 corresponds to d(WC10) feature. Since there was usually substantial decay in unit amplitude in space, d(WC10) can be thought of as a good approximation for top channel distance of the two units. During the tracking process, the spatial unit of WC2 and WC10 was specified in multiples of the shortest inter-electrode distance (pitch) instead of mm to accommodate electrode designs with slightly different dimensions. 1-Top4Xcorr was the per-channel averaged max waveform cross-correlation at each of the top 4 channels between the two units subtracted by 1 to convert correlation to distance. When the top 4 channels of the two units are not the same, the top 4 channels are expanded to the top N channels, where N contains a set of channels that is the union of the top 4 channels between the two. The XPPS feature was calculated for each pair unit by summing their percentage difference in top N channel P2P amplitude (N=1 by default). When the top channel of the two units differed, the union of top channels was selected. Then the percentage difference in peak amplitude for each top channel was computed as the difference in their amplitude divided by the larger of the two amplitude values of that channel before summing across all top channels. When the two units share a common top channel, XPPS is multiplied by 2 so it has the same scale as pairs whose top channels differ.

MNN initialization was performed twice. In the preliminary round, the goal was to use MNN to find a reasonable thresholding value for the 5-adaptive-threshold distance features. The 5 features were d(P2P), the absolute difference in P2P amplitude of the mean waveform; d(FWHM), the absolute difference in FWHM of the top channel of the two units; Waveform correlation distance d(correlation) is 1 – correlation, where correlation was Pearson's correlation coefficient of all channels serialized waveform; XPPS and top3channel XPPS.

In this round, 42 pairwise distance matrices were computed. They were masked by fixed-threshold features at d(WC2) <4.5 AND d(WC10) <4.5 AND (d(WC2) <2.25 OR d(WC10) <2.25) AND Top4Xcorr<0.3, Mutual nearest neighbor match was performed for each of the 42 masked distance matrices individually, forming 42 groups of tracks. For each group of tracks, 5% (empirical) of tracks were hypothesized to contain false positive members in the track. Therefore, for each of the tracks within the 5%, the minimum pairwise distance of the 5 adaptive-thresholds features was computed to define the maximally allowed thresholding value. These values were then averaged across 42 realizations to define commonly acceptable adaptive threshold values for all distance metrics.

In the first round of tree building, 42 pairwise distance matrices were masked by both the same fixed-threshold features as the MNN first round and the 5 adaptive-threshold features. In addition, if two units were from nearby sessions, stricter masks were applied (d(WC2) <1.5 OR d(WC10) <1.5), Top4Xcorr<0.2. MNN was recomputed. Within each MNN, the tracked units served as leaf nodes for hierarchical clustering with 4 types of linkage function, resulting in 42x4=168 hierarchical clustering trees. The agglomerative trees attempted to gradually merge all tracks together in the final root node. Then, the goal was to simply decide a set of stopping nodes between leaf nodes and the root node by first rejecting close-to-root nodes that violate distance masking criteria while repeatedly asking for user feedback on the rest of uncertain nodes, similar to the wizard function[17] for efficient sorting curation in a single recording session as previously reported. The detailed algorithm and ground truth dataset validation are described previously[18]. Briefly, this method allowed efficient labeling in all 168 trees simultaneously by sharing user feedback. On one hand, whenever the user approved any node, the next query target was searched across all trees to identify nodes whose members were superset of that of the current node. Such candidate nodes with the least number of members were prioritized for the next query to ensure judicious lengthening of the tracks. On the other hand, when a user rejected any node, those nodes across all 168 trees being the superset of the current one were automatically rejected simultaneously. Our ensemble method reduces to the efficient single tree binary search method[19] in situations where the user approves a node, yet the non-rejected superset nodes (next query candidates) can only be found in one tree.

During the labeling process, a series of overlaid plots of waveforms from a node (same units identified by the program) were presented to the user for reviewing. The user could either approve or reject this tracking. When decisions were hard to make, more information could be requested, in which case an interactive movie showing the tracked units' waveform in temporal order would be played for the user to view or scroll through. Autocorrelogram[20] (Supplementary Fig20a) in both the 50ms and 500ms window of the tracked units were plotted with the mean waveform as well. Additionally, CCG[20] (cross-correlogram) could be requested when units from the same session were included in the track for checking potential within-session merging. When a track contained all but a few units that the user approved, the user was allowed to enter the specific unit IDs to leave those units out while approving the rest. Another type of user feedback was delaying the decision to the next iteration in which a putative rejection was placed for the current decision to advance overall labeling progress while this decision was not respected when initializing next rounds.

In the second round of tree building, a stack of masks (d(WC2) <4.5) AND (d(WC10) <4.5) AND (d(WC2) <0.75 OR d(WC10) <0.75) AND (Top4Xcorr<0.1) AND (XPPS<0.6) was used. Moreover, user-rejected elements from the previous round were masked. The same 168 trees were created with the new masking, and only nodes that cannot be automatically labeled by past user decisions were sent for user review (e.g., those that the user labeled uncertain or newly proposed tracks). After this round, users were prompted to first additionally review all tracks that contained only one element (a unit that only showed up only once in the 15 sessions, putative noise) to decide if they should be permanently rejected as noise and prevented from merging with all other units. Consistent with past work, such manual noise rejection was required even if multiple quality metrics were applied to screen noise[5]. Secondly, for all remaining tracks, a random member was selected as track representative, its waveform was separately overlaid with 6 other representatives that were spatially close. The user was to decide which of the 6 were so different and can be confidently rejected even by checking this static representative with no temporal information. Neither those pairs of representatives nor the rest of the members in the tracks they represent were allowed to merge in the future rounds.

In the third or later round of tree building, the stack of mask applied was (d(WC2) <4.5) AND (d(WC10) <4.5) AND (d(WC2) <2.25 OR d(WC10) <2.25) AND (Top4Xcorr<0.3), nearby session unit pairs (d(WC2) <1.5) OR (d(WC10) <1.5) AND (Top4Xcorr<0.2). Moreover, user-rejected elements from the previous round were masked, noise units were masked, and the same 168 trees were created with the new masking. Only nodes that cannot be automatically labeled by past user decisions were sent for user review.

A final manual curation step would attempt to correct any obvious fixed-threshold masking-induced false negative (fragmented) tracks. For each track, the overlaid plot of all members was exported, and the plots were sorted by depth location while users scroll through different tracks to see if there were spatially adjacent tracks that were highly similar in waveform, showing up in complementary time-segments but not merged. The average time for answering each question was approximately 30 seconds, resulting in one month for fully reviewing tracks from all 25 probes.

**Explanation of tracking duration and the tracking probability analysis in Fig. 1e and Fig. 6f-g**
For example, if a neuron missed day1, day6 and day15 but appeared in all other days, then:
a) The range of this neuron was 13 days since the neuron's first and last appearance was on day2 and day14 respectively: 14-2+1=13.
b) The range bound of this neuron was 14 days, since the neuron first appeared on day2, the maximal possible tracking range this neuron was 15-2+1=14 days.
c) The appearance/attendance of this neuron was 12 days, since the neuron missed a total of 3 days: 15-3=12.
d) The (maximal) consecutive (appeared) days of this neuron was 8 days (from day7 to day14), while shorter segment of consecutive appearance (e.g., 4 days, from day2 to day5) were disregarded.
The four colored lines in Fig. 1e and Supplementary Fig13e represented a cumulative histogram of the number of the neurons that at least satisfy any given longevity criterion. Finally, Fig. 6f presented the probability of tracking a neuron for given amount of duration. Conceptually, it's the day-wise division of the other 3 curves against the range bound curve. Formally, it is calculated by dividing the number of neurons that satisfy a given longevity criterion by the total number of neurons that could have been tracked for that long (range bound). For example, a neuron first appearing on day 14 (range bound = 2) is only included in the calculation for P(tracked for 1 day) and P(tracked for 2 days) but is not counted in the numerator for P(tracked for 10 days). Fig. 6f tracked individual pairs of connections rather than individual units, but the calculation and presentation remain similar, with the exception that the range bound of a appear were decided based on the range between the first and last co-appearing day of the two units forming the pair. Fig. 6g represents the expected fraction of non-trackable

(lost) pairs after N future days, normalized by the elapsed days N. Fig. 6g (portion lost) is inherently related to Fig. 6f (portion tracked), it's defined as (1-portion tracked)/elapsed days. For the best comparison with past literature, our tracked portion analysis is similar to those in Chung et al., 2019.

**Motion videography**

Under IR illumination from Infrared LEDs, two Teledyne FLIR BFS cameras with their IR filters removed were used to capture videos from awake head-fixed mouse at 30 frames per second. One camera targeted the left pupil while the other targeted the forelimb and face of the mice. The cameras were synchronized with simultaneous electrical recording and visual stimulation through TTL signals generated from their general-purpose input and output (GPIO) ports.

**Automatic extraction of mouse behaviors.**

The behavioral variables were extracted from the videos through DeepLabCut[21] and YOLO[22]. Specifically, the DeepLabCut was used to automatically extract the positions of the front limbs as two-point sources and two points on the two edges of the pupil. The center position and size of the pupil were approximately derived from YOLO labeled rectangular bounding boxes.
